## Supplementary File 1 for "The Flexiscope: a Low Cost, Flexible, Convertible, and Modular Microscope with Automated Scanning and Micromanipulation": ReadMe.txt

This folder contains the four matlab files to control the Automated Piezo Scanning Stage. Guidelines to assemble and run these files can be found in 'UserGuidelines_AutomatedAcquisition_PiezoStage_Matlab.pdf'

Matlab files:

StageAutomation_Cam_Piezo_Variable_SetUp.m 

StageAutomation_GUI_1.fig

StageAutomation_GUI_1.m

StageAutomation_AcquisitionMotionCommands.m
