## Supplementary File 1 for "The Flexiscope: a Low Cost, Flexible, Convertible, and Modular Microscope with Automated Scanning and Micromanipulation": StageAutomation_AcquisitionMotionCommands.m

```
%imgnum is an important variable for image numenclature and also used to determine
%whether Z-Stack is achieved by moving Stage up or down. Even imgnum (2,4,6,8 etc) moves
%stage down while uneven (1,3,5,7 etc) imgnum moves stage up.
imgnum =ImgNumStart; %Start at ImgNumStart defined by user in GUI. imgnum increases by 1 with each new FOV


%%%Home all channels to 0 when tissue is positioned at the imaging starting point
h.SetPositionSteps(0,0);
h.SetPositionSteps(1,0);
h.SetPositionSteps(2,0);

for v=1:FOV_Y %Y-motion FOV loop
    
    if strcmpi(lens, '4X'); %if user defined lens as 4X in GUI
        FOVrightpiezo = -90000; 
        FOVleftpiezo = 90000;
         
    elseif strcmpi(lens, '20X'); %if user defined lens as 20X in GUI
        FOVrightpiezo = -18000;
        FOVleftpiezo = 18000;
    end

for hz=1:FOV_X %X-motion FOV loop


 for z=1:Z %Z stack acquisition loop
RGB = getsnapshot(vid); %get image from live video stream 
even = mod(imgnum,2);   %determine if imgnum is odd or even
%Z stack acquisition with most in focus values obtained
if even==1; %if imgnum is odd
    h.MoveJog(0, 1); %move Z-axis anticlockwise: move stage down
     pause(ZWait); %wait in seconds
     Image = rgb2gray(RGB); %RGB image to gray scale image
     Rx = imfilter(double(Image), Gx, 'conv', 'replicate');% Auto-focus algorithm: see Geusebroek et al, 2000 
     Ry = imfilter(double(Image), Gy, 'conv', 'replicate');% Auto-focus algorithm: see Geusebroek et al, 2000 
     FM = Rx.^2+Ry.^2;% Auto-focus algorithm: see Geusebroek et al, 2000 
     FMmaster(z) = mean2(FM);%This saves all the focus values for each image in a Z-stack as a matrix
%Z stack acquisition without most in focus values obtained    
elseif even ==0; %if imgnum is even
     h.MoveJog(0, 2); %%move Z-axis clockwise: move stage up
     pause(ZWait); %wait in seconds
end
 N4 = num2str(imgnum); %imgnum as string
 N7 = strcat(N1, N5, N2, lens, N5, N4, N6); %add all image name prefixes together
 fname = sprintf(N7, z);%add Z-stack number to the end of image name
 imwrite(RGB, [fname, '.tif'], 'Compression', 'none'); %save image with appropriate image name as a TIFF  
 end

%Correcting Z-axis to ensure region of interest remains in focus
     if even==0; %Don't perform this correction if stage has just moved up i.e. imgnum is even    
     elseif even ==1; %Only performs this correction if stage has just moved down i.e imgnum is odd
         [M, F1] = max(FMmaster); %find the max value in FMmaster and where it is located in the matrix. This tells us the Z-stack which was most in focus. We want to keep The most in focus region in the middle of the Z-stack.
        F2 = ceil(((Z/2)-F1)*Zadj)% Z/2 is the middle of the Z-Stack. F1 is the most in focus image from the last Z-stack. For example if Z is 20 we want the most in focus Z-image to be 10. Zadj corrects for discrepency is step size when stage moving up compared to down.
         if F2<=0; % if Z-correction value is less than 0 then no correction is required
             disp('No Z focus correction required');
         else 
             for f=1:F2; %Z-drift correction loop
                 h.MoveJog(0, 2);%%move Z-axis clockwise: move stage up
                 pause(ZWaitcorrect);
             end
         end
     end


imgnum = imgnum +1 %imgnum increases by 1 with each new FOV

 if hz==FOV_X; %If X-motion loop has reached final FOV_X
    disp('end of row');
 elseif hz<=FOV_X %If X-Motion loop hasn�t reached final FOV_X
    rightorleft = mod(v,2); %Is Y-motion loop value odd or even?
    if rightorleft==b ; %user clicked right on GUI and Y-motion loop value is odd or user clicked left and Y-motion loop value is even
    h.MoveAbsoluteStepsEx(2, FOVrightpiezo, false); %Move X-axis right to new FOV (this command moves from the current position (number on piezo GUI) to the defined position (FOVrightpiezo))
    FOVrightpiezo = FOVrightpiezo + FOVrightpiezoadd; %X-motion counter to the right(FOVrightpiezo increases by FOVrightpiezoadd with each X-motion loop)
    pause(FOVrightleftpiezoWait);%wait until move finished
elseif rightorleft==a ; %user clicked left on gui and Y-motion loop value is odd or user clicked right and Y-motion loop value is even
    h.MoveAbsoluteStepsEx(2, FOVleftpiezo, false);%Move X-axis left to new FOV (this command moves from the current position (number on piezo GUI) to the defined position (FOVleftpiezo))
    FOVleftpiezo = FOVleftpiezo + FOVleftpiezoadd; %X-motion counter to the left(FOVleftpiezo increases by FOVleftpiezoadd with each X-motion loop)
    pause(FOVrightleftpiezoWait); %wait until move finished
end

 end 
 
end


if v==FOV_Y; %If Y-motion loop has reached final FOV_Y
elseif v<=FOV_Y; %If Y-Motion loop hasn�t reached final FOV_Y
if Down == 1; %If user clicked down box on GUI 
h.MoveAbsoluteStepsEx(1, FOVdownpiezo, false);%Move Y-axis down to new FOV (this command moves from the current position (number on piezo GUI) to the defined position (FOVdownpiezo))
FOVdownpiezo = FOVdownpiezo + FOVdownpiezoadd; %Y-motion counter down(FOVdownpiezo increases by FOVdownpiezoadd with each Y-motion loop)
pause(FOVdownuppiezoWait);%wait until move finished
elseif Up == 1; %If user clicked up box on GUI 
h.MoveAbsoluteStepsEx(1, FOVuppiezo, false); %Move Y-axis up to new FOV (this command moves from the current position (number on piezo GUI) to the defined position (FOVuppiezo))
FOVuppiezo = FOVuppiezo + FOVuppiezoadd; %Y-motion counter up(FOVuppiezo increases by FOVuppiezoadd with each Y-motion loop)
pause(FOVdownuppiezoWait);%wait until move finished
end
h.SetPositionSteps(2,0);%zero X-axis on Piezo GUI
    end 
end

 
```
