## Supplementary File 1 for "The Flexiscope: a Low Cost, Flexible, Convertible, and Modular Microscope with Automated Scanning and Micromanipulation": StageAutomation_Cam_Piezo_Variable_SetUp.m

```
%%%Set up Camera
vid = videoinput('pointgrey', 1, 'F7_RGB_1384x1032_Mode7');
src = getselectedsource(vid);
src.FrameRateMode = 'Off';
src.ExposureMode = 'Off';
src.SaturationMode = 'Off';
src.SharpnessMode = 'Off';
src.WhiteBalanceRBMode = 'Off';
src.Exposure = 0.11984;
src.GammaMode = 'Off';
src.HueMode = 'Off';
src.TriggerDelayMode = 'Off';
src.WhiteBalanceRBMode = 'Off';
src.Shutter= S; %shutter/exposure setting in milliseconds
preview(vid); %start live video

pause(10)  %wait 10 seconds

%Set up Piezo APT Controller in MATLAB figure
fpos    = get(0,'DefaultFigurePosition'); % figure default position
fpos(3) = 650; % figure window size; Width
fpos(4) = 450; % figure window size; Height
f = figure('Position', fpos,...
           'Menu','None',...
           'Name','APT GUI');
h = actxcontrol('APTPZMOTOR.APTPZMotorCtrl.1',[20 20 600 400 ], f); %Change this for your system
%h = actxcontrol('progid', position, fig_handle)
h.StartCtrl; % Start Control
SN = 65863980; % Set the Serial Number of your device
set(h,'HWSerialNum', SN); % Set up your controller with above information 
h.Identify; %flash LED on T-cube

pause(10) %wait 10 seconds

N5 = '_'; % image name prefix
N6 = '_%02d'; %Z stack loop value used to determine Z-stack number for image name prefix

%Auto-focus variables: see Geusebroek et al, 2000 for more details
WSize = 15; % Size of local window
N = floor(WSize/2);
sig = N/2.5;
[x,y] = meshgrid(-N:N, -N:N);
G = exp(-(x.^2+y.^2)/(2*sig^2))/(2*pi*sig);
Gx = -x.*G/(sig^2);
Gx = Gx/sum(abs(Gx(:)));
Gy = -y.*G/(sig^2);
Gy = Gy/sum(abs(Gy(:)));


%Do you want scanning to begin to the right or to the left? Based on GUI
%click box results
if Right == 1; %Right box was clicked
    a = 0;
    b = 1;
elseif Left == 1; %left box was clicked
     a = 1;
     b = 0;
end 
    
%%%Lens variables based lens drop down menu choice in GUI
if strcmpi(lens, '4X') %user clicked 4X
    ZStepSize = 1000; %arbitrary value given to the Z-piezo to move a specific distance. This variable will be different depending on the magnification chosen. 
    FOVrightpiezo = -90000; %Arbitrary value sent to the X-Piezo to travel an appropriate distance to reveal a new ‘field of view’ (FOV) to the right.
    FOVrightpiezoadd = -90000; %This variables adds this value to itself with each X-piezo loop to continue scanning to the right for the defined number of ‘FOVs’
    FOVleftpiezo = 90000; %Arbitrary value sent to the X-Piezo to travel an appropriate distance to reveal a new FOV to the left
    FOVleftpiezoadd = 90000; %This variables adds this value to itself with each X-piezo loop to continue scanning to the left for the defined number of X-axis ‘FOVs’
    FOVdownpiezo = -90000; %Arbitrary value sent to the Y-Piezo to travel an appropriate distance to reveal a new FOV down.
    FOVdownpiezoadd = -90000; %This variables adds this value to itself with each Y-piezo loop to continue scanning down for the defined number of Y-axis ‘FOVs’ 
    FOVuppiezo = 90000; %Arbitrary value sent to the Y-Piezo to travel an appropriate distance to reveal a new FOV up.
    FOVuppiezoadd = 90000; %This variables adds this value to itself with each Y-piezo loop to continue scanning up for the defined number of Y-axis ‘FOVs’ 
    Zadj = 1.5; %Correction factor to take into account the discrepancy in distance travelled when the Z-piezo is moving up or down.
    ZWait = (ceil(S/1000)) +1;  %Time to wait in seconds during Z-stack acquisition. Shutter in milliseconds divided by 1000 plus 1 (one added incase shutter is less than a second, ceil function rounds the number up)
    ZWaitcorrect = 3; %Time to wait in seconds when Z-correction is occurring
    FOVrightleftpiezoWait = 290;  %Time to wait in seconds when X-Piezo moving
    FOVdownuppiezoWait = 225; %Time to wait in seconds when Y-piezo moving

elseif strcmpi(lens, '20X') %user clicked 20X
    ZStepSize = 250;
    FOVrightpiezo = -18000;
    FOVleftpiezo = 18000;
    FOVrightpiezoadd = -18000;
    FOVleftpiezoadd = 18000;
    FOVdownpiezo = -18000;
    FOVdownpiezoadd = -18000;
    FOVuppiezo = 18000;
    FOVuppiezoadd = 18000;
    Zadj = 1.6; 
    ZWait = (ceil(S/1000)) +1; 
    ZWaitcorrect = 1;
    FOVrightleftpiezoWait = 50; 
    FOVdownuppiezoWait = 45;
end


h.SetJogMode(0, 2); %set jog mode for channel 1 (Z-axis)
h.SetJogStepSize(0,ZStepSize); %set jog step size for channel 1 (Z-axis). Depends on magnification
```
