## Supplementary File 1 for "The Flexiscope: a Low Cost, Flexible, Convertible, and Modular Microscope with Automated Scanning and Micromanipulation": StageAutomation_GUI_1.m

```
function varargout = StageAutomation_GUI_1(varargin)
% STAGEAUTOMATION_GUI_1 MATLAB code for StageAutomation_GUI_1.fig
%      STAGEAUTOMATION_GUI_1, by itself, creates a new STAGEAUTOMATION_GUI_1 or raises the existing
%      singleton*.
%
%      H = STAGEAUTOMATION_GUI_1 returns the handle to a new STAGEAUTOMATION_GUI_1 or the handle to
%      the existing singleton*.
%
%      STAGEAUTOMATION_GUI_1('CALLBACK',hObject,eventData,handles,...) calls the local
%      function named CALLBACK in STAGEAUTOMATION_GUI_1.M with the given input arguments.
%
%      STAGEAUTOMATION_GUI_1('Property','Value',...) creates a new STAGEAUTOMATION_GUI_1 or raises the
%      existing singleton*.  Starting from the left, property value pairs are
%      applied to the GUI before StageAutomation_GUI_1_OpeningFcn gets called.  An
%      unrecognized property name or invalid value makes property application
%      stop.  All inputs are passed to StageAutomation_GUI_1_OpeningFcn via varargin.
%
%      *See GUI Options on GUIDE's Tools menu.  Choose "GUI allows only one
%      instance to run (singleton)".
%
% See also: GUIDE, GUIDATA, GUIHANDLES

% Edit the above text to modify the response to help StageAutomation_GUI_1

% Last Modified by GUIDE v2.5 05-Dec-2017 16:07:16

% Begin initialization code - DO NOT EDIT
gui_Singleton = 1;
gui_State = struct('gui_Name',       mfilename, ...
                   'gui_Singleton',  gui_Singleton, ...
                   'gui_OpeningFcn', @StageAutomation_GUI_1_OpeningFcn, ...
                   'gui_OutputFcn',  @StageAutomation_GUI_1_OutputFcn, ...
                   'gui_LayoutFcn',  [] , ...
                   'gui_Callback',   []);
if nargin && ischar(varargin{1})
    gui_State.gui_Callback = str2func(varargin{1});
end

if nargout
    [varargout{1:nargout}] = gui_mainfcn(gui_State, varargin{:});
else
    gui_mainfcn(gui_State, varargin{:});
end
% End initialization code - DO NOT EDIT


% --- Executes just before StageAutomation_GUI_1 is made visible.
function StageAutomation_GUI_1_OpeningFcn(hObject, eventdata, handles, varargin)
% This function has no output args, see OutputFcn.
% hObject    handle to figure
% eventdata  reserved - to be defined in a future version of MATLAB
% handles    structure with handles and user data (see GUIDATA)
% varargin   command line arguments to StageAutomation_GUI_1 (see VARARGIN)

% Choose default command line output for StageAutomation_GUI_1
handles.output = hObject;

% Update handles structure
guidata(hObject, handles);

% UIWAIT makes StageAutomation_GUI_1 wait for user response (see UIRESUME)
% uiwait(handles.figure1);


% --- Outputs from this function are returned to the command line.
function varargout = StageAutomation_GUI_1_OutputFcn(hObject, eventdata, handles) 
% varargout  cell array for returning output args (see VARARGOUT);
% hObject    handle to figure
% eventdata  reserved - to be defined in a future version of MATLAB
% handles    structure with handles and user data (see GUIDATA)

% Get default command line output from handles structure
varargout{1} = handles.output;


% --- Executes on selection change in lens.
function lens_Callback(hObject, eventdata, handles)
% hObject    handle to lens (see GCBO)
% eventdata  reserved - to be defined in a future version of MATLAB
% handles    structure with handles and user data (see GUIDATA)

% Hints: contents = cellstr(get(hObject,'String')) returns lens contents as cell array
%        contents{get(hObject,'Value')} returns selected item from lens


% --- Executes during object creation, after setting all properties.
function lens_CreateFcn(hObject, eventdata, handles)
% hObject    handle to lens (see GCBO)
% eventdata  reserved - to be defined in a future version of MATLAB
% handles    empty - handles not created until after all CreateFcns called

% Hint: popupmenu controls usually have a white background on Windows.
%       See ISPC and COMPUTER.
if ispc && isequal(get(hObject,'BackgroundColor'), get(0,'defaultUicontrolBackgroundColor'))
    set(hObject,'BackgroundColor','white');
end


function N1_Callback(hObject, eventdata, handles)
% hObject    handle to N1 (see GCBO)
% eventdata  reserved - to be defined in a future version of MATLAB
% handles    structure with handles and user data (see GUIDATA)

% Hints: get(hObject,'String') returns contents of N1 as text
%        str2double(get(hObject,'String')) returns contents of N1 as a double

% --- Executes during object creation, after setting all properties.
function N1_CreateFcn(hObject, eventdata, handles)
% hObject    handle to N1 (see GCBO)
% eventdata  reserved - to be defined in a future version of MATLAB
% handles    empty - handles not created until after all CreateFcns called

% Hint: edit controls usually have a white background on Windows.
%       See ISPC and COMPUTER.
if ispc && isequal(get(hObject,'BackgroundColor'), get(0,'defaultUicontrolBackgroundColor'))
    set(hObject,'BackgroundColor','white');
end


function N2_Callback(hObject, eventdata, handles)
% hObject    handle to N2 (see GCBO)
% eventdata  reserved - to be defined in a future version of MATLAB
% handles    structure with handles and user data (see GUIDATA)


% Hints: get(hObject,'String') returns contents of N2 as text
%        str2double(get(hObject,'String')) returns contents of N2 as a double

% --- Executes during object creation, after setting all properties.
function N2_CreateFcn(hObject, eventdata, handles)
% hObject    handle to N2 (see GCBO)
% eventdata  reserved - to be defined in a future version of MATLAB
% handles    empty - handles not created until after all CreateFcns called

% Hint: edit controls usually have a white background on Windows.
%       See ISPC and COMPUTER.
if ispc && isequal(get(hObject,'BackgroundColor'), get(0,'defaultUicontrolBackgroundColor'))
    set(hObject,'BackgroundColor','white');
end


function N3_Callback(hObject, eventdata, handles)
% hObject    handle to N3 (see GCBO)
% eventdata  reserved - to be defined in a future version of MATLAB
% handles    structure with handles and user data (see GUIDATA)

% Hints: get(hObject,'String') returns contents of N3 as text
%        str2double(get(hObject,'String')) returns contents of N3 as a double


% --- Executes during object creation, after setting all properties.
function N3_CreateFcn(hObject, eventdata, handles)
% hObject    handle to N3 (see GCBO)
% eventdata  reserved - to be defined in a future version of MATLAB
% handles    empty - handles not created until after all CreateFcns called

% Hint: edit controls usually have a white background on Windows.
%       See ISPC and COMPUTER.
if ispc && isequal(get(hObject,'BackgroundColor'), get(0,'defaultUicontrolBackgroundColor'))
    set(hObject,'BackgroundColor','white');
end


function S_Callback(hObject, eventdata, handles)
% hObject    handle to S (see GCBO)
% eventdata  reserved - to be defined in a future version of MATLAB
% handles    structure with handles and user data (see GUIDATA)

% Hints: get(hObject,'String') returns contents of S as text
%        str2double(get(hObject,'String')) returns contents of S as a double


% --- Executes during object creation, after setting all properties.
function S_CreateFcn(hObject, eventdata, handles)
% hObject    handle to S (see GCBO)
% eventdata  reserved - to be defined in a future version of MATLAB
% handles    empty - handles not created until after all CreateFcns called

% Hint: edit controls usually have a white background on Windows.
%       See ISPC and COMPUTER.
if ispc && isequal(get(hObject,'BackgroundColor'), get(0,'defaultUicontrolBackgroundColor'))
    set(hObject,'BackgroundColor','white');
end


function Z_Callback(hObject, eventdata, handles)
% hObject    handle to Z (see GCBO)
% eventdata  reserved - to be defined in a future version of MATLAB
% handles    structure with handles and user data (see GUIDATA)


% Hints: get(hObject,'String') returns contents of Z as text
%        str2double(get(hObject,'String')) returns contents of Z as a double

% --- Executes during object creation, after setting all properties.
function Z_CreateFcn(hObject, eventdata, handles)
% hObject    handle to Z (see GCBO)
% eventdata  reserved - to be defined in a future version of MATLAB
% handles    empty - handles not created until after all CreateFcns called

% Hint: edit controls usually have a white background on Windows.
%       See ISPC and COMPUTER.
if ispc && isequal(get(hObject,'BackgroundColor'), get(0,'defaultUicontrolBackgroundColor'))
    set(hObject,'BackgroundColor','white');
end


function FOV_X_Callback(hObject, eventdata, handles)
% hObject    handle to FOV_X (see GCBO)
% eventdata  reserved - to be defined in a future version of MATLAB
% handles    structure with handles and user data (see GUIDATA)

% Hints: get(hObject,'String') returns contents of FOV_X as text
%        str2double(get(hObject,'String')) returns contents of FOV_X as a double


% --- Executes during object creation, after setting all properties.
function FOV_X_CreateFcn(hObject, eventdata, handles)
% hObject    handle to FOV_X (see GCBO)
% eventdata  reserved - to be defined in a future version of MATLAB
% handles    empty - handles not created until after all CreateFcns called

% Hint: edit controls usually have a white background on Windows.
%       See ISPC and COMPUTER.
if ispc && isequal(get(hObject,'BackgroundColor'), get(0,'defaultUicontrolBackgroundColor'))
    set(hObject,'BackgroundColor','white');
end


function FOV_Y_Callback(hObject, eventdata, handles)
% hObject    handle to FOV_Y (see GCBO)
% eventdata  reserved - to be defined in a future version of MATLAB
% handles    structure with handles and user data (see GUIDATA)

% Hints: get(hObject,'String') returns contents of FOV_Y as text
%        str2double(get(hObject,'String')) returns contents of FOV_Y as a double

% --- Executes during object creation, after setting all properties.
function FOV_Y_CreateFcn(hObject, eventdata, handles)
% hObject    handle to FOV_Y (see GCBO)
% eventdata  reserved - to be defined in a future version of MATLAB
% handles    empty - handles not created until after all CreateFcns called

% Hint: edit controls usually have a white background on Windows.
%       See ISPC and COMPUTER.
if ispc && isequal(get(hObject,'BackgroundColor'), get(0,'defaultUicontrolBackgroundColor'))
    set(hObject,'BackgroundColor','white');
end


% --- Executes on button press in GO.
function GO_Callback(hObject, eventdata, handles)
% hObject    handle to GO (see GCBO)
% eventdata  reserved - to be defined in a future version of MATLAB
% handles    structure with handles and user data (see GUIDATA)
N1=get(handles.N1,'String'); %file name - Prefix 1
assignin('base', 'N1',N1); %send to workspace

N2=get(handles.N2,'String'); %file name - Prefix 2
assignin('base', 'N2',N2);

lens1 = get(handles.lens,'String'); %Lens used
lens2  = get(handles.lens,'Value');
lens = (lens1{lens2});
assignin('base', 'lens',lens);

Z= str2num(get(handles.Z,'String')); %number of Z stacks
assignin('base', 'Z',Z);

S=str2num(get(handles.S,'String')); %exposure time in ms
assignin('base', 'S',S);

FOV_X = str2num(get(handles.FOV_X,'String')); %How many FOVs in the X direction
assignin('base', 'FOV_X',FOV_X);

FOV_Y = str2num(get(handles.FOV_Y,'String')); %How many FOVs in the Y direction
assignin('base', 'FOV_Y',FOV_Y)

Right = get(handles.Right,'Value'); %Begin moving right (camera perspective)
assignin('base', 'Right',Right);

Left = get(handles.Left,'Value'); %Begin moving left (camera perspective)
assignin('base', 'Left',Left);

Down = get(handles.Down,'Value'); %Begin moving down (camera perspective)
assignin('base', 'Down',Down);

Up = get(handles.Up,'Value'); %Begin moving up (camera perspective)
assignin('base', 'Up',Up);

ImgNumStart = str2num(get(handles.ImgNumStart,'String'));
assignin('base', 'ImgNumStart',ImgNumStart);

close('StageAutomation_GUI_1')

% --- Executes on key press with focus on GO and none of its controls.
function GO_KeyPressFcn(hObject, eventdata, handles)
% hObject    handle to GO (see GCBO)
% eventdata  structure with the following fields (see MATLAB.UI.CONTROL.UICONTROL)
%	Key: name of the key that was pressed, in lower case
%	Character: character interpretation of the key(s) that was pressed
%	Modifier: name(s) of the modifier key(s) (i.e., control, shift) pressed
% handles    structure with handles and user data (see GUIDATA)


% --- Executes on button press in Right.
function Right_Callback(hObject, eventdata, handles)
% hObject    handle to Right (see GCBO)
% eventdata  reserved - to be defined in a future version of MATLAB
% handles    structure with handles and user data (see GUIDATA)

% Hint: get(hObject,'Value') returns toggle state of Right


% --- Executes on button press in Left.
function Left_Callback(hObject, eventdata, handles)
% hObject    handle to Left (see GCBO)
% eventdata  reserved - to be defined in a future version of MATLAB
% handles    structure with handles and user data (see GUIDATA)

% Hint: get(hObject,'Value') returns toggle state of Left


% --- Executes on button press in Down.
function Down_Callback(hObject, eventdata, handles)
% hObject    handle to Down (see GCBO)
% eventdata  reserved - to be defined in a future version of MATLAB
% handles    structure with handles and user data (see GUIDATA)

% Hint: get(hObject,'Value') returns toggle state of Down


% --- Executes on button press in Up.
function Up_Callback(hObject, eventdata, handles)
% hObject    handle to Up (see GCBO)
% eventdata  reserved - to be defined in a future version of MATLAB
% handles    structure with handles and user data (see GUIDATA)

% Hint: get(hObject,'Value') returns toggle state of Up


function ImgNumStart_Callback(hObject, eventdata, handles)
% hObject    handle to ImgNumStart (see GCBO)
% eventdata  reserved - to be defined in a future version of MATLAB
% handles    structure with handles and user data (see GUIDATA)

% Hints: get(hObject,'String') returns contents of ImgNumStart as text
%        str2double(get(hObject,'String')) returns contents of ImgNumStart as a double


% --- Executes during object creation, after setting all properties.
function ImgNumStart_CreateFcn(hObject, eventdata, handles)
% hObject    handle to ImgNumStart (see GCBO)
% eventdata  reserved - to be defined in a future version of MATLAB
% handles    empty - handles not created until after all CreateFcns called

% Hint: edit controls usually have a white background on Windows.
%       See ISPC and COMPUTER.
if ispc && isequal(get(hObject,'BackgroundColor'), get(0,'defaultUicontrolBackgroundColor'))
    set(hObject,'BackgroundColor','white');
end
```
