## Supplementary File 1 for "The Flexiscope: a Low Cost, Flexible, Convertible, and Modular Microscope with Automated Scanning and Micromanipulation": UserGuidelines_AutomatedAcquisition_PiezoStage_Matlab.pdf

### User Guidelines: Automated Acquisition and Piezoelectric Stage Scanning in Matlab

**March 2019**

Step1 and 2 should be performed once when you are establishing your system.

Steps 3-5 are performed each time you want to set up a tissue sample for automated scanning.

**Step1: Matlab ActiveX Control of Piezoelectric Actuators (MC2) using T-cube (MC3) and APT Software**

A. Type actxcontrolselect into Matlab command window to determine the ActiveX control ProgID of your T-cube (Figure 1) using the guided user interface (GUI).
B. Get the Serial Number of your T-cube (found physically on the device).
C. Alter 'Set up Piezo APT Controller in MATLAB figure' commands in 'StageAutomation_Cam_Piezo_Variable_SetUp' to suit your system.

**Step2: Determine Optimal Camera Settings within Matlab**

A. The camera used in our system for fluorescent imaging is CO23. The 'Camera Settings' commands found in 'StageAutomation_Cam_Piezo_Variable_SetUp' are therefore optimal for our system. If you are using a different camera you will need to open your camera in the image acquisition toolbox in Matlab (make sure you have installed appropriate add on packages for your camera:

https://uk.mathworks.com/help/imaq/installing-the-support-packages-for-image-acquisition-toolbox-adaptors.html).

B. Alter the 'acquisition parameters' until you achieved the desired settings. The appropriate commands for these specific settings can be seen in 'Session Log'.
C. Input the appropriate commands into the 'Camera Settings' commands in 'StageAutomation_Cam_Piezo_Variable_SetUp'

**Step3: Time to do your first scan!**

A. Set up camera settings directly in the FlyCap software as seen in figure 2 and determine optimal shutter (exposure time in ms).
B. Find your starting position within or at the edge of the tissue sample.
C. Ensure sufficient travel in the X and Y-axis in the direction you want the stage to scan (for example: scanning right and down requires the X- and Y- axis to be rotated fully anticlockwise until range of travel has been reached to allow maximum travel clockwise which is the desired direction of scanning)
D. Adjust Z axis:

- Starting imgnum odd: Above most in focus region i.e. rotating Z actuator anticlockwise (stage down) so that the image comes in and out of focus
- Starting imgnum is even: Below most in focus region i.e. rotating Z actuator clockwise (stage up) so that the image comes in and out of focus

**Step4: Matlab – User settings GUI**

A. Start Matlab
B. Navigate 'current folder' to folder where you would like to save your images
C. Ensure the following files are in this folder:
    A. StageAutomation_GUI_1.fig
    B. StageAutomation_GUI_1.m
    C. StageAutomation_AcquisitionMotionCommands.m
    D. StageAutomation_Cam_Piezo_Variable_SetUp.m
    E. Type into Run('StageAutomation_GUI_1.m') into command window. The Stage automation GUI should begin (Figure 3)

D. Complete all edit boxes with desired settings
E. Click Begin Acquisition
F. Make sure you're happy with settings (variables now in workspace)

**Step5: Matlab - Live stream camera display and piezo GUI set up**

A. Type Run('StageAutomation_Cam_Piezo_Variable_SetUp.m') into command window. The video preview window and the Piezo APT GUI should begin along with additional variables in the workspace (Figure 4).
B. Ensure you are happy with camera view and variables in workspace (tip: if you're not happy with any variable values simply restart from step 4 or directly alter them in workspace. For example, to change exposure time to four seconds type: S=4000'; src.Shutter= S; into command window and the live video will update itself automatically)
C. When you are ready to start scanning type Run('StageAutomation_AcquisitionMotionCommands.m') into command window.

**Variables Explained:**

S: exposure time in milliseconds
N1: image name prefix 1
N2: image name prefix 2
N5: image name prefix
N6: Z stack loop value used to determine Z-stack number for image name prefix
Lens: lens magnification defined in GUI
XChoice: right or left X direction scanning to begin (this is from my camera's perspective, this may be opposite on your system)
YChoice: Y direction for scanning up or down (this is from my camera's perspective, this may be opposite on your system)
ZStepSize: arbitrary value given to the Z-piezo to move a specific distance. This variable will be different depending on the magnification chosen in the GUI.
FOVrightpiezo: Arbitrary value sent to the X-Piezo to travel an appropriate distance to reveal a new 'field of view' (FOV) to the right.

- some overlap to allow image stitching later

- Movement to the right is from my camera's perspective, this may be opposite on your system
- This variable will be different depending on the magnification defined in the GUI

FOVleftpiezo: Arbitrary value sent to the X-Piezo to travel an appropriate distance to reveal a new FOV to the left

- some overlap to allow image stitching later
- Movement to the left is from my camera's perspective, this may be opposite on your system
- This variable will be different depending on the magnification defined in the GUI

FOVrightpiezoadd: This variables adds this value to itself with each X-piezo loop to continue scanning to the right for the defined number of 'FOVs'

FOVleftpiezoadd: This variables adds this value to itself with each X-piezo loop to continue scanning to the left for the defined number of X-axis 'FOVs'

FOVdownpiezo: Arbitrary value sent to the Y-Piezo to travel an appropriate distance to reveal a new FOV down.

- some overlap to allow image stitching later
- Movement down is from my camera's perspective, this may be opposite on your system
- This variable will be different depending on the magnification defined in the GUI

FOVdownpiezoadd: This variables adds this value to itself with each Y-piezo loop to continue scanning down for the defined number of Y-axis 'FOVs'

FOVuppiezo: Arbitrary value sent to the Y-Piezo to travel an appropriate distance to reveal a new FOV up.

- some overlap to allow image stitching later
- Movement up is from my camera's perspective, this may be opposite on your system
- This variable will be different depending on the magnification defined in the GUI

FOVuppiezoadd: This variables adds this value to itself with each Y-piezo loop to continue scanning up for the defined number of Y-axis 'FOVs'

Z: Number of Z Stacks as defined by user in GUI

Zadj: Correction factor to take into account the discrepancy in distance travelled when the Z-piezo is moving up or down. This can be adjusted within 'StageAutomation_Cam_Piezo_Variable_SetUp' depending on your system.

ZWait = (ceil(S/1000)+1): Time to wait in seconds during Z-stack acquisition. Exposure time in milliseconds divided by 1000 plus 1 (one added in case shutter is less than a second, ceil function rounds the number up)

ZWaitcorrect: Time to wait in seconds when Z-drift correction loop is occurring

FOVrightleftpiezoWait: Time to wait in seconds when X-Piezo is moving right or left

FOVdownuppiezoWait: Time to wait in seconds when Y-piezo is moving up or down

Auto-focus variables: see (Geusebroek et al., 2000) for more details

```
WSize = 15; % Size of local window
N = floor(WSize/2);
sig = N/2.5;
[x,y] = meshgrid(-N:N, -N:N);
G = exp(-(x.^2+y.^2)/(2*sig^2))/(2*pi*sig);
Gx = -x.*G/(sig^2);
Gx = Gx/sum(abs(Gx(:)));
Gy = -y.*G/(sig^2);
Gy = Gy/sum(abs(Gy(:)));
```

ImgNumStart: Image name number prefix for first image. Defined in GUI

imgnum: imgnum is an important variable for image nomenclature and also used to determine whether Z-Stack is achieved by moving Stage up or down (Z-piezo). Even imgnum moves stage up while odd imgnum moves stage down.

Left: if this variables is equal to 1 X-motion scanning will begin to the left (it looks like the FOV is moving to the left from the camera's perspective - clockwise X-piezo actuation- stage physically moving right). If this variable is equal to 0, X-motion scanning will begin to the right. Defined in GUI with tick boxes. Movement left is from my camera's perspective, this may be opposite on your system

Right: if this variables is equal to 1 X-motion scanning will begin to the right (it looks like the FOV is moving to the right from the camera's perspective - anticlockwise X-piezo actuation-stage physically moving left). If this variable is equal to 0, X-motion scanning will begin to the left. Defined in GUI with tick boxes. Movement right is from my camera's perspective, this may be opposite on your system

Up: Do not confuse up in the Y-axis with up in the Z-axis. Up in the Y-axis is technically moving the stage forward (clockwise Y-piezo actuation) but from the camera perspective it looks like the FOV is moving up so this naming stuck. If this variables is equal to 1 Y-motion scanning will continue up. If this variable is equal to 0, Y-motion scanning will continue down. Defined in GUI with tick boxes. Movement up is from my camera's perspective, this may be opposite on your system.

Down: Do not confuse down in the Y-axis with down in the Z-axis. Down in the Y-axis is technically moving the stage backward (anticlockwise Y-piezo actuation) but from the camera perspective it looks like the FOV is moving down so this naming stuck. If this variables is equal to 1 Y-motion scanning will continue down. If this variable is equal to 0, Y-motion scanning will continue up. Defined in GUI with tick boxes. Movement down is from my camera's perspective, this may be opposite on your system

FOV_X: X-motion number of new FOVs aka number of times X-motion loop will repeat

FOV_Y: Y-motion number of new FOVs aka number of times Y-motion loop will repeat

**X/Y scanning and Z-stack acquisition explained:**

Figure 5 explains the workflow to acquire Z-stacks at regions of the sample in the X and Y direction. The commands used to control the specimen stage followed a logic in which the image sequence of the tissue sample is considered a two dimensional array. To acquire images of the whole tissue (or a region of interest) the stage must be moved in the X-axis a defined number of times and then moved in the Y-axis once, this cycle can then be repeated until the tissue (or region of interest) has been imaged in its entirety. Each movement in X or Y dimension reveals a new'field of view' (FOV) and a Z-stack is subsequently acquired. The motion control and image acquisition code is explained schematically in figure 6.

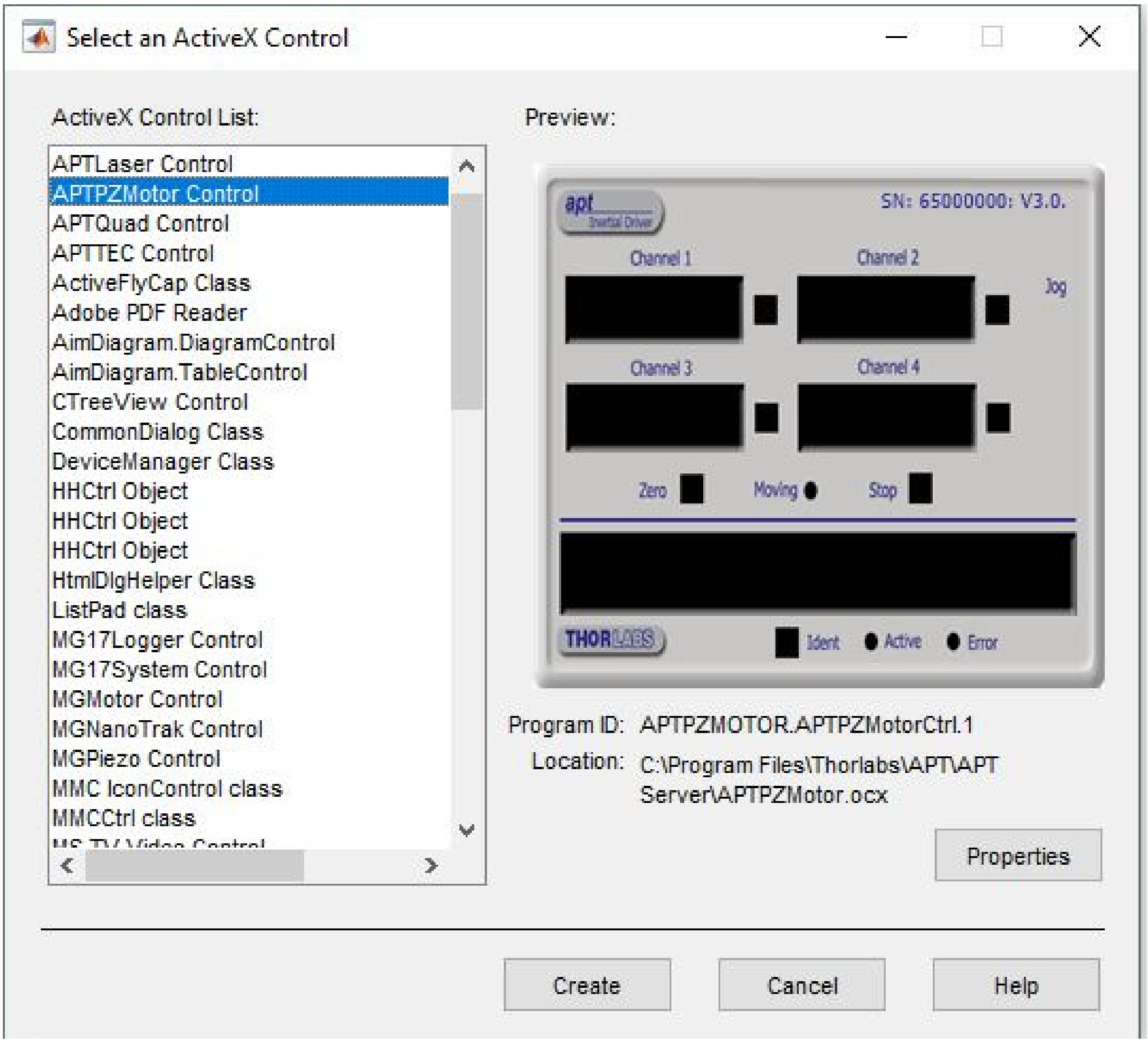


Figure1. ActiveX control select GUI to determine Program ID of your device.

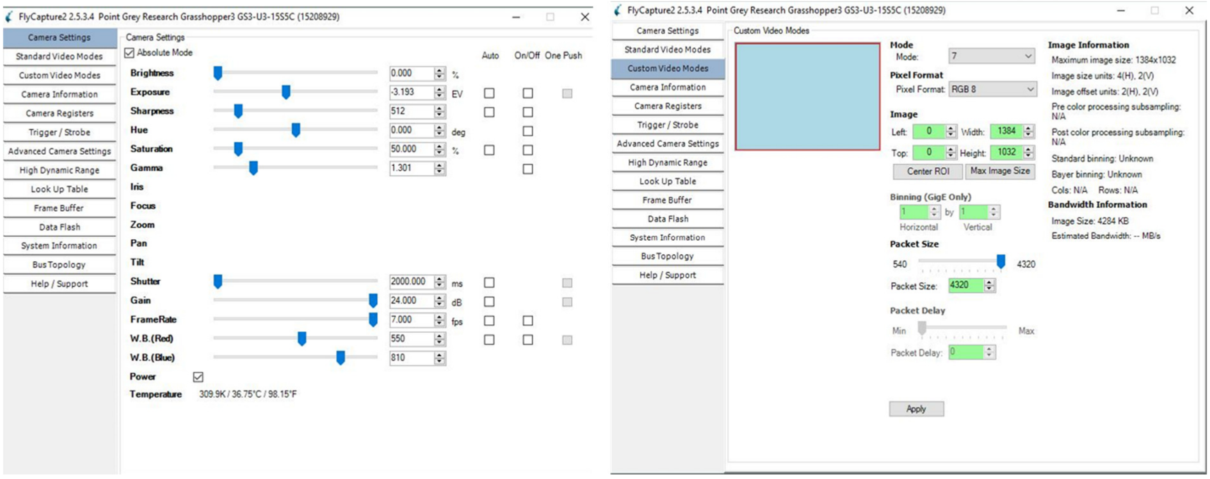


Figure2. FlyCap settings for CO23 for fluorescent imaging

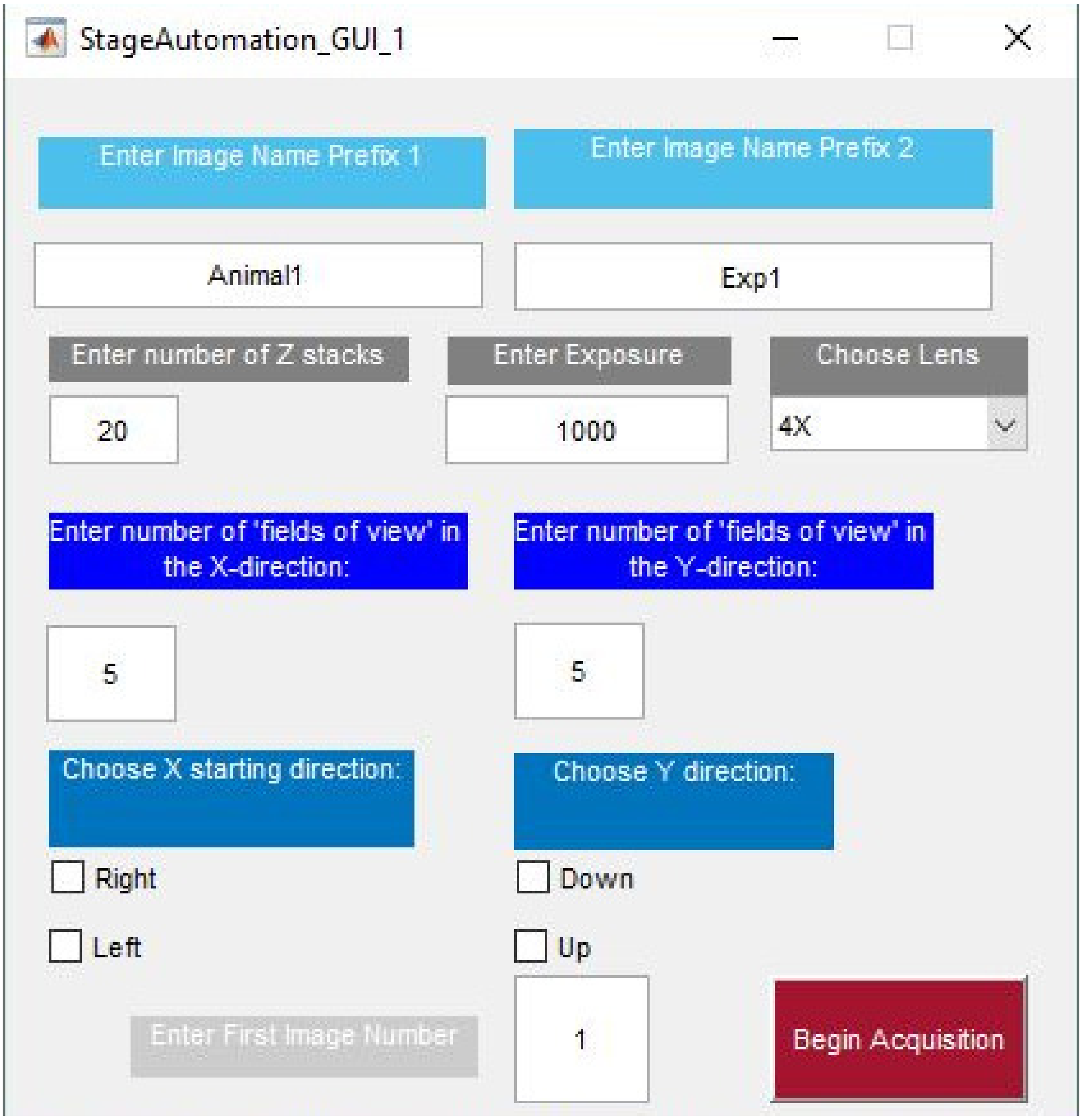


Figure 3. Automated piezoelectric stage motion control GUI in Matlab

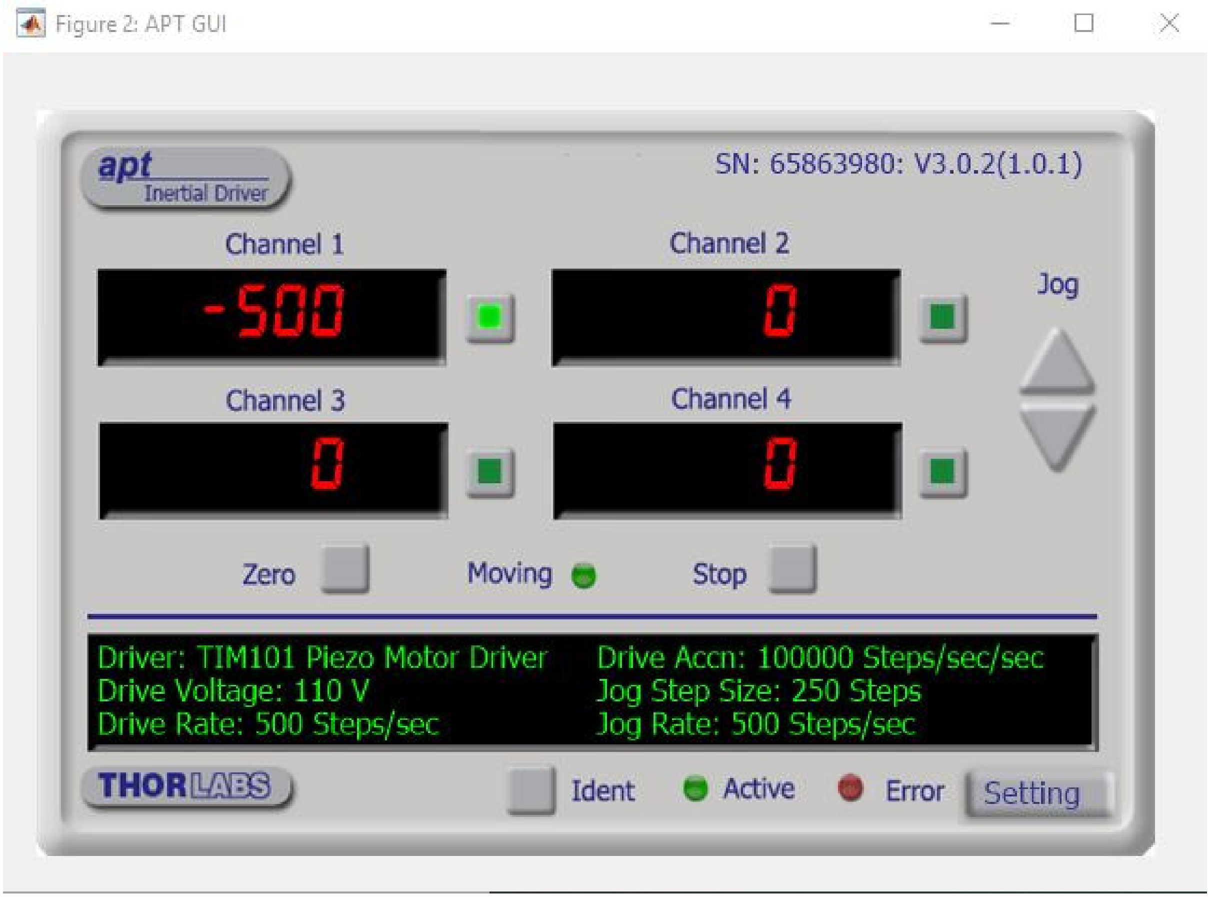


Figure4. APT GUI in Matlab figure window

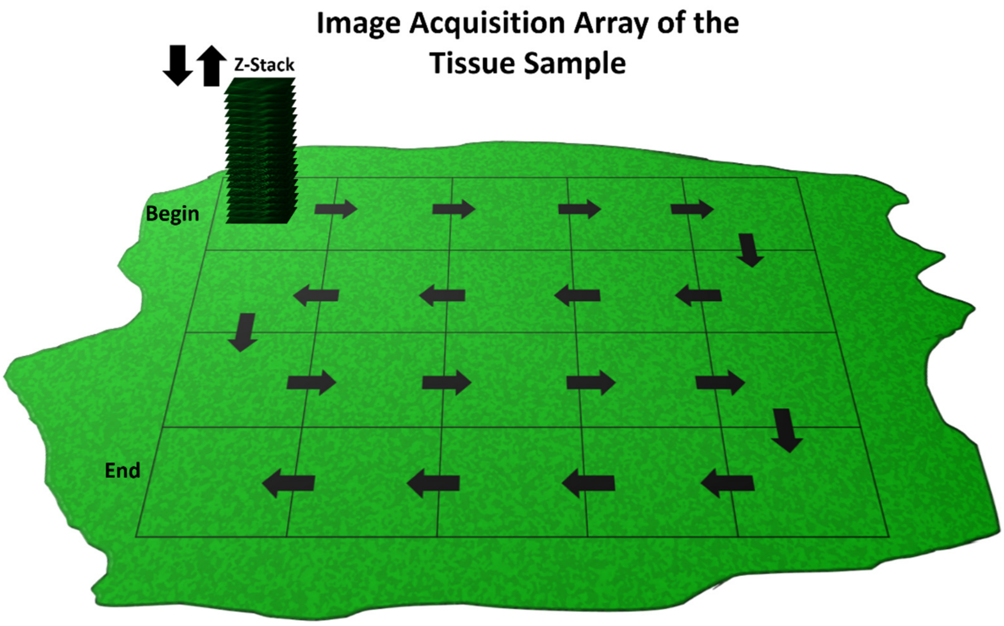

Figure5. Schematic representation of the automated stage motion control

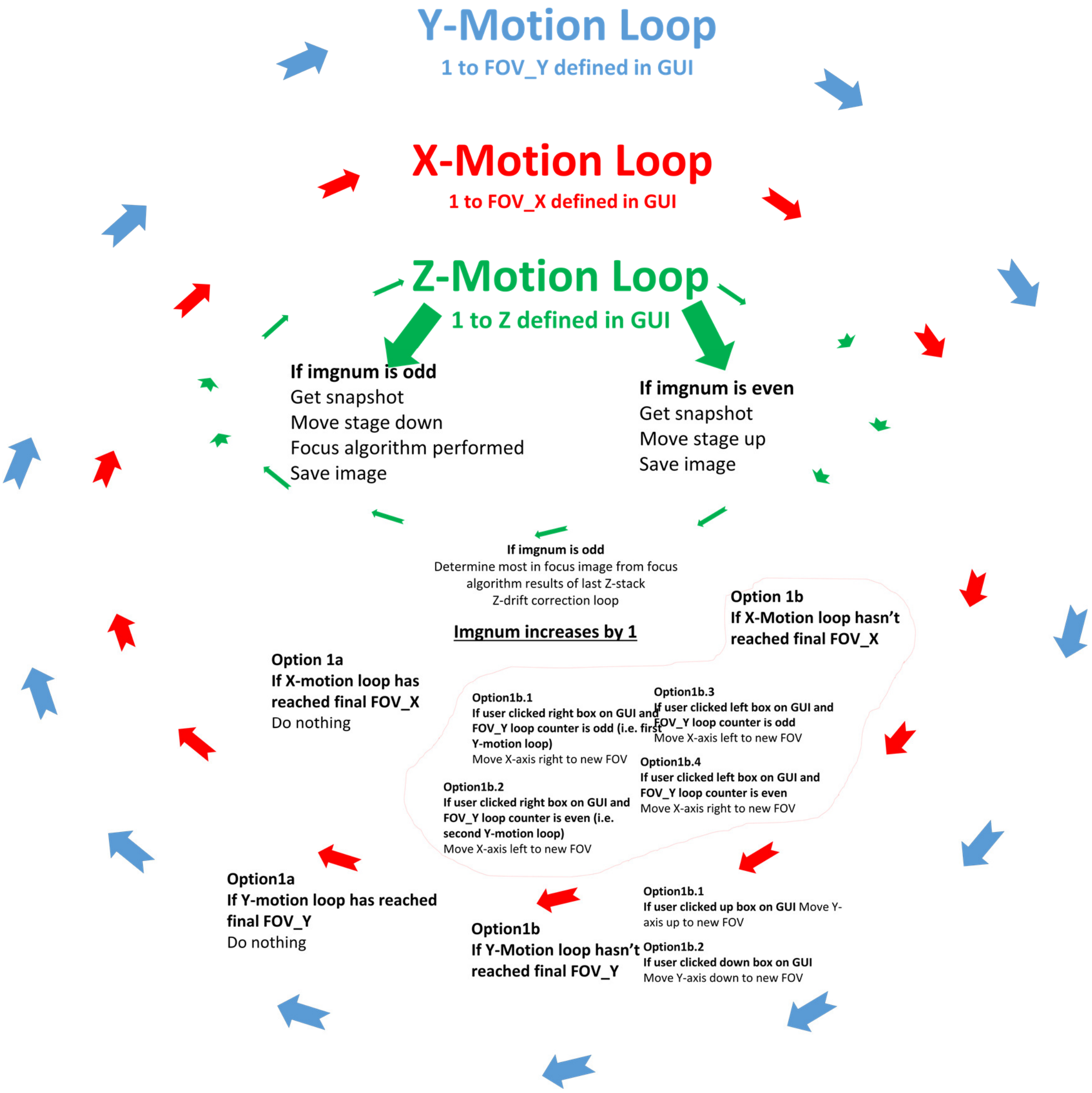


Figure6. Schematic representation of the automated piezoelectric motion control and image acquisition code.
