## Supplementary File 2 for "The Flexiscope: a Low Cost, Flexible, Convertible, and Modular Microscope with Automated Scanning and Micromanipulation": ReadMe.txt

This folder contains the four matlab files to control the Automated Stepper Scanning Stage. Guidelines to assemble and run these files can be found in 'UserGuidelines_AutomatedAcquisition_StepperStage_Matlab.pdf'

Matlab files:

StageAutomation_AcquisitionMotionCommands_Steppers.m

StageAutomation_Cam_Steppers_Variable_SetUp.m

StageAutomation_GUI_1.fig

StageAutomation_GUI_1.m
