## Supplementary File 2 for "The Flexiscope: a Low Cost, Flexible, Convertible, and Modular Microscope with Automated Scanning and Micromanipulation": StageAutomation_AcquisitionMotionCommands_Steppers.m

```
%imgnum is an important variable for image numenclature and also used to determine
%whether Z-Stack is achieved by moving Stage up or down. Even imgnum (2,4,6,8 etc) moves
%stage up while uneven (1,3,5,7 etc) imgnum moves stage down.
imgnum =ImgNumStart; %Start at ImgNumStart defined by user in GUI.  imgnum increases by 1 with each new FOV

for v=1:FOV_Y %Y-motion FOV loop
    
for hz=1:FOV_X %X-motion FOV loop
    
 for z=1:Z %Z stack acquisition loop
RGB = getsnapshot(vid); %get image from live video stream 
even = mod(imgnum,2);   %determine if imgnum is odd or even
%Z stack acquisition with most in focus values obtained
if even==1; %if imgnum is odd
     fprintf(gbrl,ZStageDown); %Gcode command: move Z-axis anticlockwise aka move stage down
     pause(ZWait); %wait in seconds
elseif even ==0; %if imgnum is even
     fprintf(gbrl,ZStageUp); %Gcode command: move Z-axis clockwise - move stage up
     pause(ZWait); %wait in seconds
end
     Image = rgb2gray(RGB); %RGB image to gray scale image
     Rx = imfilter(double(Image), Gx, 'conv', 'replicate');% Auto-focus algorithm: see Geusebroek et al, 2000 
     Ry = imfilter(double(Image), Gy, 'conv', 'replicate');% Auto-focus algorithm: see Geusebroek et al, 2000 
     FM = Rx.^2+Ry.^2;% Auto-focus algorithm: see Geusebroek et al, 2000 
     FMmaster(z) = mean2(FM);%This saves all the focus values for each image in a Z-stack as a matrix
    N4 = num2str(imgnum); %imgnum as string
    N7 = strcat(N1, N5, N2, lens, N5, N4, N6); %add all image name prefixes together
    fname = sprintf(N7, z);%add Z-stack number to the end of image name
    imwrite(RGB, [fname, '.tif'], 'Compression', 'none'); %save image with appropriate image name as a TIFF  
 end
 
 
MeanI = mean2(RGB) % get mean intensity of last image captured at this FOV
%Correcting Z-axis to ensure region of interest remains in focus
[M, F1] = max(FMmaster);%find the max value in FMmaster and where it is located in the matrix. This tells us the Z-stack which was most in focus. We want to keep The most in focus region in the middle of the Z-stack.
F2 = ceil(((Z/2))-F1)% Z/2 is the middle of the Z-Stack. F1 is the most in focus image from the last Z-stack. For example if Z is 20 we want the most in focus Z-image to be 10.

 if F2<=0 || MeanI<=2;% if Z-correction value is less than 0 or if mean of last image is less than 5 (no tissue in view) then no correction is required
             disp('No Z-axis focus correction required');
 else
      
     if even==1; %Only performs this correction if stage has just moved down i.e. imgnum is odd
             for f=1:F2; %Z-drift correction loop
                 fprintf(gbrl,ZStageUp);%%move Z-axis clockwise: move stage up
                 pause(ZWaitcorrect);
             end
     elseif even ==0; %Only performs this correction if stage has just moved up i.e. imgnum is even
             for f=1:F2; %Z-drift correction loop
                 fprintf(gbrl,ZStageDown);%%move Z-axis clockwise: move stage up
                 pause(ZWaitcorrect);
             end
     end
  end


imgnum = imgnum +1 %imgnum increases by 1 with each new FOV

 if hz==FOV_X; %If X-motion loop has reached final FOV_X
    disp('End of row');
 elseif hz<=FOV_X %If X-Motion loop hasn�t reached final FOV_X
    rightorleft = mod(v,2); %Is Y-motion loop value odd or even?
    if rightorleft==b ; %user clicked right on gui and Y-motion loop value is odd or user clicked left and Y-motion loop value is even
    fprintf(gbrl,XFOVright);  %Gcode command: Move X-axis right to new FOV
    disp('New Field of View to the RIGHT complete');
    pause(FOVrightleftWait);%wait until move finished
elseif rightorleft==a ; %user clicked left on gui and Y-motion loop value is odd or user clicked right and Y-motion loop value is even
    fprintf(gbrl,XFOVleft);%Gcode command: Move X-axis left to new FOV 
    disp('New Field of View to the LEFT complete');
    pause(FOVrightleftWait); %wait until move finished
end

 end 
 
end


if v==FOV_Y; %If Y-motion loop has reached final FOV_Y
elseif v<=FOV_Y; %If Y-Motion loop hasn�t reached final FOV_Y
if Down == 1; %If user clicked down box on GUI 
fprintf(gbrl,YFOVdown);%Gcode command: Move Y-axis down to new FOV
disp('New Field of View to the DOWN complete');
pause(FOVdownupWait);%wait until move finished
elseif Up == 1; %If user clicked up box on GUI 
fprintf(gbrl,YFOVup); % Gcode command: Move Y-axis up to new FOV
disp('New Field of View to the UP complete');
pause(FOVdownupWait);%wait until move finished
end

    end 
end

 
```
