## Supplementary File 2 for "The Flexiscope: a Low Cost, Flexible, Convertible, and Modular Microscope with Automated Scanning and Micromanipulation": StageAutomation_Cam_Steppers_Variable_SetUp.m

```
%%%Set up Camera
vid = videoinput('pointgrey', 1, 'F7_RGB_1384x1032_Mode7');
src = getselectedsource(vid);
src.FrameRateMode = 'Off';
src.ExposureMode = 'Off';
src.SaturationMode = 'Off';
src.SharpnessMode = 'Off';
src.WhiteBalanceRBMode = 'Off';
src.Exposure = 0.11984;
src.GammaMode = 'Off';
src.HueMode = 'Off';
src.TriggerDelayMode = 'Off';
src.WhiteBalanceRBMode = 'Off';
src.Shutter= S; %shutter/exposure setting in milliseconds
preview(vid); %start live video


%Set up serial communication between Matlab and Arduino
gbrl=serial('COM5','BAUD', 115200);
fopen(gbrl);
%fclose(gbrl);

N5 = '_'; % image name prefix
N6 = '_%02d'; %Z stack loop value used to determine Z-stack number for image name prefix

%Auto-focus variables: see Geusebroek et al, 2000 for more details
WSize = 15; % Size of local window
N = floor(WSize/2);
sig = N/2.5;
[x,y] = meshgrid(-N:N, -N:N);
G = exp(-(x.^2+y.^2)/(2*sig^2))/(2*pi*sig);
Gx = -x.*G/(sig^2);
Gx = Gx/sum(abs(Gx(:)));
Gy = -y.*G/(sig^2);
Gy = Gy/sum(abs(Gy(:)));


%Do you want scanning to begin to the right or to the left? Based on GUI
%click box results
if Right == 1; %Right box was clicked
    a = 0;
    b = 1;
elseif Left == 1; %left box was clicked
     a = 1;
     b = 0;
end 
    
%%%Lens variables based lens drop down menu choice in GUI
if strcmpi(lens, '4X') %user clicked 4X
    ZStageDown = 'G21G91Z.02F10'; %Z-axis stepper gcode command: move stage 0.020mm down 
    
    ZStageUp = 'G21G91Z-.02F10'; %Z-axis stepper gcode command: move stage 0.020mm up
    
    XFOVright = 'G21G91X2.2F10'; %X-axis stepper gcode command: move stage 2.2mm to the right (camera perspective)
    
    XFOVleft = 'G21G91X-2.2F10'; %X-axis stepper gcode command: move stage 2.2mm to the left (camera perspective)
    
    YFOVdown = 'G21G91Y1.6F10'; %Y-axis stepper gcode command: move stage 1.6mm down (camera perspective)
    
    YFOVup = 'G21G91Y-1.6F10'; %Y-axis stepper gcode command: move stage 1.6mm up (camera perspective)
     
    ZWait = (ceil(S/1000)+1); %Time to wait in seconds during Z-stack acquisition. Shutter in milliseconds divided by 1000 plus 1 (one added incase shutter is less than a second, ceil function rounds the number up) 
    
    ZWaitcorrect = (3);  %Time to wait in seconds when Z-correction is occurring
    
    FOVrightleftWait = 15 + ZWait ;  %Time to wait in seconds when X-stepper moving

    FOVdownupWait = 15 + ZWait; %Time to wait in seconds when Y-stepper moving

elseif strcmpi(lens, '20X') %user clicked 20X
    ZStageDown = 'G21G91Z.006F10'; %Z-axis stepper gcode command: move stage 0.006mm down 
    
    ZStageUp = 'G21G91Z-.006F10';  %Z-axis stepper gcode command: move stage 0.006mm up
    
    XFOVright = 'G21G91X.43F10'; %X-axis stepper gcode command: move stage 0.43mm to the right (camera perspective)
    
    XFOVleft = 'G21G91X-.43F10'; %X-axis stepper gcode command: move stage 0.43mm to the left (camera perspective)
    
    YFOVdown = 'G21G91Y.32F10'; %Y-axis stepper gcode command: move stage 0.32mm to the down (camera perspective)
    
    YFOVup = 'G21G91Y-.32F10'; %Y-axis stepper gcode command: move stage 0.32mm to the up (camera perspective)

    ZWait = (ceil(S/1000)+2);
    
    ZWaitcorrect = (2);

    FOVrightleftWait = 5 + ZWait; 

    FOVdownupWait = 5 + ZWait;
end
```
