## Supplementary File 2 for "The Flexiscope: a Low Cost, Flexible, Convertible, and Modular Microscope with Automated Scanning and Micromanipulation": UserGuidelines_AutomatedAcquisition_StepperStage_Matlab.pdf

### User Guidelines: Automated Acquisition and Stepper Stage Scanning in Matlab

**March 2019**

Step1, 2 and 3 should be performed once when you are establishing your system.

Steps 4-6 are performed each time you want to set up a tissue sample for automated scanning.

**Step1: Prepare Arduino Uno, CNC Shield and DRV8825 Drivers**

a. Solder female jumper cables (S2) to all three steppers (S5).
b. Each axis on the CNC shield has three jumpers. Attach female shorting link (S5) to the X, Y and Z-axis (nine in total). This enables 1/32 microstepping in all axes (figure 1, B).
c. DRV8825 (S3) – Vref Adjustment

   DRV8825 current limit equation:
   current limit = Vref x 2

   Stepper (S3) current limit = 1.3A
   Vref = 650mV

   The Vref of all three DRV8825 drivers was adjusted to 650mV.

This adjustment is performed by setting up all components as seen in figure 2 and rotating the potentiometer on the driver until 600mV is seen on the oscilloscope. An in depth description of this process and the DRV8825 drivers can be found at https://www.pololu.com/product/2133.

d. Attach the drivers and steppers to the CNC shield as seen in figure 1, A. Attach the CNC shield to the Arduino Uno.
e. Download gbrl on to the Arduino Uno (S1) (https://github.com/grbl/grbl/wiki/Flashing-Grbl-to-an-Arduino).
f. Attach the 12V Power supply.

**Step2: Assemble XYZ stage for stepper motor motion control**

a. 3D print mounting and coupling components (3D5 x3, 3D6 and 3D7)
b. Assemble stage with 3D printed and MakerBeam components as seen in figure 3.
c. Test mechanical and electrical components using the universal Gcode sender (UGS) https://winder.github.io/ugs_website/.
d. Calculate the number of steps required to move each axis 1mm. This can be read directly from the micrometers. Our steppers have 200 steps per revolution. To move 1 mm the stepper must revolve twice (400 steps). At 1/32 microstepping this equates to 12800 steps/mm.
e. Adjust gbrl settings by typing $$ in to the UGS command line (figure 4).

**Step3: Determine Optimal Camera Settings within Matlab**

A. The camera used in our system for fluorescent imaging is CO23. The 'Camera Settings' commands found in 'StageAutomation_Cam__Steppers_Variable_SetUp.m' are therefore optimal for our system. If you are using a different camera you will need to open your camera in the image acquisition toolbox in Matlab (make sure you have installed appropriate add on packages for your camera: https://uk.mathworks.com/help/imaq/installing-the-support-packages-for-image-acquisition-toolbox-adaptors.html).
B. Alter the 'acquisition parameters' until you achieved the desired settings. The appropriate commands for these specific settings can be seen in 'Session Log'.

C. Input the appropriate commands into the ‘Camera Settings’ commands in ‘StageAutomation_Cam__Steppers_Variable_SetUp.m’.

**Step4: Time to do your first scan!**

A. Set up camera settings in directly in the FlyCap software as seen in figure 5 and determine optimal shutter (exposure time in ms).
B. Find your starting position within or at the edge of the tissue sample
C. Ensure sufficient travel in the X and Y-axis in the direction you want the stage to scan (for example: scanning right and down requires the X- and Y- axis to be rotated fully anticlockwise until range of travel has been reached to allow maximum travel clockwise which is the desired direction of scanning)
D. Adjust Z axis:

- Starting imgnum odd: Above most in focus region i.e. rotating Z actuator anticlockwise (stage down) so that the image comes in and out of focus
- Starting imgnum is even: Below most in focus region i.e. rotating Z actuator clockwise (stage up) so that the image comes in and out of focus

**Step5: Matlab – User settings GUI**

A. Start Matlab
B. Navigate ‘current folder’ to folder where you would like to save your images
C. Ensure the following files are in this folder:
    - StageAutomation_GUI_1.fig
    - StageAutomation_GUI_1.m
    - StageAutomation_AcquisitionMotionCommands_Steppers.m
    - StageAutomation_Cam__Steppers_Variable_SetUp.m
D. Type into Run(‘StageAutomation_GUI_1.m’) into command window. The Stage automation guided user interface (GUI) should begin (figure 6)
E. Complete all edit boxes with desired settings
F. Click Begin Acquisition
G. Make sure you’re happy with settings (variables now in workspace).

**Step6: Matlab - Live stream camera display and stepper motor GUI set up**

A. Type Run(‘StageAutomation_Cam__Steppers_Variable_SetUp.m) into command window. You should see the video preview window and additional variables in the workspace.
B. Ensure you are happy with camera view and variables in workspace (tip: if you’re not happy with any variable values simply restart from step 4 or directly alter them in workspace. For example, to change exposure time to four seconds type: S=4000’; src.Shutter= S; into command window and the live video will update itself automatically)
C. When you are ready to start scanning type Run(‘StageAutomation_AcquisitionMotionCommands_Steppers.m) into command window

ZStageUp: Z-axis stepper gcode command – move stage up

XFOVright: X-axis stepper gcode command: move stage to the right (camera perspective)

XFOVleft: X-axis stepper gcode command: move stage to the left (camera perspective)

YFOVdown: Y-axis stepper gcode command: move stage to the down (camera perspective)

YFOVup: Y-axis stepper gcode command: move stage up (camera perspective)

ZWait: Time to wait in seconds during Z-stack acquisition. Shutter in milliseconds divided by 1000 plus 1 (one added incase shutter is less than a second, ceil function rounds the number up)

ZWaitcorrect: Time to wait in seconds when Z-correction is occurring

FOVrightleftWait: Time to wait in seconds when X-stepper moving

FOVdownupWait: Time to wait in seconds when Y-stepper moving

MeanI: get mean intensity of last image captured at this FOV

**X/Y scanning and Z-stack acquisition explained:**

Figure 7 explains the workflow to acquire Z-stacks at regions of the sample in the X and Y direction. The commands used to control the specimen stage followed a logic in which the image sequence of the tissue sample is considered a two dimensional array. To acquire images of the whole tissue (or a region of interest) the stage must be moved in the X-axis a defined number of times and then moved in the Y-axis once, this cycle can then be repeated until the tissue (or region of interest) has been imaged in its entirety. Each movement in X or Y dimension reveals a new 'field of view' (FOV) and a Z-stack is subsequently acquired. The motion control and image acquisition code is explained schematically in figure 8.

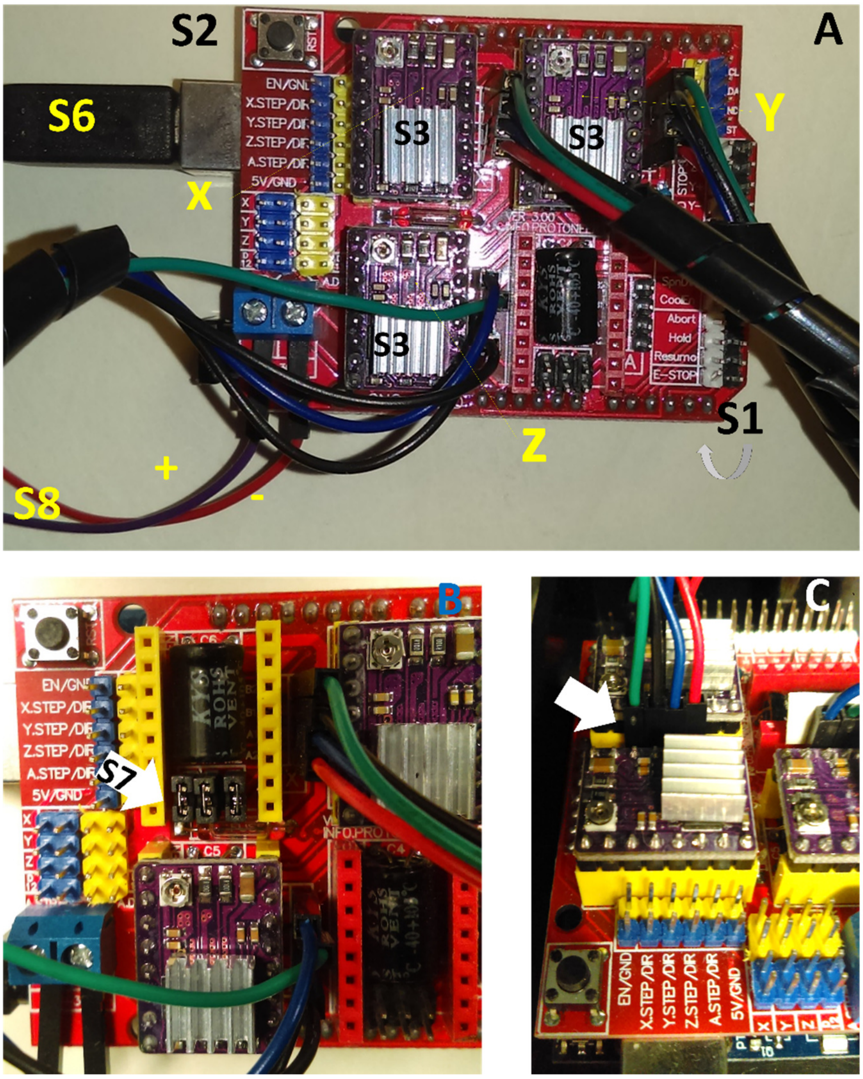


Figure 1. How to set up the Arduino Uno and CNC shield

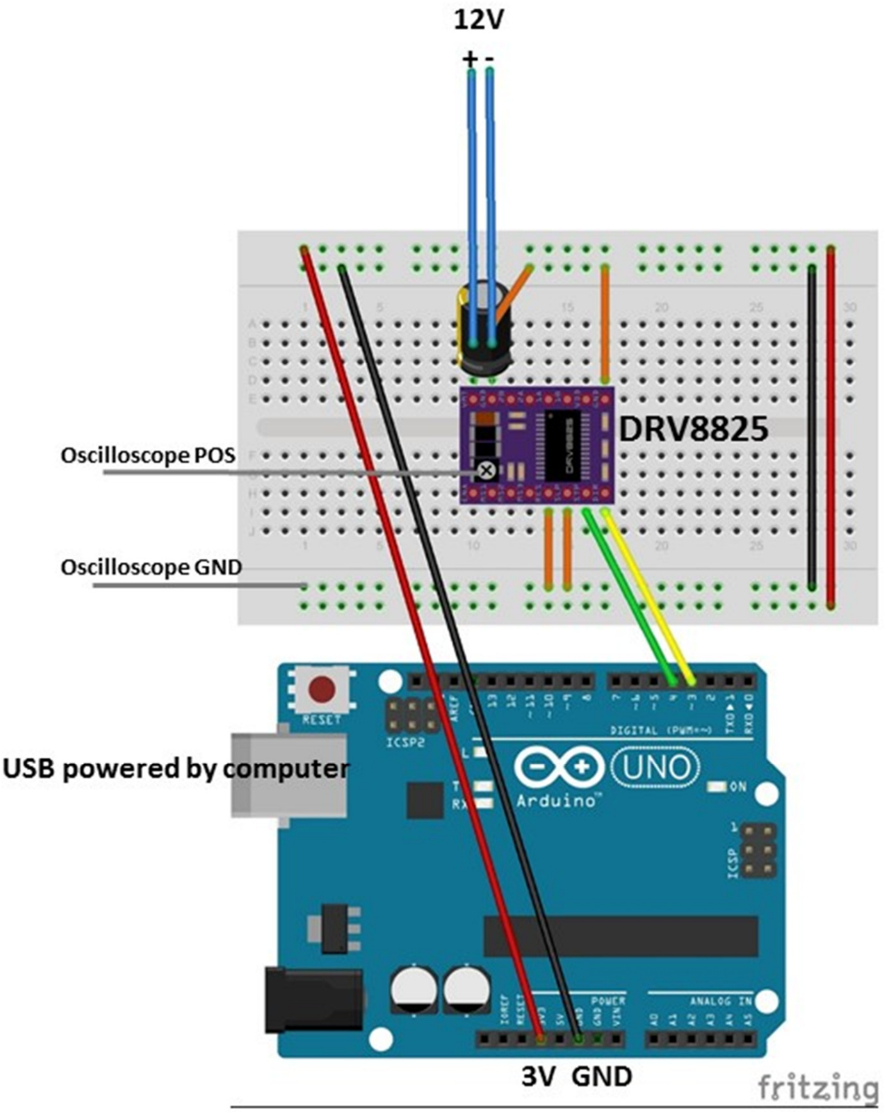


Figure 2. Wiring diagram to enable Vref adjustment of DRV8825 stepper motor drivers

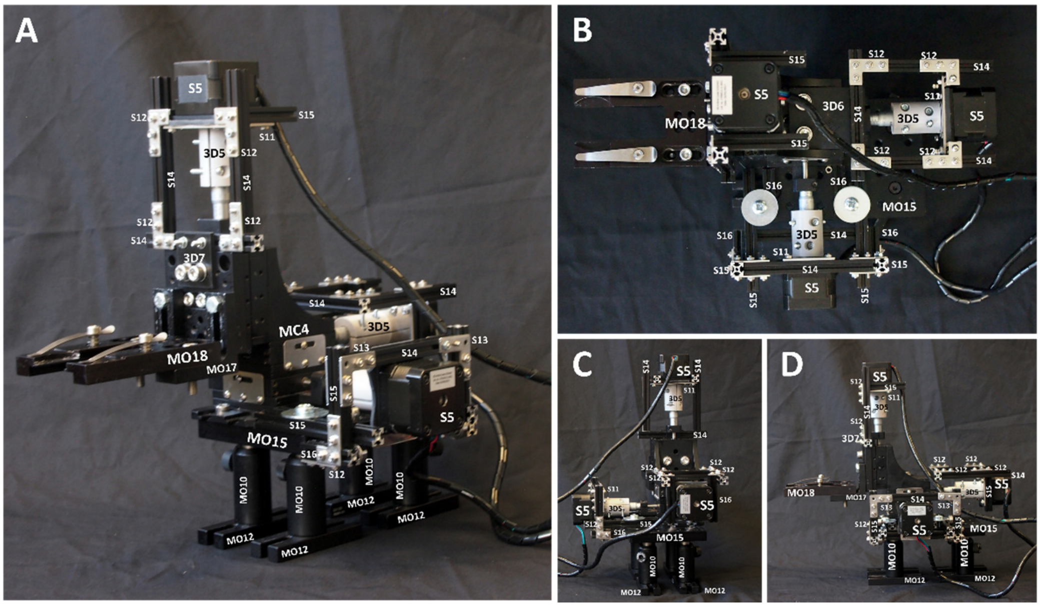


Figure 3. How to assemble the XYZ translation stage (MC4) to enable stepper motor motion (S5) control.

$0 = 10 (Step pulse time, microseconds)
$1 = 25 (Step idle delay, milliseconds)
$2 = 0 (Step pulse invert, mask)
$3 = 0 (Step direction invert, mask)
$4 = 0 (Invert step enable pin, boolean)
$5 = 0 (Invert limit pins, boolean)
$6 = 0 (Invert probe pin, boolean)
$10 = 1 (Status report options, mask)
$11 = 0.010 (Junction deviation, millimeters)
$12 = 0.002 (Arc tolerance, millimeters)
$13 = 0 (Report in inches, boolean)
$20 = 0 (Soft limits enable, boolean)
$21 = 0 (Hard limits enable, boolean)
$22 = 0 (Homing cycle enable, boolean)
$23 = 0 (Homing direction invert, mask)
$24 = 25.000 (Homing locate feed rate, mm/min)
$25 = 500.000 (Homing search seek rate, mm/min)
$26 = 250 (Homing switch debounce delay, milliseconds)
$27 = 1.000 (Homing switch pull-off distance, millimeters)
$30 = 1000 (Maximum spindle speed, RPM)
$31 = 0 (Minimum spindle speed, RPM)
$32 = 0 (Laser-mode enable, boolean)
$100 = 12800.000 (X-axis travel resolution, step/mm)
$101 = 12800.000 (Y-axis travel resolution, step/mm)
$102 = 12800.000 (Z-axis travel resolution, step/mm)
$110 = 30.000 (X-axis maximum rate, mm/min)
$111 = 30.000 (Y-axis maximum rate, mm/min)
$112 = 30.000 (Z-axis maximum rate, mm/min)
$120 = 1.000 (X-axis acceleration, mm/sec^2)
$121 = 1.000 (Y-axis acceleration, mm/sec^2)
$122 = 1.000 (Z-axis acceleration, mm/sec^2)
$130 = 13.000 (X-axis maximum travel, millimeters)
$131 = 13.000 (Y-axis maximum travel, millimeters)
$132 = 13.000 (Z-axis maximum travel, millimeters)

Figure 4. gbrl settings in UGS

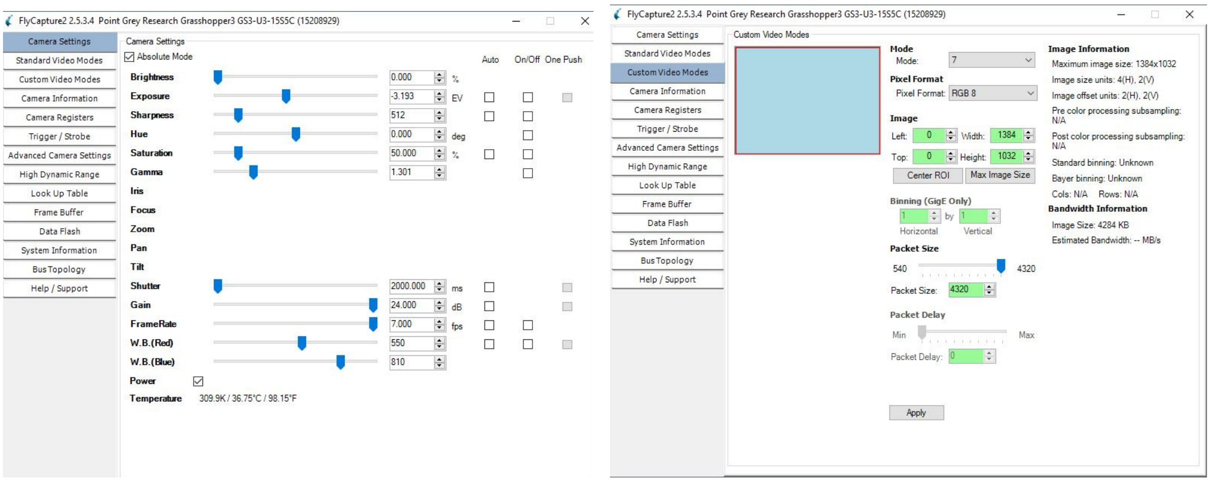


Figure 5. FlyCap settings for CO23 for fluorescent imaging

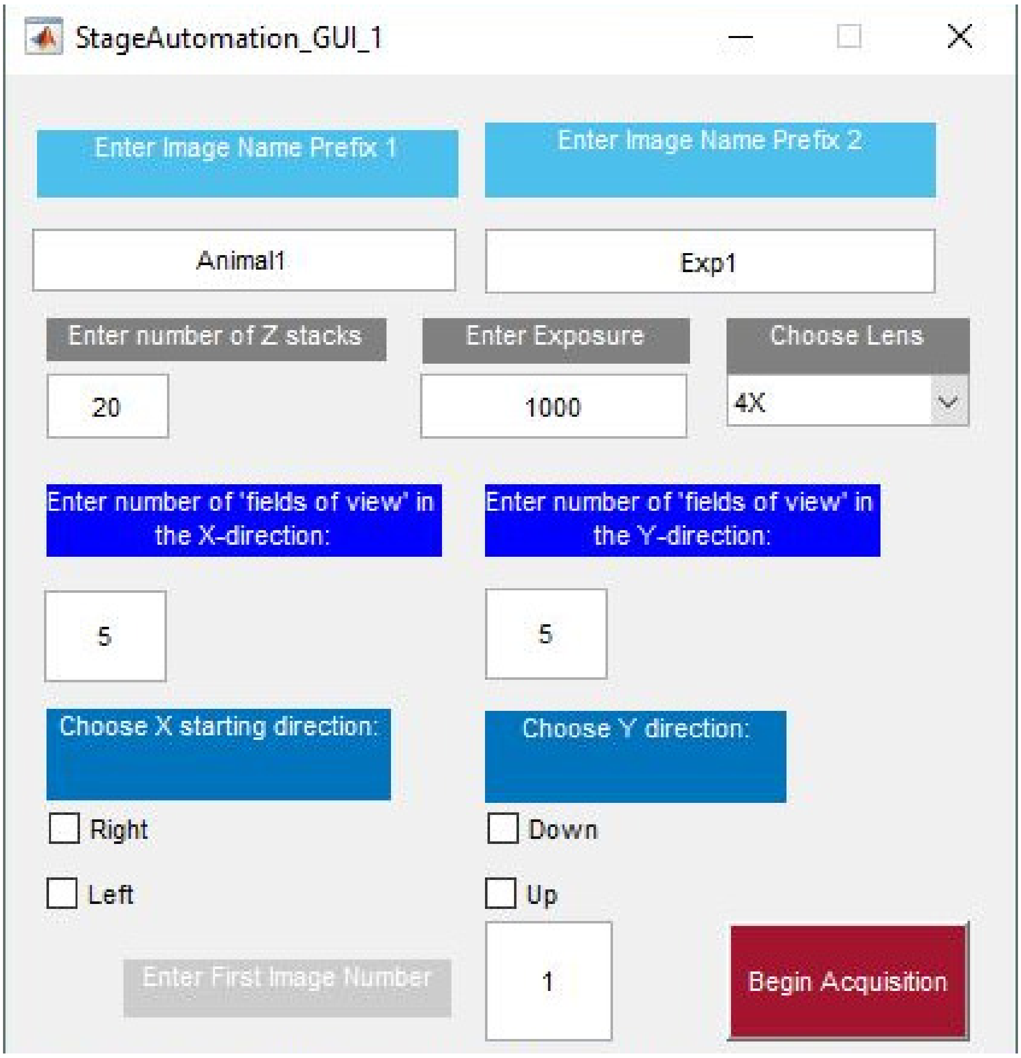


Figure 6. Automated stepper stage motion control GUI in Matlab

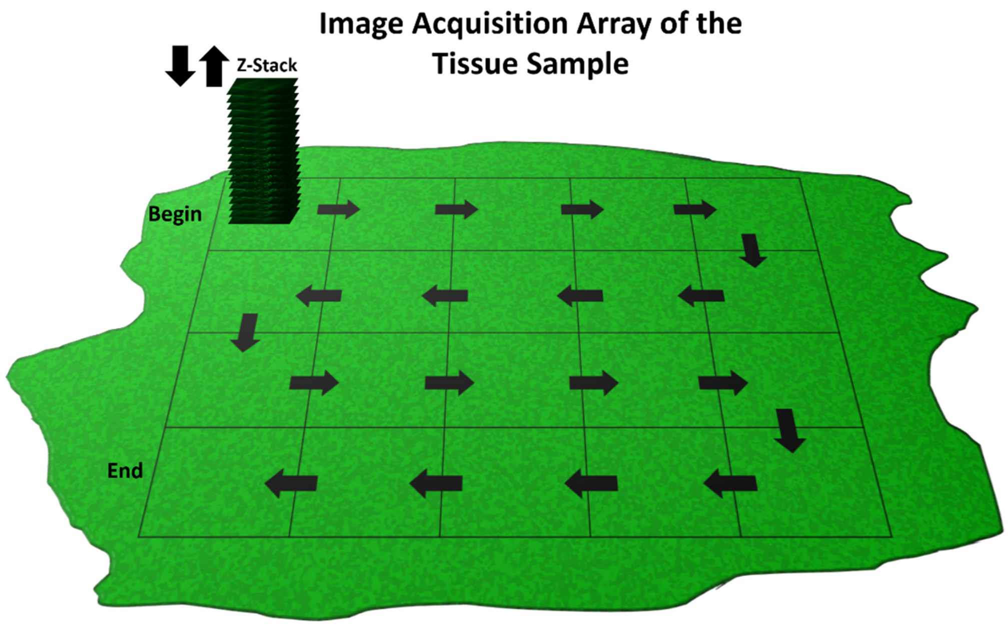

Figure 7. Schematic representation of the automated stage motion control

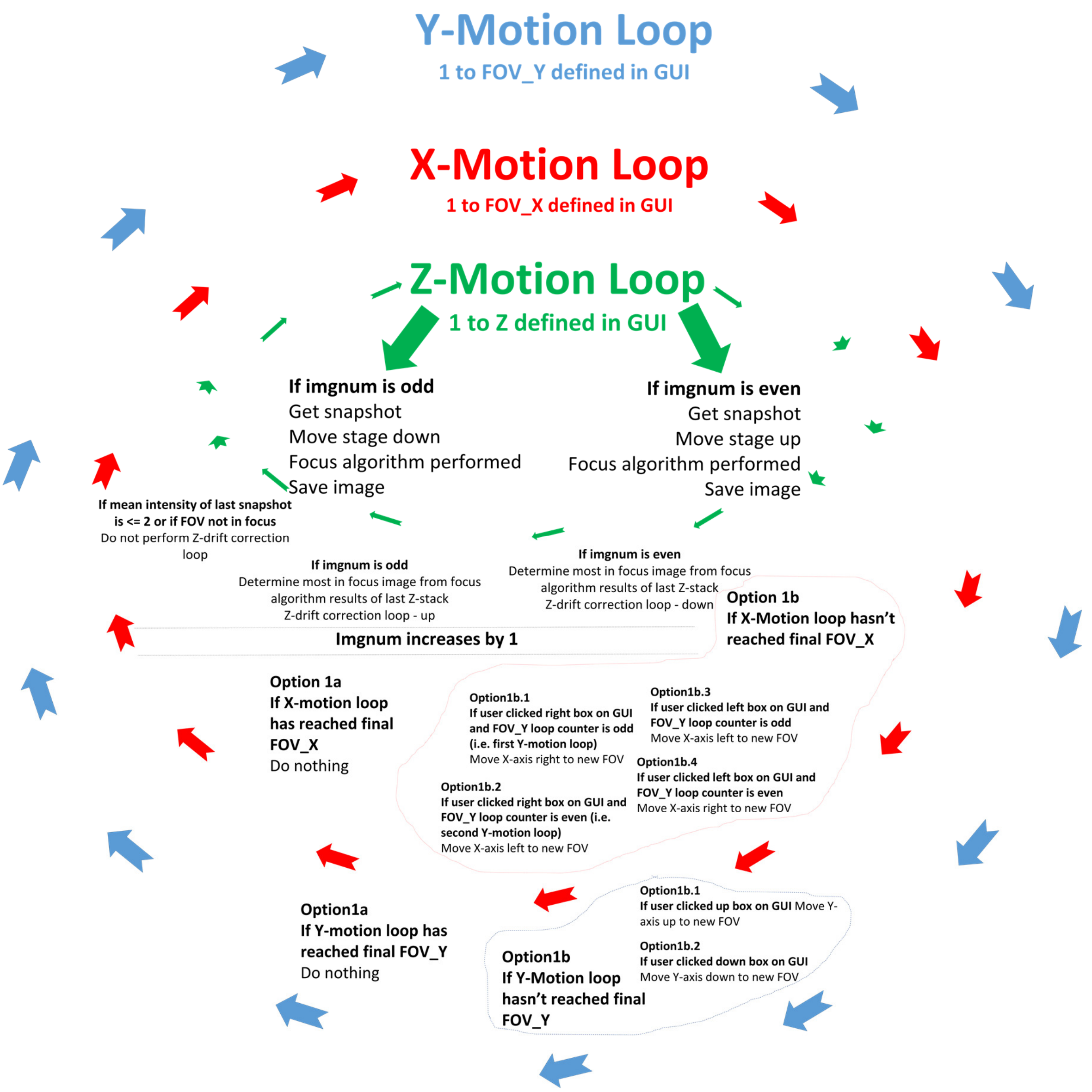


Figure 8. Schematic representation of the automated stepper motion control and image acquisition code.
