## Supplementary File 3 for "The Flexiscope: a Low Cost, Flexible, Convertible, and Modular Microscope with Automated Scanning and Micromanipulation": AutomatedAcquisition_StepperStage_Python.py

```
import PySpin
import serial
import time
import math
import numpy as np
import cv2
#in order to use the FLIR spinnaker packages with python, PySpin mist also be imported
#PySpin is part of the Spinnaker SDK available freely from FLIR to all FLIR macine vision camera owners
#All code derived from the Spinnaker SDK has been removed from this script to comply with the terms of the 
#SDK licence agreement. Where code has been removed, it has been replaced with comments describing the 
#PySpin-specific functions, which can be replaced by users with access to the SDK

#set up arduino (controls the stepper stage)
ser = serial.Serial()
ser.baudrate = 115200
ser.port = 'COM5' #Change this (this depends on where the USB is inserted)
ser #prints info
ser.open() #connect to arduino
ser.is_open # Checks it is open


lens = '20X' #Change this 
S = 1000 #Change this. Shutter/exposure setting in milliseconds as you defined in FlyCap or Spinnaker
N1= 'Animal1' #Change this. Image prefix 1 
N2= 'Exp3' #Change this. Image prefix 2
Z= 3 #Change this. Number of Z-stacks
FOV_X = 2 #Change this. Number of field of views (FOV) in the X-axis i.e. how many time the X-axis will move in a row.
FOV_Y = 2 #Change this. Number of field of views (FOV) in the Y-axis i.e. how many time the Y-axis will move after each row.
Right = 1 #Change this. Keep this as 1 and 'Left' as 0 if you want to start by scanning to the right.
Left = 0 #Change this. Change this to 1 and 'Right' to 0 if you want to start by scanning to the left.
Down = 1 #Change this. Keep this as 1 and 'Up' as 0 if you want to start by scanning down.
Up = 0 #Change this. Change this to 1 and 'Down' to 0 if you want to start by scanning up.

N5= '_' # for contructing the filename later
N8= '.tif' #Change this. This can be changed to .jpg or .png if required


#Setting up variables based on above parameters
if Right == 1: #start moving right first
    a = 1
    b = 0
elif Left == 1: #start moving left first
    a = 0
    b = 1
if lens=='4X': #If I defined the lens at 4X at the beginning
     
    ZStageDown = b'G21G91Z.02F1000\n'    #Z-axis stepper gcode command: move stage 0.020mm down

    ZStageUp = b'G21G91Z-.02F1000\n'    #Z-axis stepper gcode command: move stage 0.020mm up

    XFOVright = b'G21G91X2.2F1000\n'   #X-axis stepper gcode command: move stage 2.2mm to the right (camera perspective)

    XFOVleft = b'G21G91X-2.2F1000\n'    #X-axis stepper gcode command: move stage 2.2mm to the left (camera perspective)

    YFOVdown = b'G21G91Y1.6F1000\n'   #Y-axis stepper gcode command: move stage 1.6mm down (camera perspective)

    YFOVup = b'G21G91Y-1.6F1000\n'  #Y-axis stepper gcode command: move stage 1.6mm up (camera perspective)

    ZWait = (math.ceil(S / 1000) + 1)    #Time to wait in seconds during Z-stack acquisition. Shutter in milliseconds divided by 1000 plus 1 (one added incase shutter is less than a second, ceil function rounds the number up)

    FOVrightleftWait = 15 + ZWait    #Time to wait in seconds when X-stepper moving

    FOVdownupWait = 15 + ZWait    #Time to wait in seconds when Y-stepper moving
    
elif lens=='20X': #If I defined the lens at 20X at the beginning
        
    ZStageDown = b'G21G91Z.006F1000\n'    #Z-axis stepper gcode command: move stage 0.006mm down

    ZStageUp = b'G21G91Z-.006F1000\n'    #Z-axis stepper gcode command: move stage 0.006mm up

    XFOVright = b'G21G91X.43F1000\n'    #X-axis stepper gcode command: move stage 0.43mm to the right (camera perspective)

    XFOVleft = b'G21G91X-.43F1000\n'    #X-axis stepper gcode command: move stage 0.43mm to the left (camera perspective)

    YFOVdown = b'G21G91Y.32F1000\n'    #Y-axis stepper gcode command: move stage 0.32mm to the down (camera perspective)

    YFOVup = b'G21G91Y-.32F1000\n'    #Y-axis stepper gcode command: move stage 0.32mm to the up (camera perspective)

    ZWait = (math.ceil(S / 1000) + 2) #Time to wait in seconds during Z-stack acquisition. Shutter in milliseconds divided by 1000 plus 1 (one added incase shutter is less than a second, ceil function rounds the number up)

    FOVrightleftWait = 5 + ZWait #Time to wait in seconds when X-stepper moving

    FOVdownupWait = 5 + ZWait #Time to wait in seconds when Y-stepper moving

#
    
#This code block described the camera set up using PySpin    
# set up the camera - this script assumes a single USB3 camera is connected

#the variable "system" is defined as a system instance

#the variable "cam_list" is defined as the list of cameras
cam = cam_list[0]

# initiate camera (cam)
time.sleep(1)

#the variable "nodemap" is defined as the camera node map

#set the acquisition mode to continuous
#End of camera set up code block
#

#start the motion control/acquisition loop
imgnum =1 #Change this
for v in range(1, FOV_Y+1): #Y-motion FOV loop
    for i in range(1, FOV_X+1):#X-motion FOV loop
        for z in range(1, Z+1): #Z stack acquisition loop
            FM_master = []
            if (imgnum % 2) == 1: #determine if imgnum is odd or even
                #this occurs when imgnum is odd
                print('take photo')
                #insert PySpin code here
                #begin acquisition (cam)
                #insert PySpin code here
                #define variable "image_result": retrieve the next image from cam
                print('move Z stage down')
                ser.write(ZStageDown) #move Z stage down
                time.sleep(ZWait) #wait in seconds

                print('name photo')
                #insert PySpin code here
                #image_converted = Convert the image (image_result) to BGR8
                N4 = str(imgnum) #image number as a string
                N6 = str(z) #Z-stack number as a string
                filename  = N1 + N5 + N2 + N5 + lens + N5 + N4 + N5 + N6 + N8 #Combine strings to form filename
                print('save photo')
                #insert PySpin code here
                #save image (image_converted)
                print('release image')
                #insert PySpin code here
                #release image (image_result)
                #insert PySpin code here
                #end acquisition (cam)
                
                print('calculate focus')
                img = cv2.imread(filename) #read in image which was just saved
                gray = cv2.cvtColor(img, cv2.COLOR_BGR2GRAY)#convert to gray
                FM = cv2.Laplacian(gray, cv2.CV_64F).var()#Determine focus measure
                print(FM)
                
                print('store all focus measurements')
                FM_master.append(FM) #Save all the focus values for each image in a Z-stack as a list
                
                print('calculate mean intensity')
                MeanI = np.mean(gray) #mean intensity of the last image saved
                print(MeanI)

            else: #this occurs when imgnum is even
                print('take photo')
                #insert PySpin code here
                #begin acquisition (cam)
                #insert PySpin code here
                #define variable "image_result": retrieve the next image from cam
                print('move Z stage up')
                ser.write(ZStageUp) #move Z stage up
                time.sleep(ZWait) #wait in seconds
                
                print('name photo')
                #insert PySpin code here
                #image_converted = Convert the image (image_result) to BGR8
                N4 = str(imgnum) #image number as a string
                N6=str(z) #Z-stack number as a string
                filename  = N1 + N5 + N2 + N5 + lens + N5 + N4 + N5 + N6 + N8 #Combine strings to form filename
                
                print('save photo')
                #insert PySpin code here
                #save image (image_converted)
                
                print('release image')
                #insert PySpin code here
                #release image (image_result)
                #insert PySpin code here
                #end acquisition (cam)
                
                print('calculate focus')
                img = cv2.imread(filename) #read in image which was just saved
                gray = cv2.cvtColor(img, cv2.COLOR_BGR2GRAY)#convert to gray
                FM = cv2.Laplacian(gray, cv2.CV_64F).var()#Determine focus measure
                print(FM)
                
                print('store all focus measurements')
                FM_master.append(FM)  #Save all the focus values for each image in a Z-stack as a list
                
                print('calculate mean intensity')
                MeanI = np.mean(gray) #mean intensity of the last image saved
                print(MeanI)
                
    
        #Correcting Z-axis to ensure region of interest remains in focus
        F2= (FM_master.index(max(FM_master)))+1 #%find the max value in FMmaster and where it is located in the list. This tells us the Z-stack which was most in focus. We want to keep The most in focus region in the middle of the Z-stack.
        Zdrift = math.ceil(((Z/2))-F2) #Z/2 is the middle of the Z-Stack. F2 is the most in focus image from the last Z-stack. For example if Z is 20 we want the most in focus Z-image to be 10.
        print(Zdrift)
        if (MeanI <=2 or Zdrift<=1): #Change this! if Z-correction value is less than 1 or if mean of last image is less than 2 (no tissue in view) then no correction is required 
             print ('no Z adjust required')
        else:
            if (imgnum % 2) == 1: #Only performs this correction if stage has just moved down i.e. imgnum is odd
                for f in range(1, Zdrift+1): #Z-drift correction loop
                    print('move stage up') 
                    ser.write(ZStageUp) #move Z stage up
                    time.sleep(ZWait) #wait in seconds
                     
            elif (imgnum % 2) == 0: #Only performs this correction if stage has just moved up i.e. imgnum is even
                for f in range(1, Zdrift+1): #Z-drift correction loop
                    print('move stage down')
                    ser.write(ZStageDown) #move Z stage down
                    time.sleep(ZWait) #wait in seconds
                     
        imgnum = imgnum +1 #imgnum increases by 1 with each new FOV
        print(imgnum)
        if i == FOV_X: #If X-motion loop has reached final FOV_X
            print('end of row')
        else: #If X-Motion loop hasnt reached final FOV_X
            Dir = v % 2 #Is Y-motion loop value odd or even?
            if Dir == a: # I chose to start moving right at the beginning. 
                print('new FOV to the right')
                ser.write(XFOVright) #Move X-axis right to new FOV
                time.sleep(FOVrightleftWait)
            elif Dir == b: # I chose to start moving left at the beginning. 
                print('new FOV to the left')
                ser.write(XFOVleft) #Move X-axis left to new FOV
                time.sleep(FOVrightleftWait)
                    
    if v == FOV_Y: #If Y-motion loop has reached final FOV_Y
        print('do nothing')
    else: #If Y-Motion loop hasnt reached final FOV_Y
        if Down == 1: #If I chose to move down at the beginning
            print('new FOV down')
            ser.write(YFOVdown) #Move Y-axis down to new FOV
            time.sleep(FOVdownupWait)
        elif Up == 1: #If I chose to move up at the beginning
            print ('new FOV Up')
            ser.write(YFOVup) #Move Y-axis up to new FOV
            time.sleep(FOVdownupWait)

#This code block uses PySpin commands to shut down the camera

#deinitialise camera(cam)
del cam
#delete cam variable

#clear camera list

#release system instance
print('Finished')
```
