## Supplementary File 3 for "The Flexiscope: a Low Cost, Flexible, Convertible, and Modular Microscope with Automated Scanning and Micromanipulation": ReadMe.txt

This folder contains one Python file to control the automated stepper scanning stage. Guidelines to assemble and run these files can be found in 'UserGuidelines_AutomatedAcquisition_StepperStage_Python.pdf'.

Python file:

AutomatedAcquisition_StepperStage_Python.py
