## Supplementary File 3 for "The Flexiscope: a Low Cost, Flexible, Convertible, and Modular Microscope with Automated Scanning and Micromanipulation": UserGuidelines_AutomatedAcquisition_StepperStage_Python.docx

**User Guidelines: Automated**

**Acquisition and Stepper Stage**

**Scanning in Python**

**March 2019**

Step1, 2 and 3 should be performed once when you are establishing your system.

Steps 4 is performed each time you want to set up a tissue sample for automated scanning.

**Step1: Prepare Arduino Uno, CNC Shield and DRV8825 Drivers**

a. Solder female jumper cables (S2) to all three steppers (S5).
b. Each axis on the CNC shield has three jumpers. Attach female shorting link (S5) to the X, Y and Z-axis (nine in total). This enables 1/32 microstepping in all axes (figure 1, B).
c. DRV8825 (S3) – Vref Adjustment

DRV8825 current limit equation:

current limit = Vref x 2

Stepper (S3) current limit = 1.3A

Vref = 650mV

The Vref of all three DRV8825 drivers was adjusted to 650mV.

This adjustment is performed by setting up all components as seen in figure 2 and rotating the potentiometer on the driver until 600mV is seen on the oscilloscope. An in depth description of this process and the DRV8825 drivers can be found at https://www.pololu.com/product/2133.

d. Attach the drivers and steppers to the CNC shield as seen in figure 1, A. Attach the CNC shield to the Arduino Uno.
e. Download gbrl on to the Arduino Uno (S1) (https://github.com/grbl/grbl/wiki/Flashing-Grbl-to-an-Arduino)
f. Attach the 12V Power supply and USB to the computer.

**Step2: Assemble XYZ stage for stepper motor motion control**

a. 3D print mounting and coupling components (3D5 x3, 3D6 and 3D7)
b. Assemble stage with 3D printed and MakerBeam components as seen in figure 3.
c. Test mechanical and electrical components using the universal Gcode sender (UGS) <https://winder.github.io/ugs_website/>.
d. Calculate the number of steps required to move each axis 1mm. This can be read directly from the micrometers. Our steppers have 200 steps per revolution. To move 1 mm the stepper must revolve twice (400 steps). At 1/32 microstepping this equates to 12800 steps/mm.
e. Adjust gbrl settings by typing $$ in to the UGS command line (figure 4).

**Step3: Install Anaconda (Python 3.6), Spinnaker SDK and relevant libraries**

A. Go to <https://repo.continuum.io/archive/> and download relevant anaconda installation file for your operating system (I used Anaconda3-5.2.0-Windows-x86_64.exe).
B. Go to <https://www.ptgrey.com/support/downloads> and download the latest version of Spinnaker SDK (I used: Anaconda3-5.2.0-Windows-x86_64.exe) and the latest Spinnaker for Python library (I used: spinnaker_python-1.20.0.15-cp36-cp36m-win_amd64.whl).
C. Open the anaconda command prompt and install pySerial and OpenCV.
D. Open Spyder (included in the anaconda installation) and open ‘AutomatedAcquisition_StepperStage_Python.py’
E. Ensure all libraries are installed by running lines 1-7.
F. Set up camera directly in the FlyCap or Spinnaker software as seen in figure 5 and determine optimal shutter (exposure time in ms).
G. Go back to Spyder and go to lines 12-31. Any variables which include ‘#Change this’ should be altered to suit your parameters. Also go to line 169, 188, 223 and 255.

**Step4: Time to do your first scan!**

A. Set up your camera directly in the FlyCap or Spinnaker software.
B. Find your starting position within or at the edge of the tissue sample
C. Ensure sufficient travel in the X and Y-axis in the direction you want the stage to scan (for example: scanning right and down requires the X- and Y- axis to be rotated fully anticlockwise until range of travel has been reached to allow maximum travel clockwise which is the desired direction of scanning)
D. Adjust Z axis:

- Starting imgnum odd: Above most in focus region i.e. rotating Z actuator anticlockwise (stage down) so that the image comes in and out of focus
- Starting imgnum is even: Below most in focus region i.e. rotating Z actuator clockwise (stage up) so that the image comes in and out of focus

E. Go back to Spyder and run ‘AutomatedAcquisition_StepperStage_Python.py’ (alter variables from Step 3 G. if required).

**Variables Explained:**

S: exposure time in milliseconds

lens: lens magnification defined in GUI

Z: Number of Z Stacks as defined by user in GUI

imgnum: imgnum is an important variable for image nomenclature and also used to determine whether Z-Stack is achieved by moving Stage up or down. Even imgnum moves stage up while odd imgnum moves stage down.

Left: if this variable is equal to 1 X-motion scanning will begin to the left (it looks like the FOV is moving to the left from the camera’s perspective - clockwise X-stepper actuation- stage physically moving right). If this variable is equal to 0, X-motion scanning will begin to the right. Defined in GUI with tick boxes. Movement left is from my camera’s perspective, this may be opposite on your system.

Right: if this variable is equal to 1 X-motion scanning will begin to the right (it looks like the FOV is moving to the right from the camera’s perspective - anticlockwise X-stepper actuation- stage physically moving left). If this variable is equal to 0, X-motion scanning will begin to the left. Defined in GUI with tick boxes. Movement right is from my camera’s perspective, this may be opposite on your system

Up: Do not confuse up in the Y-axis with up in the Z-axis. Up in the Y-axis is technically moving the stage forward (clockwise Y-stepper actuation) but from the camera perspective it looks like the FOV is moving up so this naming stuck. If this variablemis equal to 1 Y-motion scanning will continue up. If this variable is equal to 0, Y-motion scanning will continue down. Defined in GUI with tick boxes. Movement up is from my camera’s perspective, this may be opposite on your system.

Down: Do not confuse down in the Y-axis with down in the Z-axis. Down in the Y-axis is technically moving the stage backward (anticlockwise Y-stepper actuation) but from the camera perspective it looks like the FOV is moving down so this naming stuck. If this variable is equal to 1 Y-motion scanning will continue down. If this variable is equal to 0, Y-motion scanning will continue up. Defined in GUI with tick boxes. Movement down is from my camera’s perspective, this may be opposite on your system

FOV_X: X-stepper number of new FOVs aka number of times X-motion loop will repeat

FOV_Y: Y-stepper number of new FOVs aka number of times Y-motion loop will repeat

ZStageDown: Z-axis stepper gcode command – move stage down

ZStageUp: Z-axis stepper gcode command – move stage up

XFOVright: X-axis stepper gcode command: move stage to the right (camera perspective)

XFOVleft: X-axis stepper gcode command: move stage to the left (camera perspective)

YFOVdown: Y-axis stepper gcode command: move stage to the down (camera perspective)

YFOVup: Y-axis stepper gcode command: move stage up (camera perspective)

ZWait: Time to wait in seconds during Z-stack acquisition. Shutter in milliseconds divided by 1000 plus 1 (one added incase shutter is less than a second, ceil function rounds the number up)

FOVrightleftWait: Time to wait in seconds when X-stepper moving

FOVdownupWait: Time to wait in seconds when Y-stepper moving

**X/Y scanning and Z-stack acquisition explained:**

Figure 6 explains the workflow to acquire Z-stacks at regions of the sample in the X and Y direction. The commands used to control the specimen stage followed a logic in which the image sequence of the tissue sample is considered a two dimensional array. To acquire images of the whole tissue (or a region of interest) the stage must be moved in the X-axis a defined number of times and then moved in the Y-axis once, this cycle can then be repeated until the tissue (or region of interest) has been imaged in its entirety. Each movement in X or Y dimension reveals a new ‘field of view’ (FOV) and a Z-stack is subsequently acquired. The motion control and image acquisition code is explained schematically in figure 7.

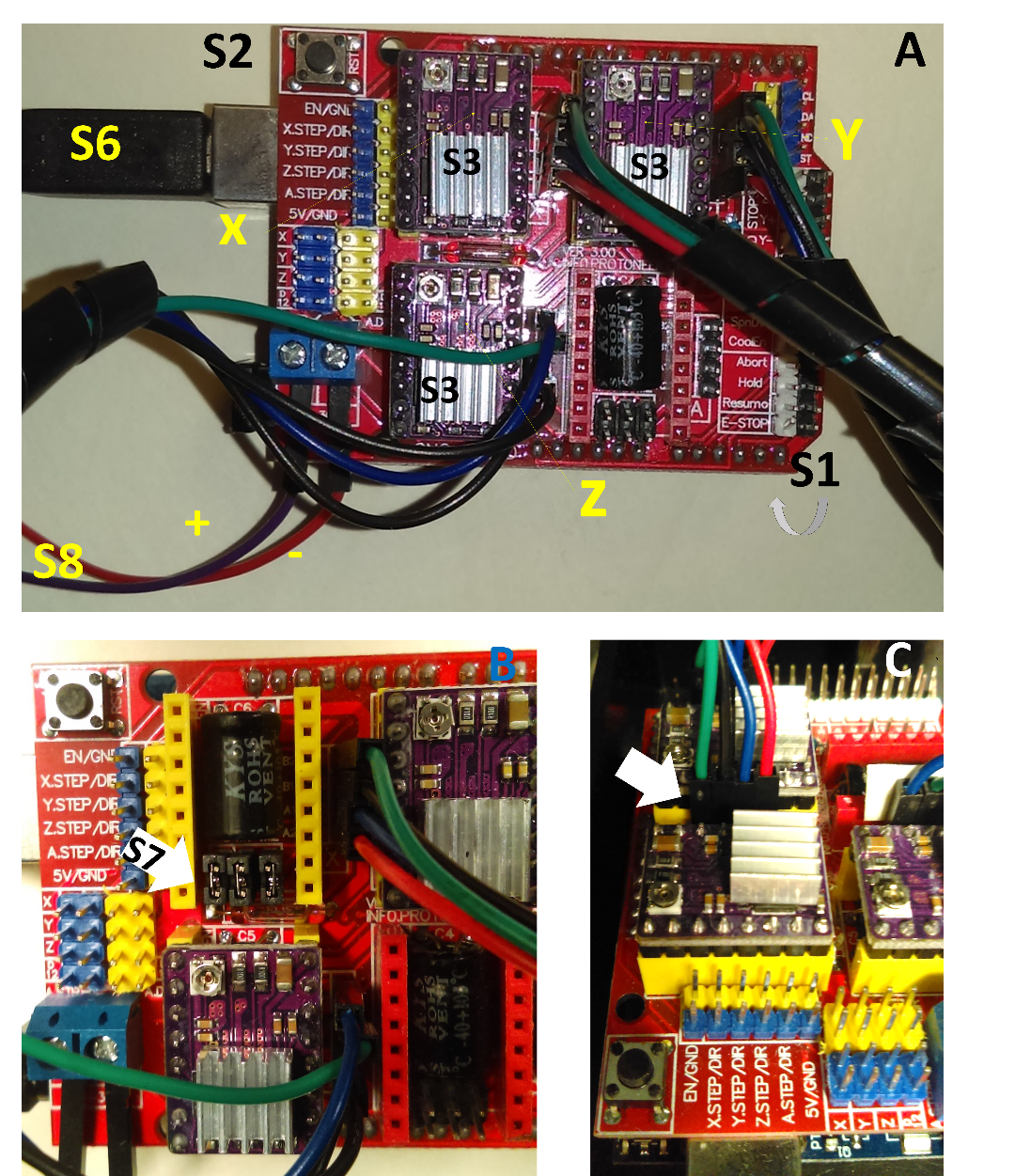

Figure 1. How to set up the Arduino Uno and CNC shield

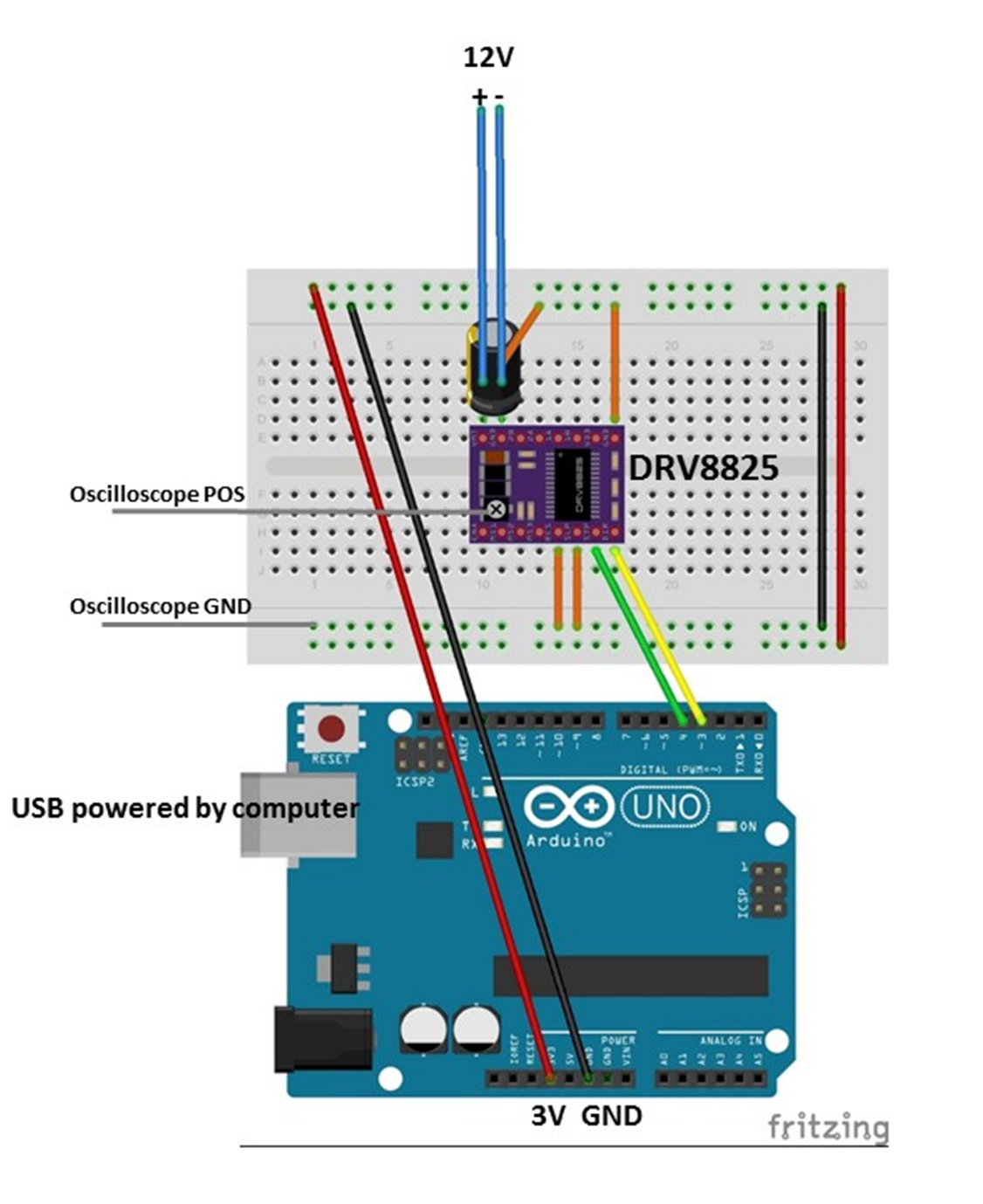

Figure 2. Wiring diagram to enable Vref adjustment of DRV8825 stepper motor drivers

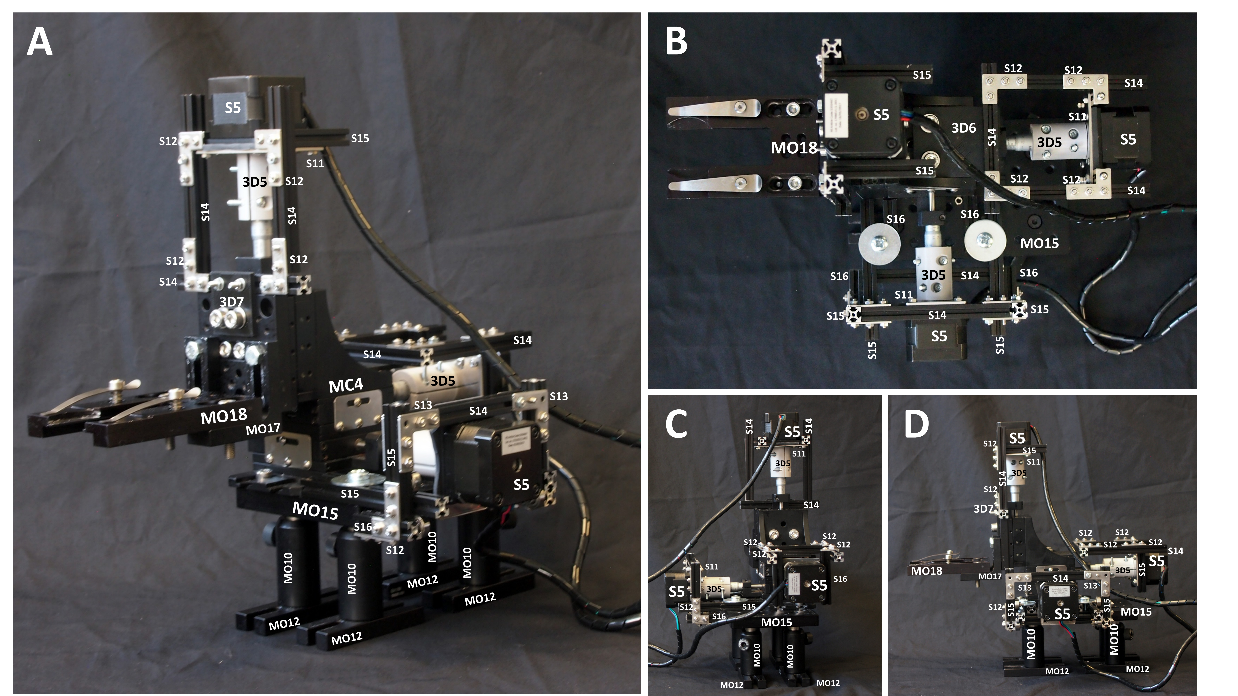

Figure 3. How to assemble the XYZ translation stage (MC4) to enable stepper motor motion (S5) control.

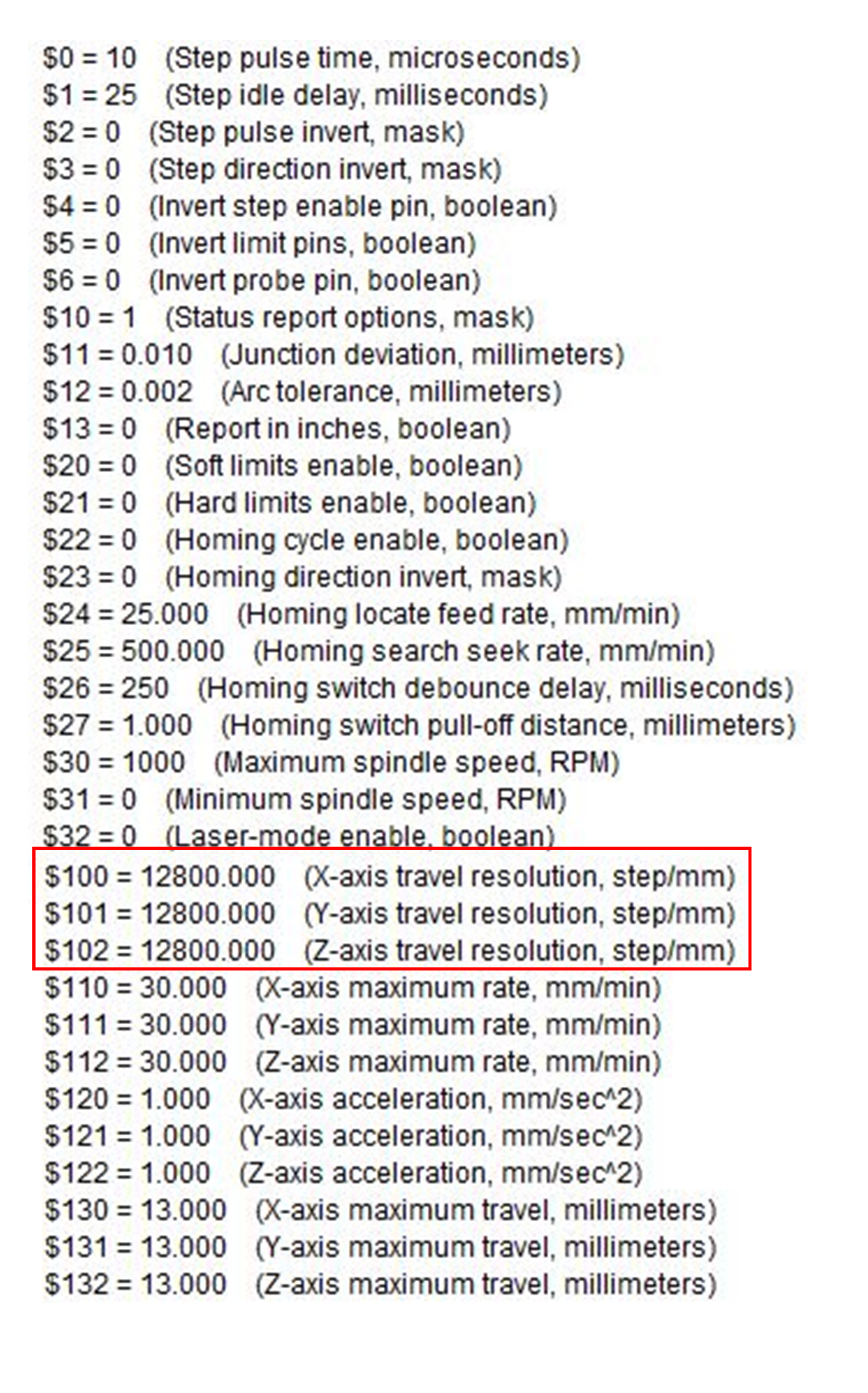

Figure 4. gbrl settings in UGS

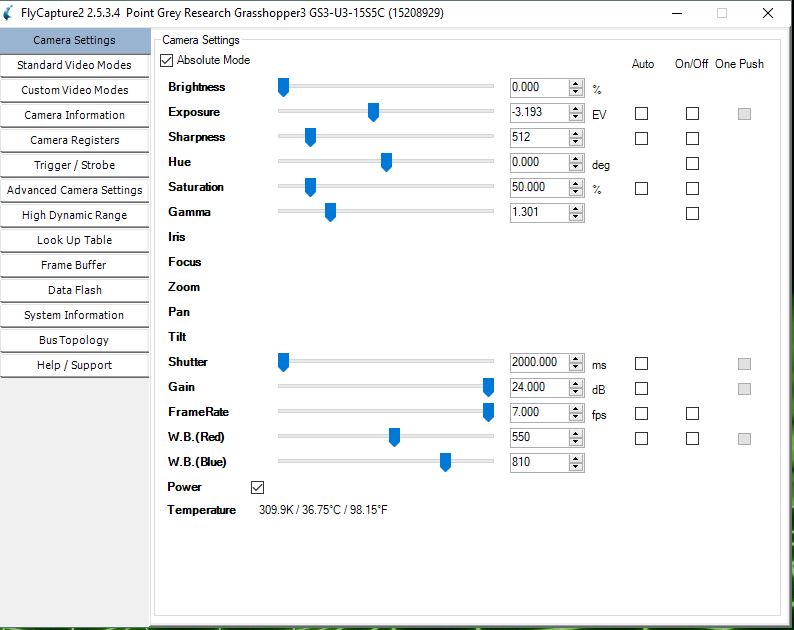

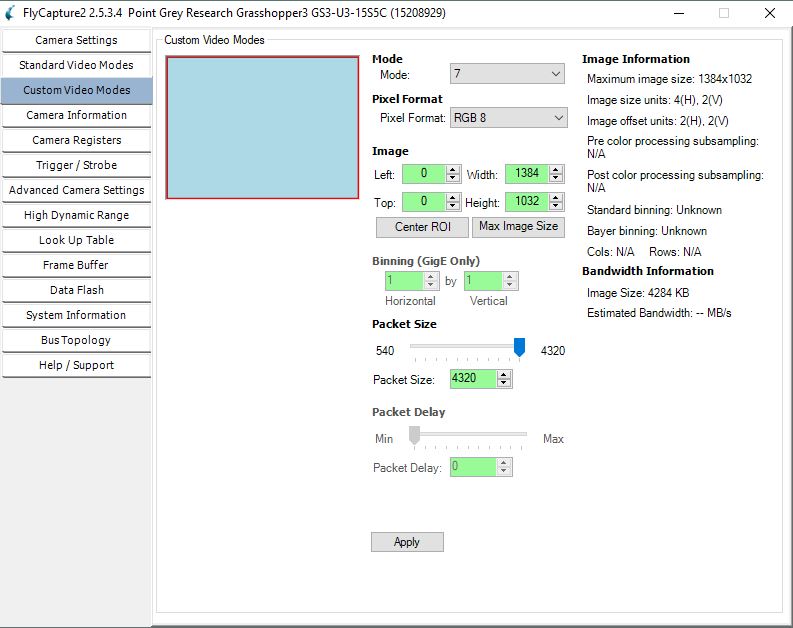

Figure 5. FlyCap settings for CO23 for fluorescent imaging

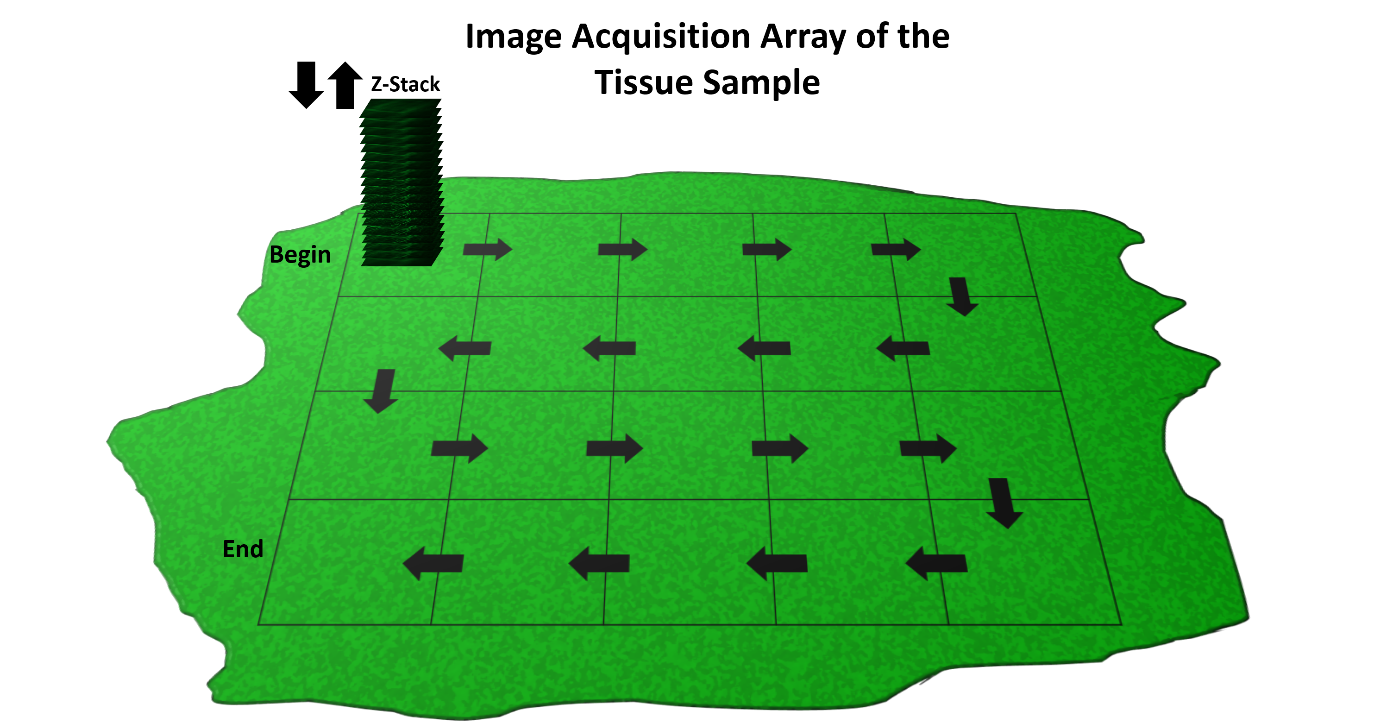

Figure 6. Schematic representation of the automated stage motion control

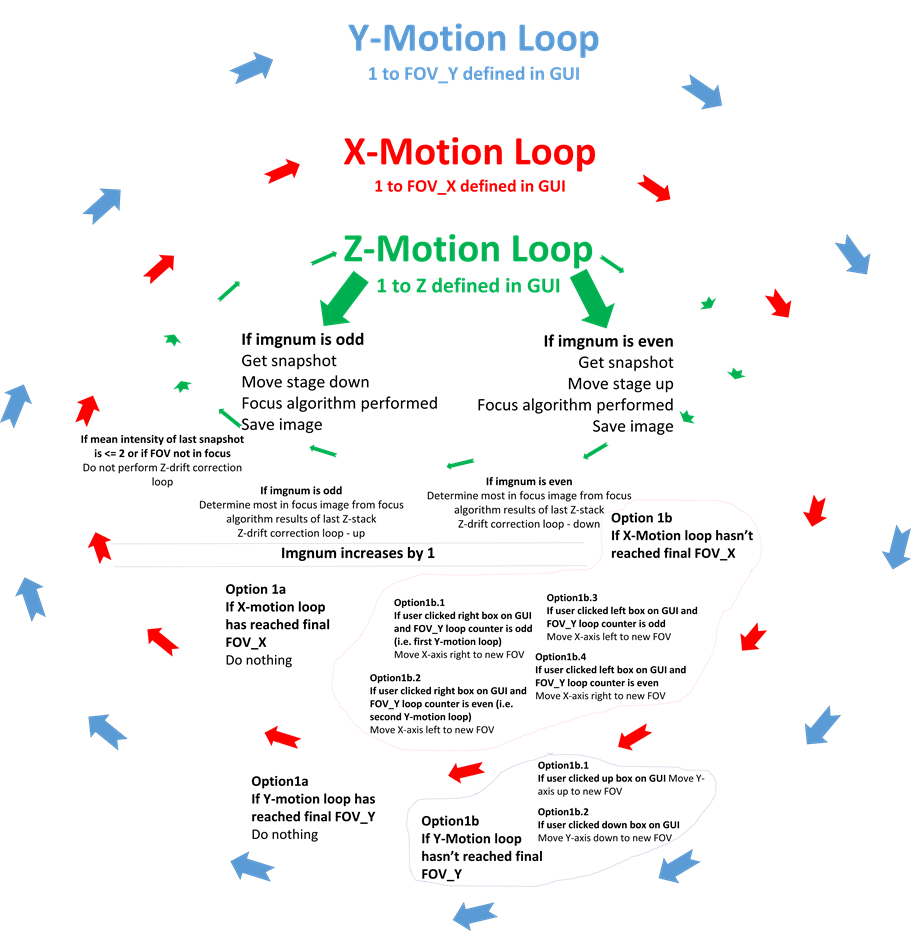

Figure 7. Schematic representation of the automated stepper stage motion control and image acquisition code.
