## Supplementary File 3 for "The Flexiscope: a Low Cost, Flexible, Convertible, and Modular Microscope with Automated Scanning and Micromanipulation": UserGuidelines_AutomatedAcquisition_StepperStage_Python.pdf

FOVrightleftWait: Time to wait in seconds when X-stepper moving

FOVdownupWait: Time to wait in seconds when Y-stepper moving

**X/Y scanning and Z-stack acquisition explained:**

Figure 6 explains the workflow to acquire Z-stacks at regions of the sample in the X and Y direction. The commands used to control the specimen stage followed a logic in which the image sequence of the tissue sample is considered a two dimensional array. To acquire images of the whole tissue (or a region of interest) the stage must be moved in the X-axis a defined number of times and then moved in the Y-axis once, this cycle can then be repeated until the tissue (or region of interest) has been imaged in its entirety. Each movement in X or Y dimension reveals a new ‘field of view’ (FOV) and a Z-stack is subsequently acquired. The motion control and image acquisition code is explained schematically in figure 7.

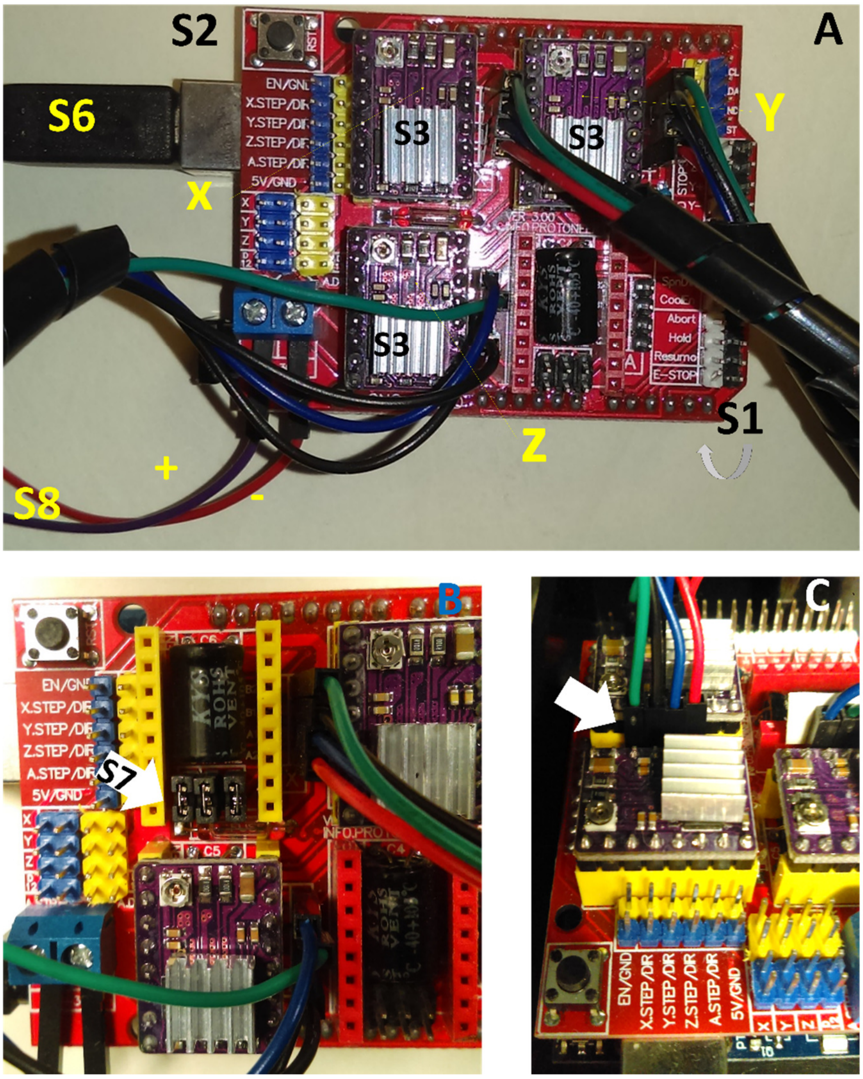


Figure 1. How to set up the Arduino Uno and CNC shield

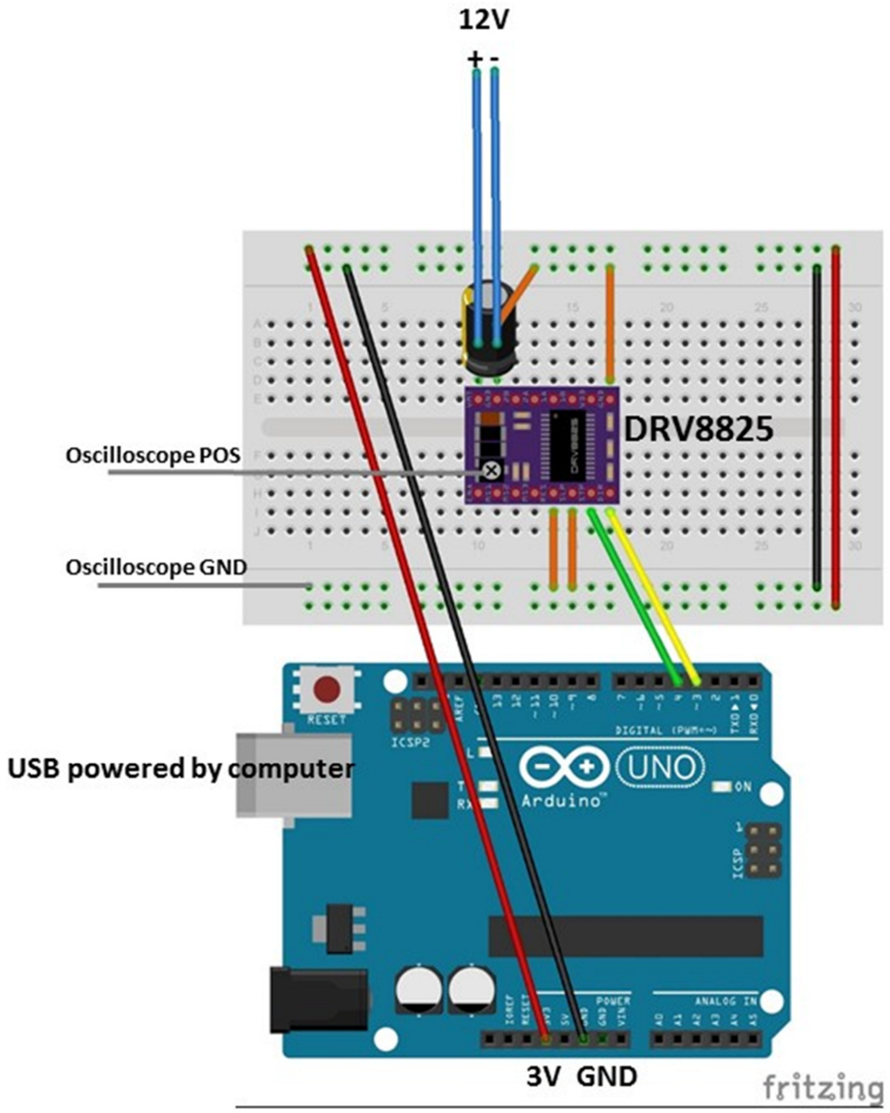


Figure 2. Wiring diagram to enable Vref adjustment of DRV8825 stepper motor drivers

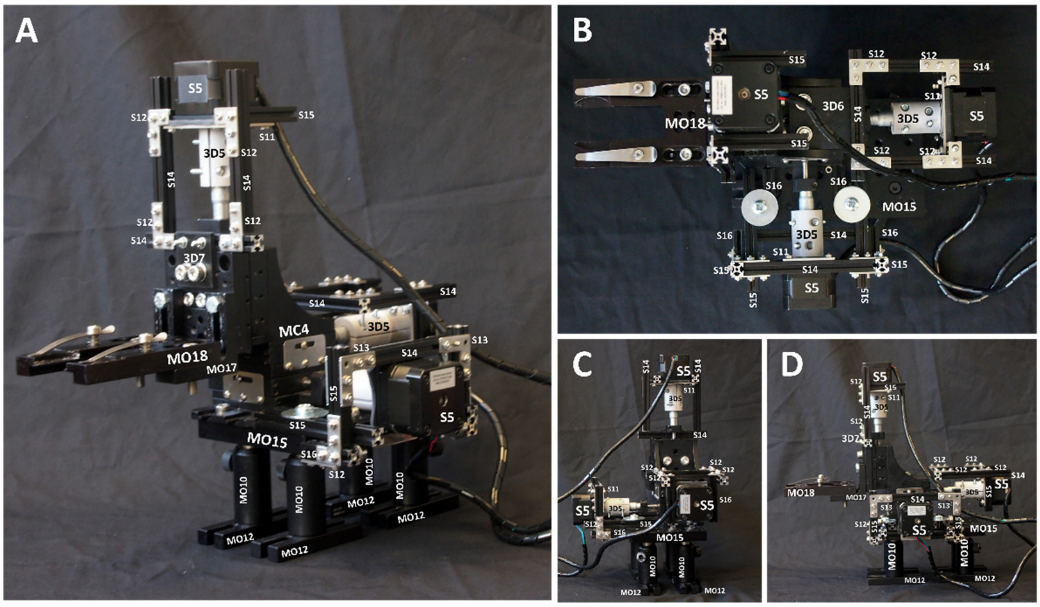


Figure 3. How to assemble the XYZ translation stage (MC4) to enable stepper motor motion (S5) control.

$0 = 10 (Step pulse time, microseconds)
$1 = 25 (Step idle delay, milliseconds)
$2 = 0 (Step pulse invert, mask)
$3 = 0 (Step direction invert, mask)
$4 = 0 (Invert step enable pin, boolean)
$5 = 0 (Invert limit pins, boolean)
$6 = 0 (Invert probe pin, boolean)
$10 = 1 (Status report options, mask)
$11 = 0.010 (Junction deviation, millimeters)
$12 = 0.002 (Arc tolerance, millimeters)
$13 = 0 (Report in inches, boolean)
$20 = 0 (Soft limits enable, boolean)
$21 = 0 (Hard limits enable, boolean)
$22 = 0 (Homing cycle enable, boolean)
$23 = 0 (Homing direction invert, mask)
$24 = 25.000 (Homing locate feed rate, mm/min)
$25 = 500.000 (Homing search seek rate, mm/min)
$26 = 250 (Homing switch debounce delay, milliseconds)
$27 = 1.000 (Homing switch pull-off distance, millimeters)
$30 = 1000 (Maximum spindle speed, RPM)
$31 = 0 (Minimum spindle speed, RPM)
$32 = 0 (Laser-mode enable, boolean)
$100 = 12800.000 (X-axis travel resolution, step/mm)
$101 = 12800.000 (Y-axis travel resolution, step/mm)
$102 = 12800.000 (Z-axis travel resolution, step/mm)
$110 = 30.000 (X-axis maximum rate, mm/min)
$111 = 30.000 (Y-axis maximum rate, mm/min)
$112 = 30.000 (Z-axis maximum rate, mm/min)
$120 = 1.000 (X-axis acceleration, mm/sec^2)
$121 = 1.000 (Y-axis acceleration, mm/sec^2)
$122 = 1.000 (Z-axis acceleration, mm/sec^2)
$130 = 13.000 (X-axis maximum travel, millimeters)
$131 = 13.000 (Y-axis maximum travel, millimeters)
$132 = 13.000 (Z-axis maximum travel, millimeters)

Figure 4. gbrl settings in UGS

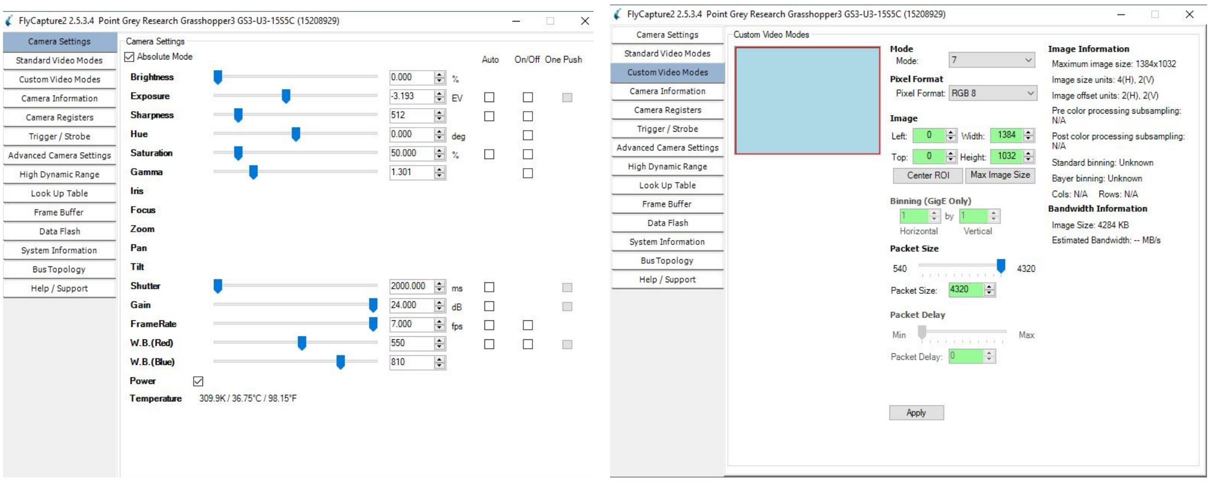


Figure 5. FlyCap settings for CO23 for fluorescent imaging

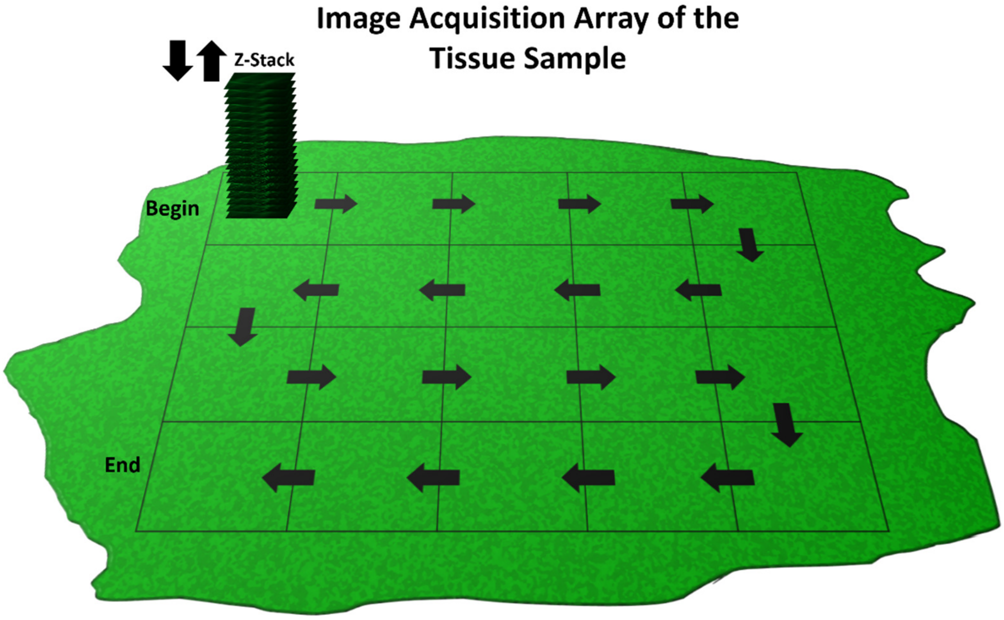


Figure 6. Schematic representation of the automated stage motion control

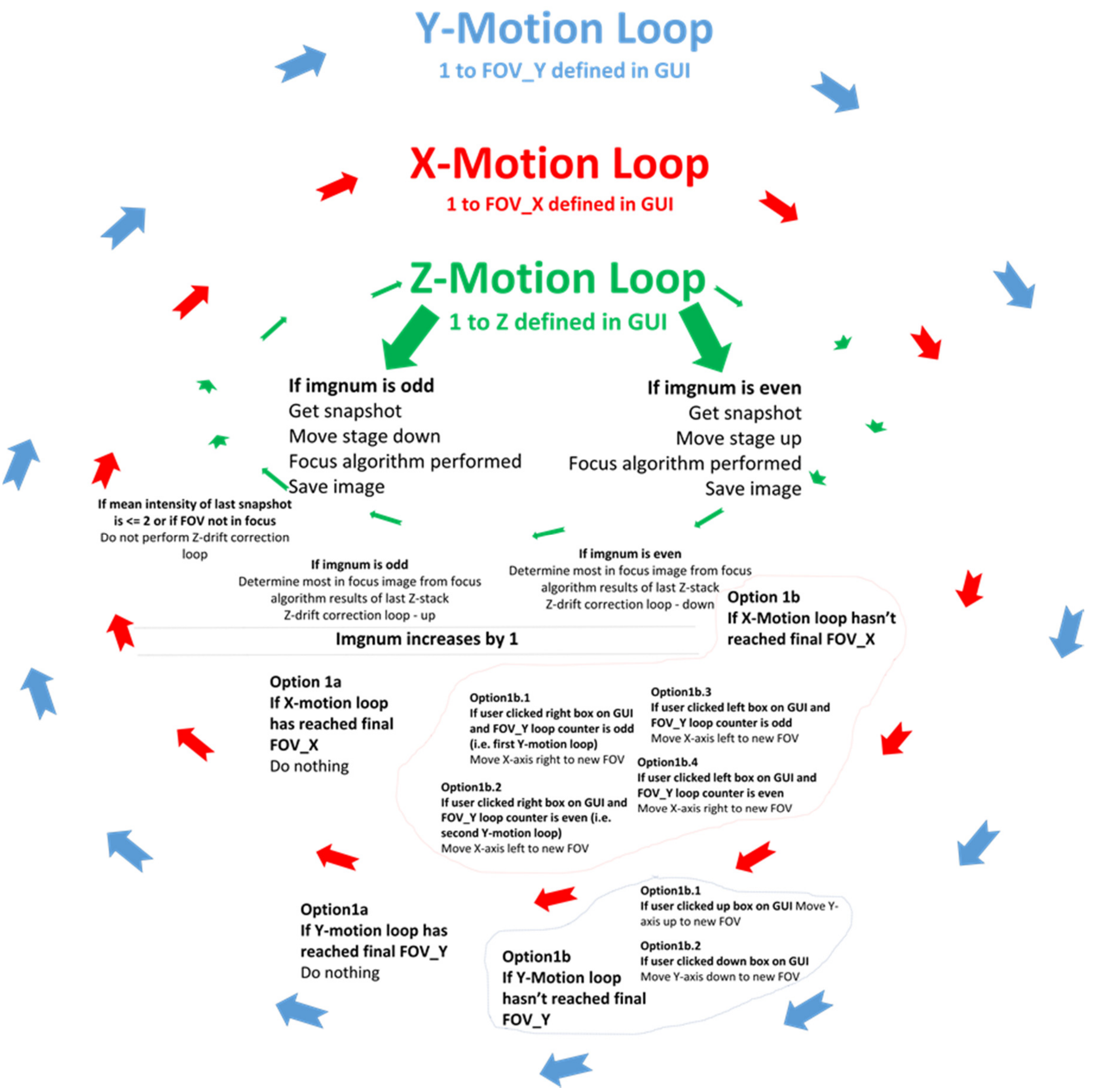


Figure 7. Schematic representation of the automated stepper stage motion control and image acquisition code.
