## Supplementary File 3 for "The Flexiscope: a Low Cost, Flexible, Convertible, and Modular Microscope with Automated Scanning and Micromanipulation": ~WRL0005.tmp

**User Guidelines: Automated**

**Acquisition and Piezoelectric Stage**

**Scanning**

Automated Scanning and Micromanipulation.

Amy Courtney*, Luke Alvey, George O.T. Merces, and Mark Pickering.

School of Medicine, University College Dublin, Ireland.

March 2019

Step1, 2 and 3 should be performed once when you are establishing your system.

Steps 4-6 are performed each time you want to set up a tissue sample for automated scanning.

**Step5: Install Anaconda (Python 3.6), Spinnaker SDK and relevant libraries**

A. Go to <https://repo.continuum.io/archive/> and download relevant anaconda installation file for your operating system (I used Anaconda3-5.2.0-Windows-x86_64.exe)
B. Go to <https://www.ptgrey.com/support/downloads> and download the latest version of Spinnaker SDK (I used: Anaconda3-5.2.0-Windows-x86_64.exe) and the latest Spinnaker for Python library (I used: spinnaker_python-1.20.0.15-cp36-cp36m-win_amd64.whl).
C. Open the anaconda command prompt and install pySerial and OpenCV.
D. Open Spyder (included in the anaconda installation) and open ‘xxxxxx.py’
E. Ensure all libraries are installed by running line xx-xx.
F. Any variables between line xx and xx which include #change this should be altered to suit your parameters. Also go to line 171, 189 and 257.

FOVrightleftWait: Time to wait in seconds when X-stepper moving

FOVdownupWait: Time to wait in seconds when Y-stepper moving

**X/Y scanning and Z-stack acquisition explained**

Figure 7 explains the workflow to acquire Z-stacks at regions of the sample in the X and Y direction. The commands used to control the specimen stage followed a logic in which the image sequence of the tissue sample is considered a two dimensional array. To acquire images of the whole tissue (or a region of interest) the stage must be moved in the X-axis a defined number of times and then moved in the Y-axis once, this cycle can then be repeated until the tissue (or region of interest) has been imaged in its entirety. Each movement in X or Y dimension reveals a new ‘field of view’ (FOV) and a Z-stack is subsequently acquired. The motion control and image acquisition code is explained schematically in figure 8.

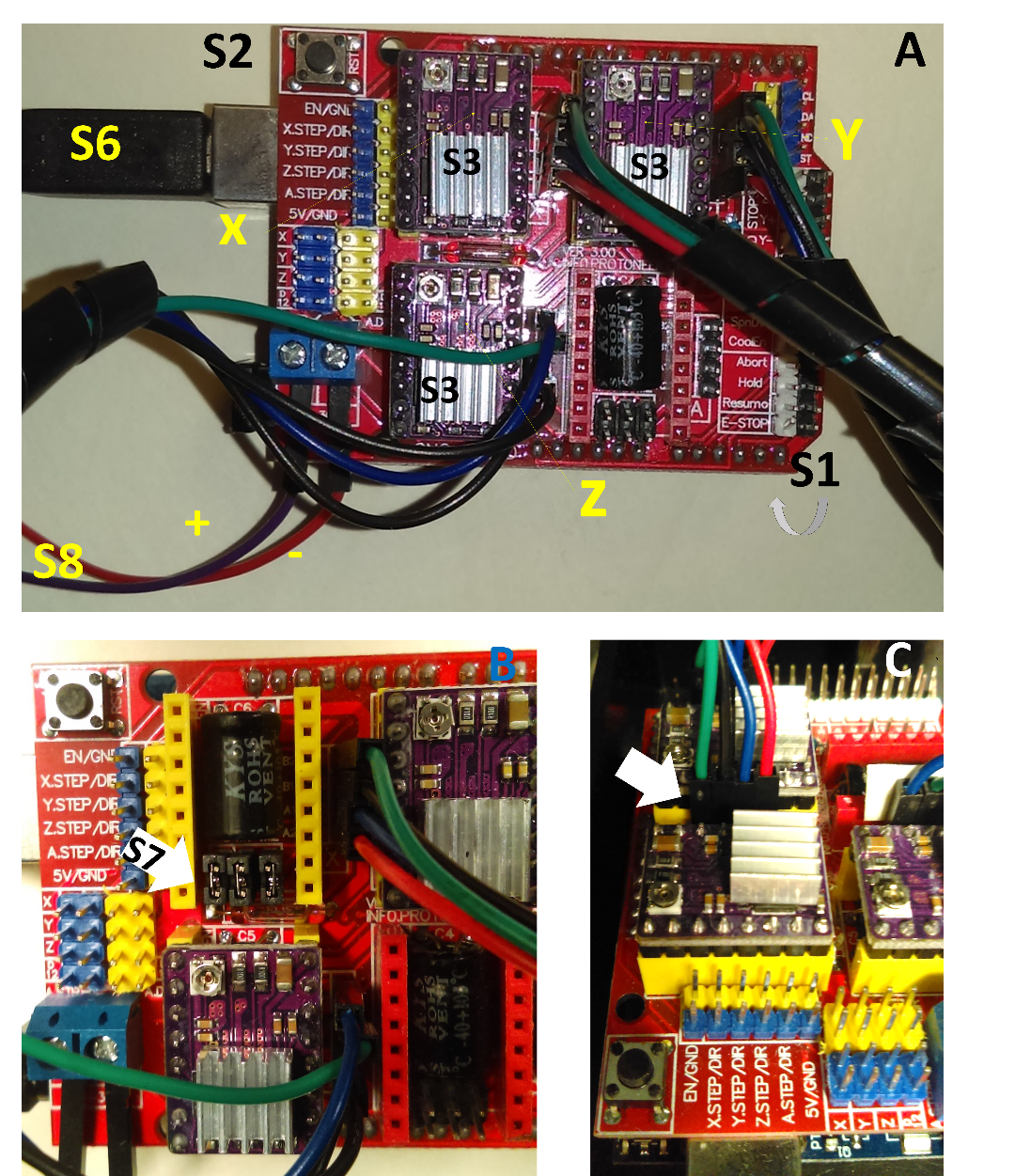

Figure 1. How to set up the Arduino Uno and CNC shield

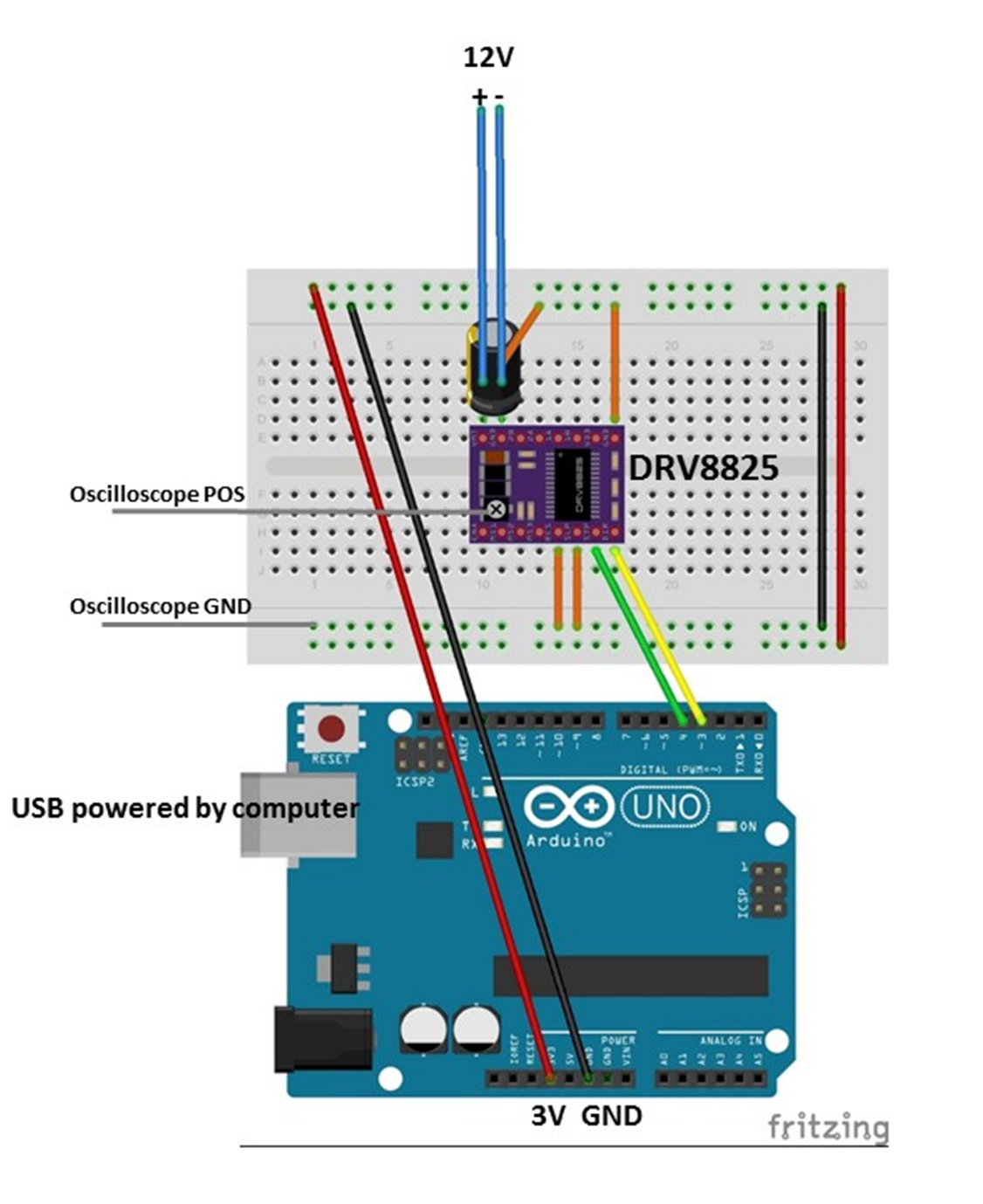

Figure 2. Wiring diagram to enable Vref adjustment of DRV8825 stepper motor drivers

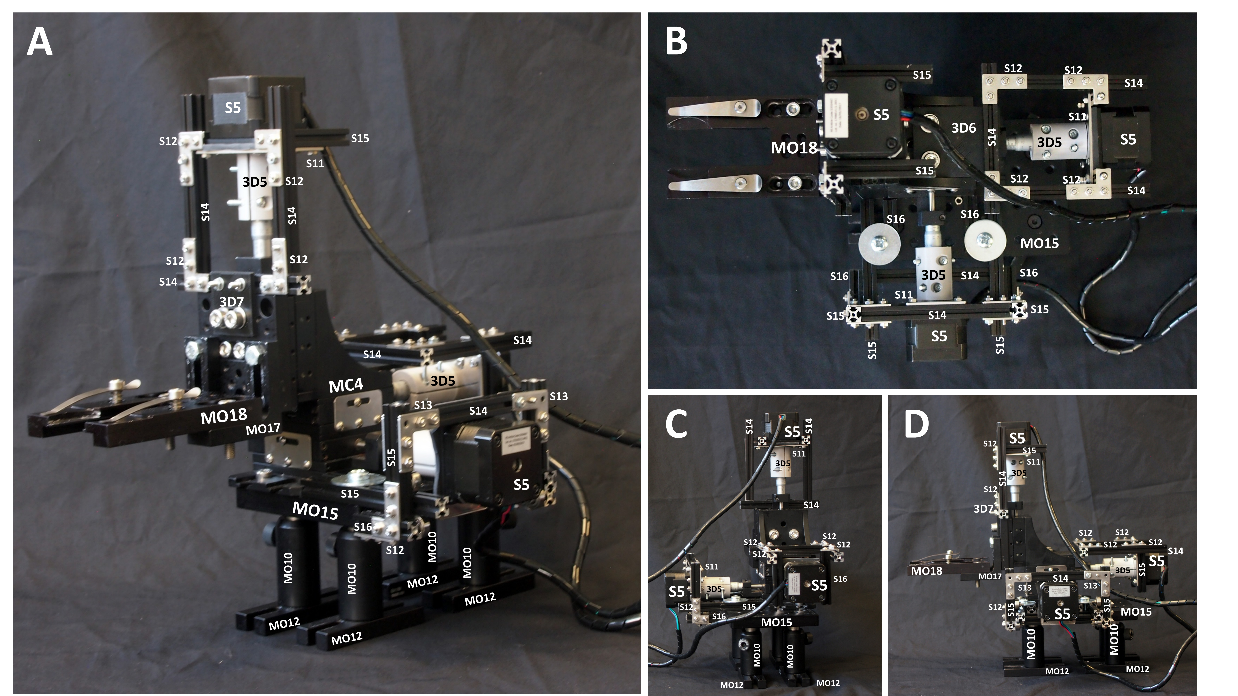

Figure 3. How to assemble the XYZ translation stage (MC4) to enable stepper motor motion (S5) control.

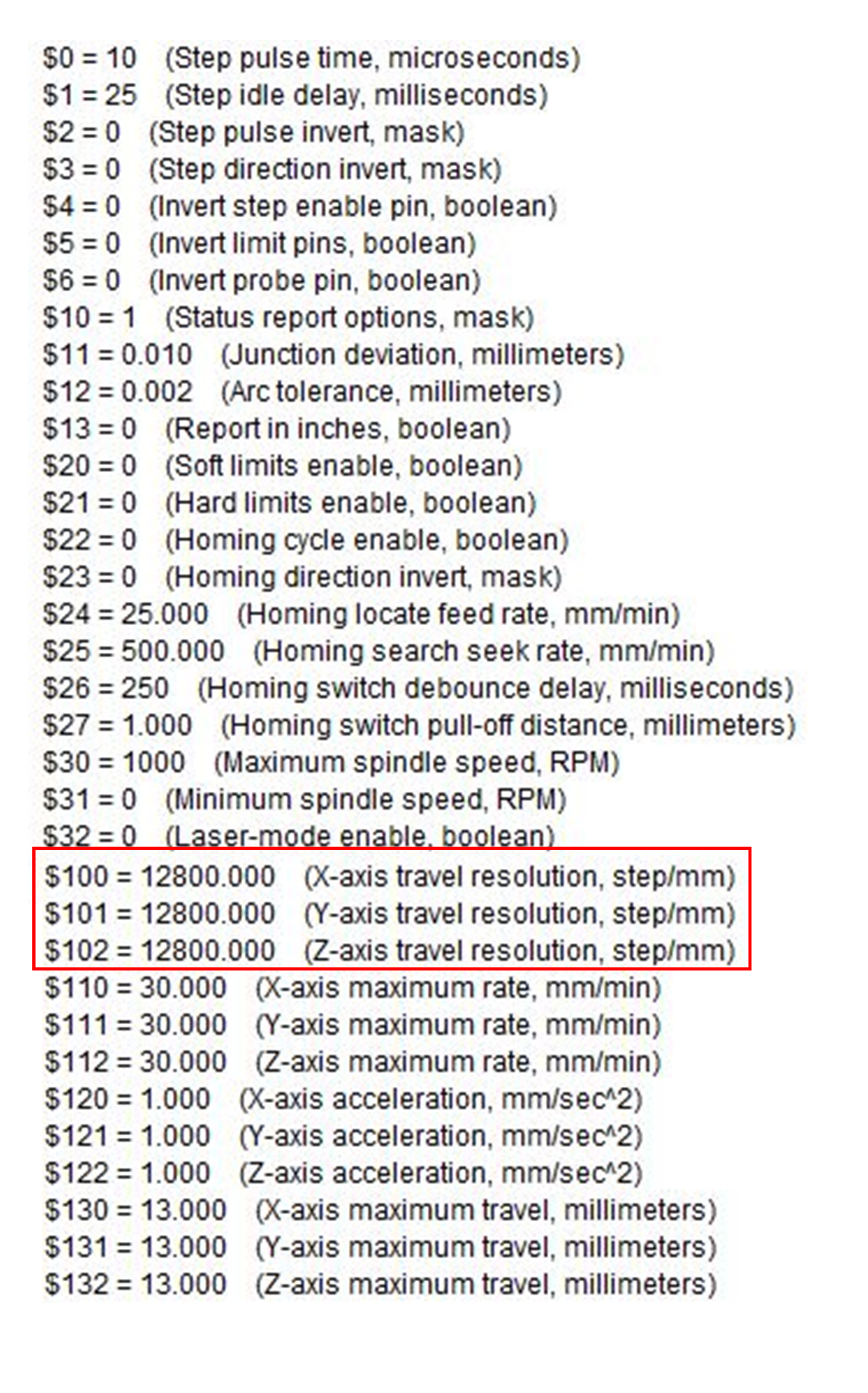

Figure 4. gbrl settings in UGS

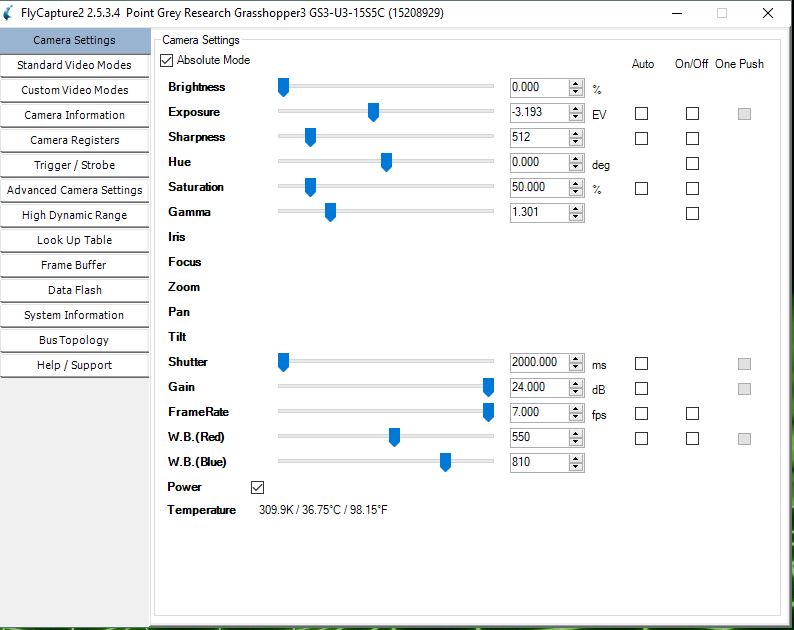

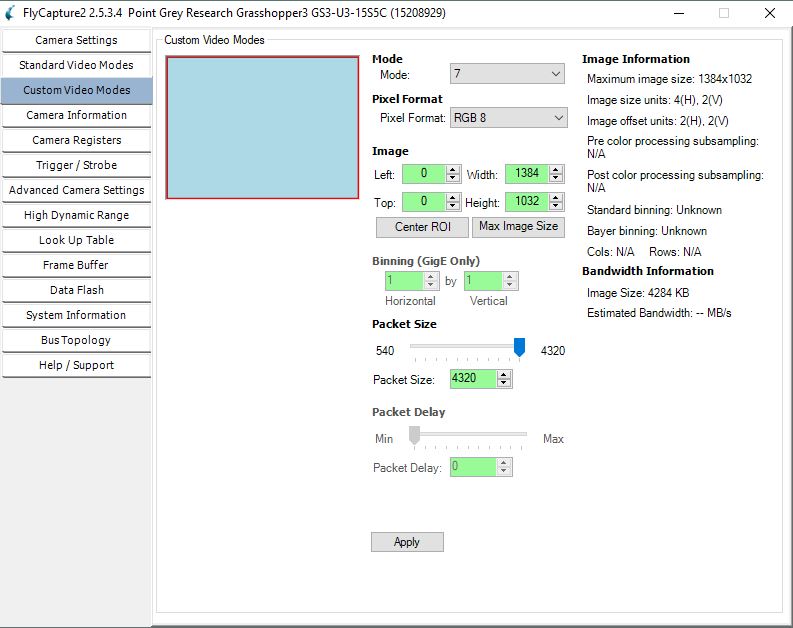

Figure 5. FlyCap settings for CO23 for fluorescent imaging

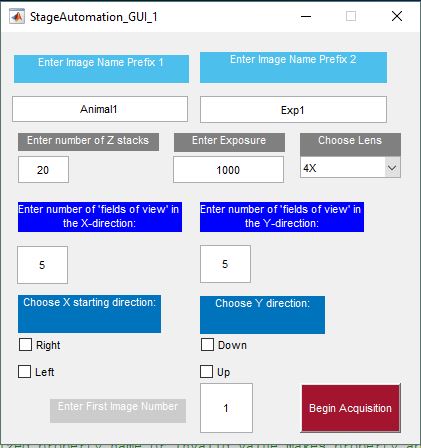

Figure 6. Automated stepper stage motion control GUI in Matlab

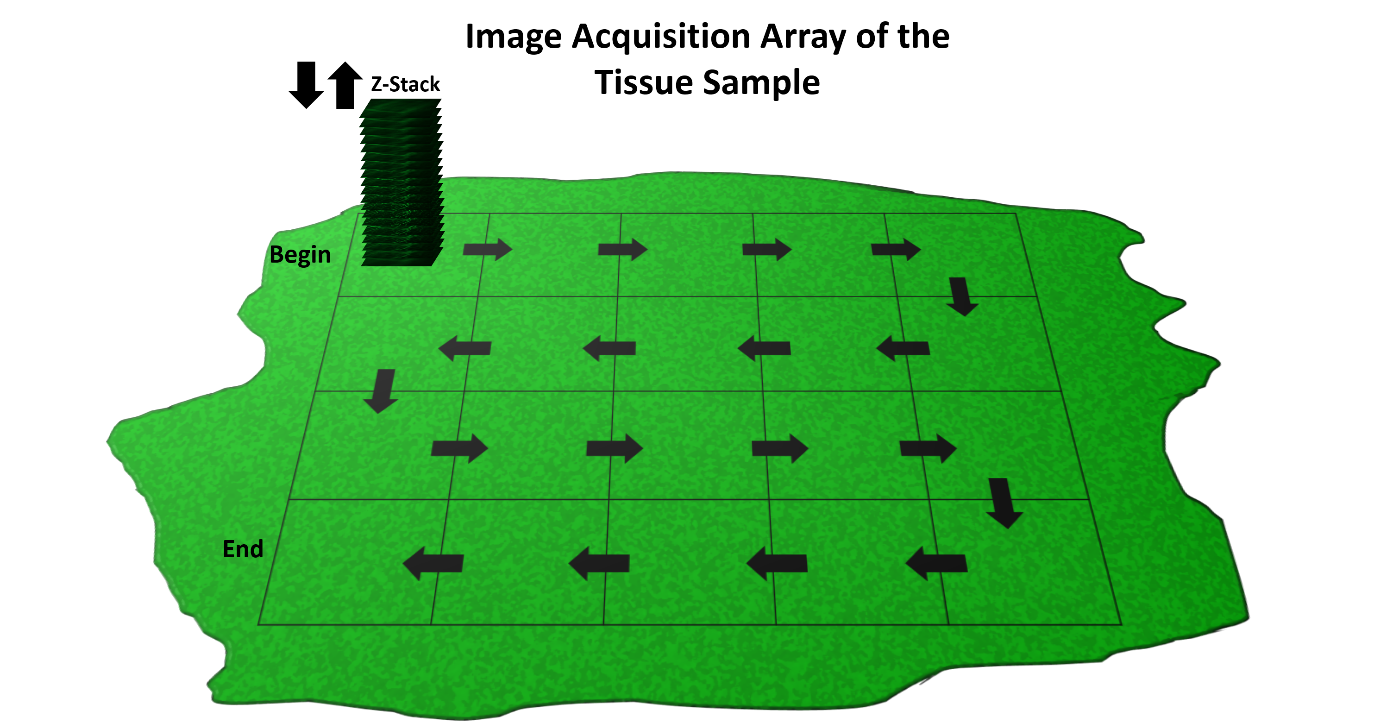

Figure 7. Schematic representation of the automated stage motion control

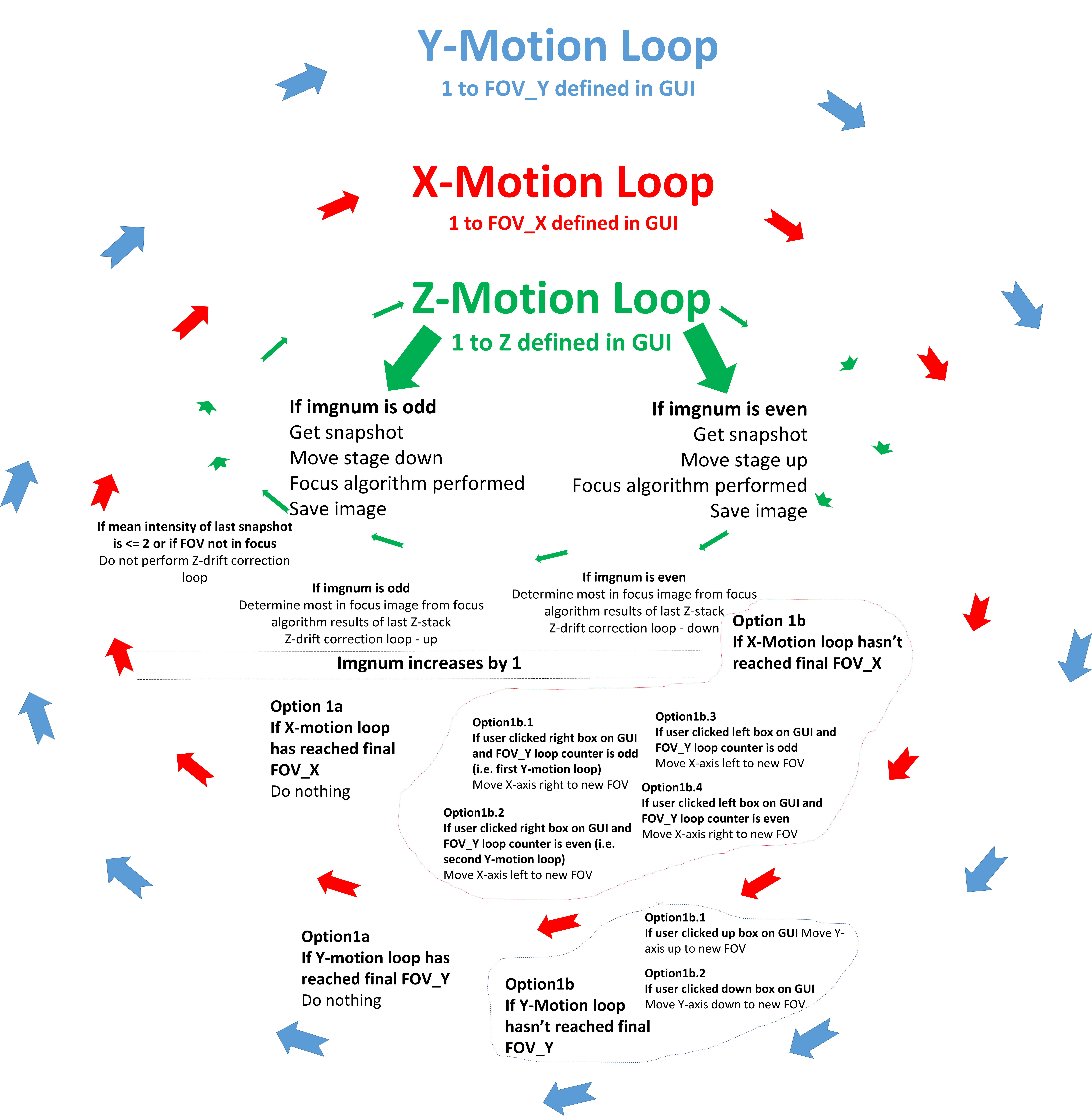

Figure 8. Schematic representation of the automated stepper stage motion control and image acquisition code.
