## Supplementary File 4 for "The Flexiscope: a Low Cost, Flexible, Convertible, and Modular Microscope with Automated Scanning and Micromanipulation": Flexiscope_AssemblyInstructions.pdf

### User Guidelines: Upright Flexiscope Assembly

**October 2019**

This instruction manual describes how to assemble the Flexiscope in the upright configuration.

As described in section 2.1 of the manuscript, the core optical components can be divided into three functional units: the fluorescence illumination module, the infinite space module and the camera module. Here, we will describe how to assemble each module and then combine and mount them on an optical table for an upright configuration. This systems utilises threaded 1” diameter lens tubes.
Each individual component has been allocated a part designation, for use throughout this instruction manual and within the manuscript. Table 1 lists each part and its designation.

#### 1. The Fluorescence Illumination Module

1.1. Begin by fitting each dichroic mounting cube (C07) with their designated dichroics (CO8 and CO9). The dichroics are mounted in the cubes as described on the manufacturer’s website.

1.2. The dichroics are incorporated into CO7 at 45° angles. There is a line mark on the side with the dichroic coating. The LED illumination from CO2 and CO3 should be directed towards this side of the respective filters (CO8 and CO9). See Figure 1.

1.3. Seal the open holes in the cubes with dust caps (CO38).

1.4. Combine the cubes (CO7) using a coupler (CO27). See Figure 2 B.

1.5. Each LED (CO1, CO2 & CO3) is collimated using an aspheric condenser lens (CO6). The lens is placed in CO37 and held in place with a retaining ring using an adjustable spanner wrench (T1).

1.6. The distance from the LED to the lens should be 20mm and was measured with an electronic calibres (Figure 3). Further details are available on the manufacturer's website. LED collimation testing is explained in section 5 of this document.

1.7. Each collimated LED is attached to the dichroic mounting cubes (CO7) using a coupler (CO27).

1.8. The Iris (CO10) is incorporated into the system using CO33 and CO27 (Figure 2 B). Note: this component is optional. We found that the iris caused issues with uneven illumination and it was therefore omitted in later configurations of the Flexiscope.

#### 2. The Infinite Space Module

2.1. Begin by configuring the swappable filter set mounts (CO15). The three CO15s contain an excitation, emission and dichroic filter corresponding to either 630nm (Texas Red, CO12), 530nm (FITC, CO13) or 460nm (BFP, CO14) emission wavelengths. The filters are incorporated into the mount as described on the manufacturer's website.

2.2. The main component in the infinite space module is the filter mount-holding cube (CO16). It is connected to the fluorescence illumination module via the iris/CO27 as seen in Figure 2 B.

2.3. Attach the turning mirror (CO11) to the filter mount-holding cube with the cage cube connector (CO29) as seen in Figure 2 D.

2.4. Attach the lens turret (CO18) to the filter mount-holding cube with CO34 and attach the infinite conjugate objectives (CO18, CO19) to the lens turret as seen in Figure 2 D.

2.5. Place the achromatic doublet lens in CO36 with a retaining ring and adjustable spanner wrench (T1).

2.6. The turning mirror (CO11) is attached to the achromatic doublet lens (CO21) via CO27 as seen in Figure 2 A.

#### 3. The Camera Module

3.1. The distance between the camera sensor and the achromatic doublet lens (CO21) should be 150mm. This is achieved by combining CO31, CO28, CO35, and CO32.

3.2. Ensure that CO31 and CO32 has the M4 mount point orientated downwards (Note: reconfiguring to the inverted configuration means the point at which the achromatic doublet is couplet to the mirror will need to be rotated to enable mounting. Do not change the distance after the achromatic to the camera sensor during reconfiguration).

3.3. Four alignment posts (CO30) are also attached as seen in Figure 2 A & C.

3.4. The camera is then attached to CO32 with a C-mount adapter (CO24). We used a 3D printed component to align the camera (3D4).

### 4. Mounting and Motion Control Components

4.1. Two optical posts (MO4) are attached to the dichroic mounting cubes (CO7) and combined with post holders (MO11) and post mounts (MO13). See Figure 2 D.

4.2. An optical post (MO5) is attached to the cage plate (CO32). A post holder (MO11) and post mount (MO13) is also used as seen in Figure 2 A.

4.3. The entire scope is then attached to the lab jack (MC1) via the post mounts using M6 bolts as seen in Figure 2 D.

4.4. The piezoelectric actuators (MC2) are incorporated into the XYZ translational stage (MC4) as seen in Figure 2 B.

4.5. Specimen holder configuration on XYZ stage: base plate (MO16), two right angle brackets (MO17) and a microscopy slide holder (MO18). See Figure 2 D.

4.6. The XYZ stage is mounted to a small breadboard (MO15).

4.7. Optical posts (MO10), post holders (MO8) and post mounts (MO13 or MO12) mounted on the breadboard/stage.

4.8. The breadboard/stage and the scope/lab jack can be aligned to each other and then mounted directly to the main breadboard (MO19) using M6 bolts.

### 5. Calibration

**5.1. Testing LED collimation:**

- In order to achieve diffuse flat illumination on your sample LED collimation may need to be adjusted. Using the fluorescent slides (CA2) show a live preview of the camera on the point grey FlyCapture software. A live histogram can be opened in the software. Adjust CO37 until the histogram is flat as seen in Figure 4.

- For the 565nm LED use the red slide, for the 470nm LED use the green slide and for the 395nm LED use the blue slide.

**5.2. Determining pixel size:**

- Determining pixel size: Acquire an image of a calibration slide and measure length of defined distance and convert from pixels to µm using imageJ as seen in Figure 5.

#### Table 1. Flexiscope Component List with Designators and a Cost Description.

| Component | Part Designator | Component Description | Product Code | Supplier | Quantity | Cost per unit (€) | Total cost (€) |
|---|---|---|---|---|---|---|---|
| ***Core Optical Components*** | | | | | | | |
| **Fluorescence Illumination Module** | CO1 | Lime (565 nm) Mounted LED | M565L3 | ThorLabs | 1 | 186.10 | 186.10 |
| | CO2 | Blue (470 nm) Mounted LED | M470L3 | ThorLabs | 1 | 236.38 | 236.38 |
| | CO3 | UV (395 nm) Mounted LED | M395L4 | ThorLabs | 1 | 249.61 | 249.61 |
| | CO4 | LED Driver | LEDD1B | ThorLabs | 3 | 258.43 | 775.29 |
| | CO5 | Power Supply for LED and Piezo T-Cubes | TPS008 | ThorLabs | 1 | 158.76 | 158.76 |
| | CO6 | Aspheric Condenser Lens (for LED collimation) | ACL2520U-DG6-A | ThorLabs | 3 | 24.43 | 73.29 |
| | CO7 | Dichroic mounting cube | CM1-DCH/M | ThorLabs | 2 | 139.36 | 278.72 |
| | CO8 | GFP Dichroic Filter (Refl. Band = 452-490 nm, Trans. Band = 505-800 nm) | MD498 | ThorLabs | 1 | 188.87 | 188.87 |
| | CO9 | BFP Dichroic Filter (Refl. Band = 360-407 nm, Trans. Band = 425-575 nm ) | MD416 | ThorLabs | 1 | 188.87 | 188.87 |
| | CO10 | Iris Diaphragm | SM1D12D | ThorLabs | 1 | 56.71 | 56.71 |
| **Infinite Space Module** | CO11 | Protected Silver Tuning Mirror | CCM1-P01/M | ThorLabs | 1 | 144.90 | 144.90 |
| | CO12 | Texas Red Filter Set (Excitation, Emission, and Dichroic) | MDF-TXRED | ThorLabs | 1 | 551.25 | 551.25 |
| | CO13 | FITC Filter Set (Excitation, Emission, and Dichroic) | MDF-FITC | ThorLabs | 1 | 551.25 | 551.25 |
| | CO14 | BFP Texas Red Filter Set (Excitation, Emission, and Dichroic) | MDF-BFP | ThorLabs | 1 | 551.25 | 551.25 |
| | CO15 | Swappable Filter Set Mount | DFMT1 | ThorLabs | 3 | 177.28 | 531.84 |
| | CO16 | Filter Mount Holding Cube | DFMB/M | ThorLabs | 1 | 90.85 | 90.85 |
| | CO17 | Filter Cube Blank Top Plate | DFM1C | ThorLabs | 1 | 32.40 | 32.40 |
| | CO18 | Objective Lens Turret | OT1 | ThorLabs | 1 | 262.53 | 262.53 |
| | CO19 | 4X Olympus Plan Achromat Objective (NA=0.10, WD=18.5 mm) | RMS4X | ThorLabs | 1 | 167.07 | 167.07 |
| | CO20 | 20X Olympus Water Immersion Objective (NA=0.5, WD=3.5 mm) | UMPLFLN20XW | Masons | 1 | 1,176.68 | 1,176.68 |
| **Camera Module** | CO21 | Tube lens: Achromatic Doublet (f=150.0 mm) | AC254-150-A | ThorLabs | 1 | 62.56 | 62.56 |
| | CO22 | USB 3.0 Monochrome Camera (Flea®3 ½" FL3-U3-13Y3M-C) | 86-767 | Edmund Optics | 1 | 783.75 | 783.75 |
| | CO23 | USB 3.0 Color Camera (Grasshopper GS3-U3-15S5C-C 2/3" ) | 33-533 | Edmund Optics | 1 | 1,435.09 | 1,435.09 |
| | CO24 | 1" Lens Tube to C-Mount Adapter (Male to Male) | SM1A39 | ThorLabs | 1 | 18.60 | 18.60 |
| | CO25 | USB 3.0 Locking Cable | 86-770 | Edmund Optics | 1 | 23.75 | 23.75 |
| | CO26 | Infrared LED array | N/A | Salvaged from CCTV Camera | 1 | N/A | N/A |
| **Coupling and Alignment** | CO27 | Male to Male Tube Coupler (0.5") | SM1T2 | ThorLabs | 6 | 17.55 | 105.30 |
| | CO28 | Male to Male Tube Coupler (2") | SM1T20 | ThorLabs | 1 | 18.74 | 18.74 |
| | CO29 | Cage Cube Connector | C4W-CC | ThorLabs | 1 | 43.57 | 43.57 |
| | CO30 | Alignment Posts (Diameter = 6 mm, L = 75 mm) | MS3R/M | ThorLabs | 4 | 7.10 | 28.40 |
| | CO31 | 1" Threaded Cage Plate (0.50" Thick) | CP02T/M | ThorLabs | 1 | 17.90 | 17.90 |
| | CO32 | 1" Threaded Cage Plate (0.35" Thick) | CP02/M | ThorLabs | 2 | 14.11 | 28.22 |
| | CO33 | 1" Lens Tube, 0.30" Thread Depth | SM1L03 | ThorLabs | 4 | 10.72 | 42.88 |
| | CO34 | 1" Lens Tube, 0.5" Thread Depth | SM1L05 | ThorLabs | 3 | 11.33 | 33.99 |
| | CO35 | 1" Lens Tube, 3" Thread Depth | SM1L30 | ThorLabs | 1 | 21.95 | 21.95 |
| | CO36 | 1" Lens Tube, 0.31" Travel Range (Adjustable) | SM1V05 | ThorLabs | 2 | 26.11 | 52.22 |
| | CO37 | 1" Lens Tube, 0.81" Travel Range (Adjustable) | SM1V10 | ThorLabs | 1 | 29.34 | 29.34 |
| | CO38 | Plastic Dust Cap for 1" Lens Tubes | SM1EC2 | ThorLabs | 10 | 1.76 | 17.64 |
| | CO39 | 1" Lens Tube, 1.5" Thread Depth | SM1L15 | ThorLabs | 1 | 14.27 | 14.27 |
| ***Other components*** | | | | | | | |
| **Motion Control Components** | MC1 | Lab Jack | L490/M | ThorLabs | 1 | 527.40 | 527.40 |
| | MC2 | Piezoelectric Actuator (13 mm Travel) | PIA13 | ThorLabs | 4 | 440.12 | 1,760.48 |
| | MC3 | T-Cube: Piezoelectric Actuator Controller | TIM101 | ThorLabs | 1 | 837.90 | 837.90 |
| | MC4 | 13 mm XYZ Translation Stage | MT3/M | ThorLabs | 1 | 798.21 | 798.21 |
| | MC5 | 13 mm Translation Stage Plate in One Dimension | MT1A/M | ThorLabs | 1 | 356.00 | 356.00 |
| | MC6 | 25 mm XYZ Translation Stage | PT3A/M | ThorLabs | 1 | 954.00 | 954.00 |
| **Mounting Components** | MO1 | Optical Post (Ø12.7 mm, L = 30 mm) | TR30/M-P5 | ThorLabs | 5 | 3.84 | 19.20 |
| | MO2 | Optical Post (Ø12.7 mm, L = 20 mm) | TR20/M-P5 | ThorLabs | 5 | 3.76 | 18.82 |
| | MO3 | Optical Post (Ø12.7 mm, L = 50 mm) | TR50/M-P5 | ThorLabs | 5 | 4.12 | 20.60 |
| | MO4 | Optical Post (Ø12.7 mm, L = 75 mm) | TR75/M-P5 | ThorLabs | 5 | 4.39 | 21.95 |
| | MO5 | Optical Post (Ø12.7 mm, L = 100 mm) | TR100/M-P5 | ThorLabs | 5 | 4.50 | 22.52 |
| | MO6 | Optical Construction Post for TR75C/M (Ø12.7 mm) | TR75T/M | ThorLabs | 1 | 14.90 | 14.90 |
| | MO7 | Optical Construction Post- Variable Angle (Ø12.7 mm) | TR75C/M | ThorLabs | 1 | 14.90 | 14.90 |
| | MO8 | Post Holders (Ø12.7 mm, L=30 mm) | PH30/M-P5 | ThorLabs | 5 | 6.33 | 31.64 |
| | MO9 | Post Holders (Ø12.7 mm, L=20 mm) | PH20/M-P5 | ThorLabs | 5 | 6.20 | 31.01 |
| | MO10 | Post Holders (Ø12.7 mm, L=50 mm) | PH50/M-P5 | ThorLabs | 5 | 6.79 | 33.96 |
| | MO11 | Post Holder (Ø12.7 mm, L=75 mm) | PH75/M-P5 | ThorLabs | 5 | 7.05 | 35.25 |
| | MO12 | Mounting Base (25 mm x 75 mm x 10 mm) | BA1/M-P5 | ThorLabs | 5 | 4.45 | 22.23 |
| | MO13 | Mounting Base (25 mm x 58 mm x 10 mm) | BA1S/M-P5 | ThorLabs | 5 | 3.99 | 19.94 |
| | MO14 | Mounting Base (50 mm x 75 mm x 10 mm) | BA2/M-P5 | ThorLabs | 5 | 5.80 | 28.98 |
| | MO15 | Breadbord (100 mm x 150 mm x 12.7 mm) | MB1015/M | ThorLabs | 1 | 36.90 | 36.90 |
| | MO16 | Base Plate for Z-Axis Stage Mounting (65 mm x 65 mm x 10 mm) | UBP2/M | ThorLabs | 1 | 30.87 | 30.87 |
| | MO17 | Right-Angle Bracket | AB90C/M | ThorLabs | 2 | 22.67 | 45.34 |
| | MO18 | Microscopy Slide Holder | MAX3SLH | ThorLabs | 1 | 105.69 | 105.69 |
| | MO19 | Composite Core Optical Breadboard (600 x 600 x 28 mm) | M-TD-22 | Newport | 1 | 488.70 | 488.70 |
| | MO20 | Sorbothane Feet (Ø27.0 mm) | AV4/M | ThorLabs | 4 | 4.69 | 18.74 |
| **Blackout Enclosure** | BE1 | Black Hardboard (610 mm x 610 mm, 5 mm Thick) | TB4 | ThorLabs | 3 | 18.81 | 56.43 |
| | BE2 | Aluminium Extrusion (25 x 25mm, L = 600 mm) | XE25L600/M | ThorLabs | 8 | 25.52 | 204.16 |
| | BE3 | Vertical Blackout Blind (600 mm x 1700 mm) | VBB060 | ThorLabs | 2 | 88.11 | 176.22 |
| | BE4 | Black Masking Tape (50mm x 55m) | T137-2.0 | ThorLabs | 1 | 13.86 | 13.86 |
| **Head Stage Mounting** | HS1 | Male M6 to Female M4 Coupler | AS4M6M | Thorlabs | 2 | 3.81 | 7.62 |
| | HS2 | Male M4 to Female M3 Coupler | MSA4/M | Thorlabs | 2 | 4.17 | 8.34 |
| | HS3 | 45° Brackets | 100281 | MakerBeam | 12 | 0.58 | 6.95 |
| | HS4 | Aluminum Extrusion (10x10mm, L= 100mm) | 100078 | MakerBeam | 16 | 0.58 | 12.25 |
| ***Additional configuration parts*** | | | | | | | |
| **3D printed Components** | 3D1 | Swappable Filter Set Mount Holder | | | 3 | N/A | N/A |
| | 3D2 | Lens Tube - Part1 | | | 1 | N/A | N/A |
| | 3D3 | Lens Tube - Part2 | | | 1 | N/A | N/A |
| | 3D4 | Camera Alignment | *stl files available in supplementary file 5* | | 1 | N/A | N/A |
| | 3D5 | Coupler: Actuator to Stepper | | | 3 | N/A | N/A |
| | 3D6 | Y-axis Mounting | | | 1 | N/A | N/A |
| | 3D7 | Z-axis Mounting | | | 1 | N/A | N/A |
| **Stepper Motor XYZ-Stage** | S1 | Arduino Uno | A000066 | Farnell | 1 | 17.15 | 17.15 |
| | S2 | CNC Shield V3.0 | N/A | gearbest.com | 1 | 4.28 | 4.28 |
| | S3 | DRV8825 Motor Drivers | RB-Pol-272 | RobotShop | 5 | 1.44 | 7.19 |
| | S4 | Jumpers | 791-6454 | Radionics | 20 | 0.23 | 4.62 |
| | S5 | NEMA17 Stepper Motor 1.8°, 0.22nm, 2.8 V, 1.33 A, 4 Wire | 535-0467 | Radionics | 3 | 24.09 | 72.27 |
| | S6 | Male USB A to Male USB B | 529-8274 | Radionics | 1 | 6.21 | 6.21 |
| | S7 | Female Shorting Link | 251-8682 | Radionics | 10 | 0.03 | 0.27 |
| | S8 | 12V Power Supply | 148-963 | Radionics | 1 | 12.89 | 12.89 |
| | S9 | Spiral Binding | 446-172 | Radionics | 1 | 7.21 | 7.21 |
| | S10 | Bearings | 100438 | MakerBeam | 10 | 1.40 | 14.00 |
| | S11 | NEMA17 Stepper Bracket | 100225 | MakerBeam | 3 | 3.50 | 10.50 |
| | S12 | 90° Brackets | 100304 | MakerBeam | 12 | 0.58 | 6.95 |
| | S13 | Right Angle Brackets | 100326 | MakerBeam | 12 | 0.58 | 6.95 |
| | S14 | 100mm Anodised MakerBeam | 100078 | MakerBeam | 16 | 0.77 | 12.25 |
| | S15 | 60mm Anodised MakerBeam | 100056 | MakerBeam | 8 | 0.38 | 3.00 |
| | S16 | 40mm Anodised MakerBeam | 100056 | MakerBeam | 8 | 0.38 | 3.00 |
| **Calibration** | CA1 | 1951 USAF Resolution Test Target | R3L1S4P | ThorLabs | 1 | 194.82 | 194.82 |
| | CA2 | Fluorescent Calibration Microscope Slides | FSK5 | ThorLabs | 1 | 16.85 | 16.85 |
| **Tools** | T1 | Adjustable Spanner Wrench | SPW801 | ThorLabs | 1 | 91.75 | 91.75 |

**Figure 1. Schematic representation of Flexiscope with a clear view of the fluorescent illumination module.**

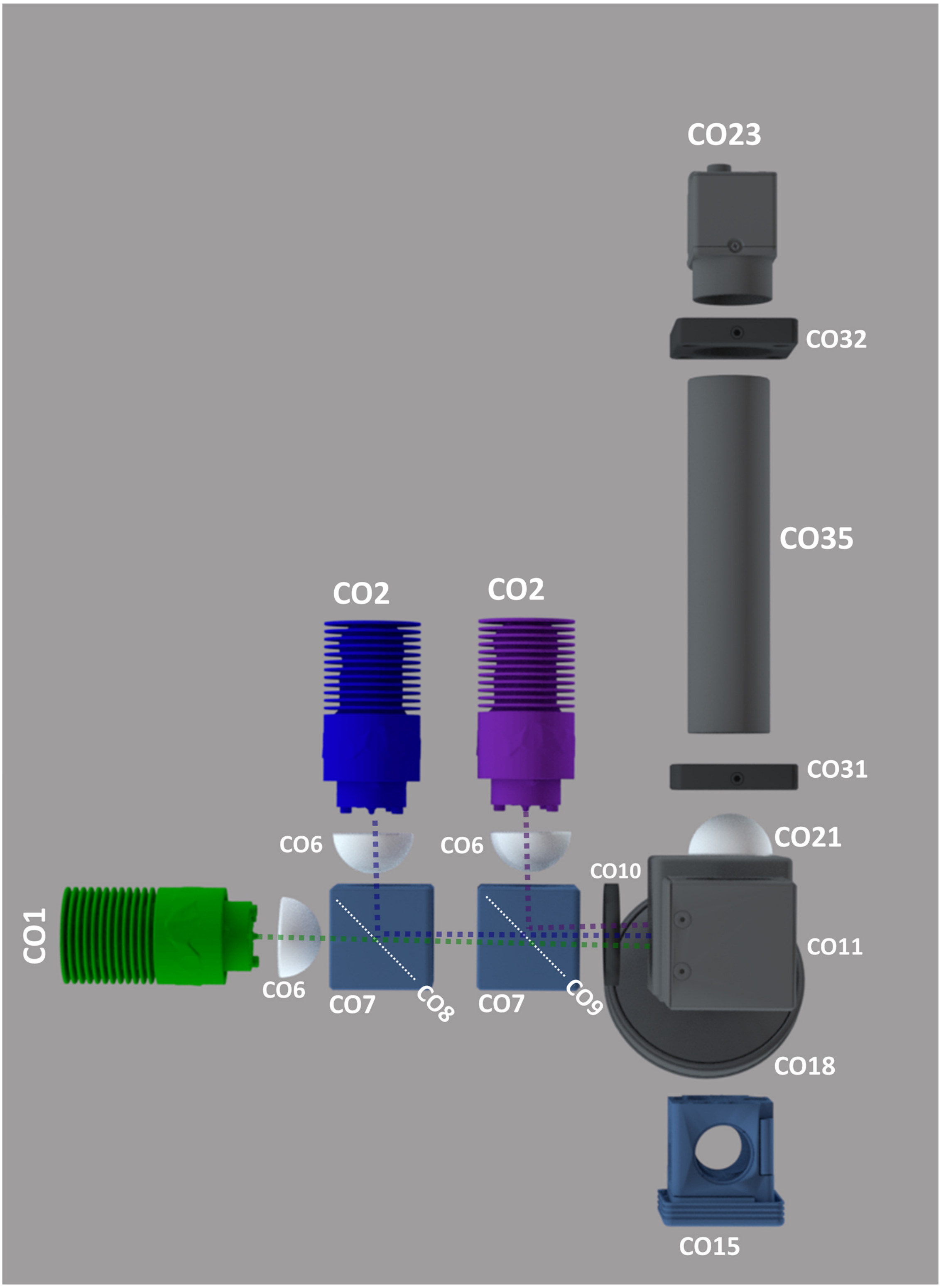

### Figure 2. Photographs of the Flexiscope in the upright configuration with part designators labelled

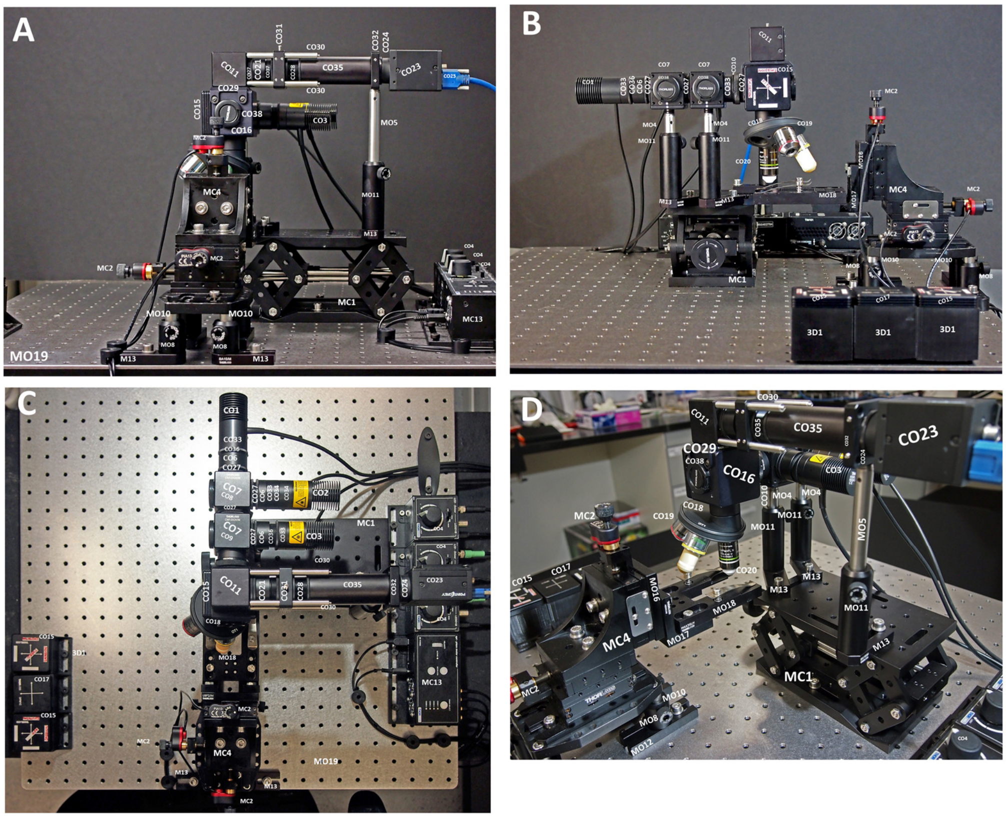

**Figure 3. Mounting aspheric condenser lens for LED collimation.**

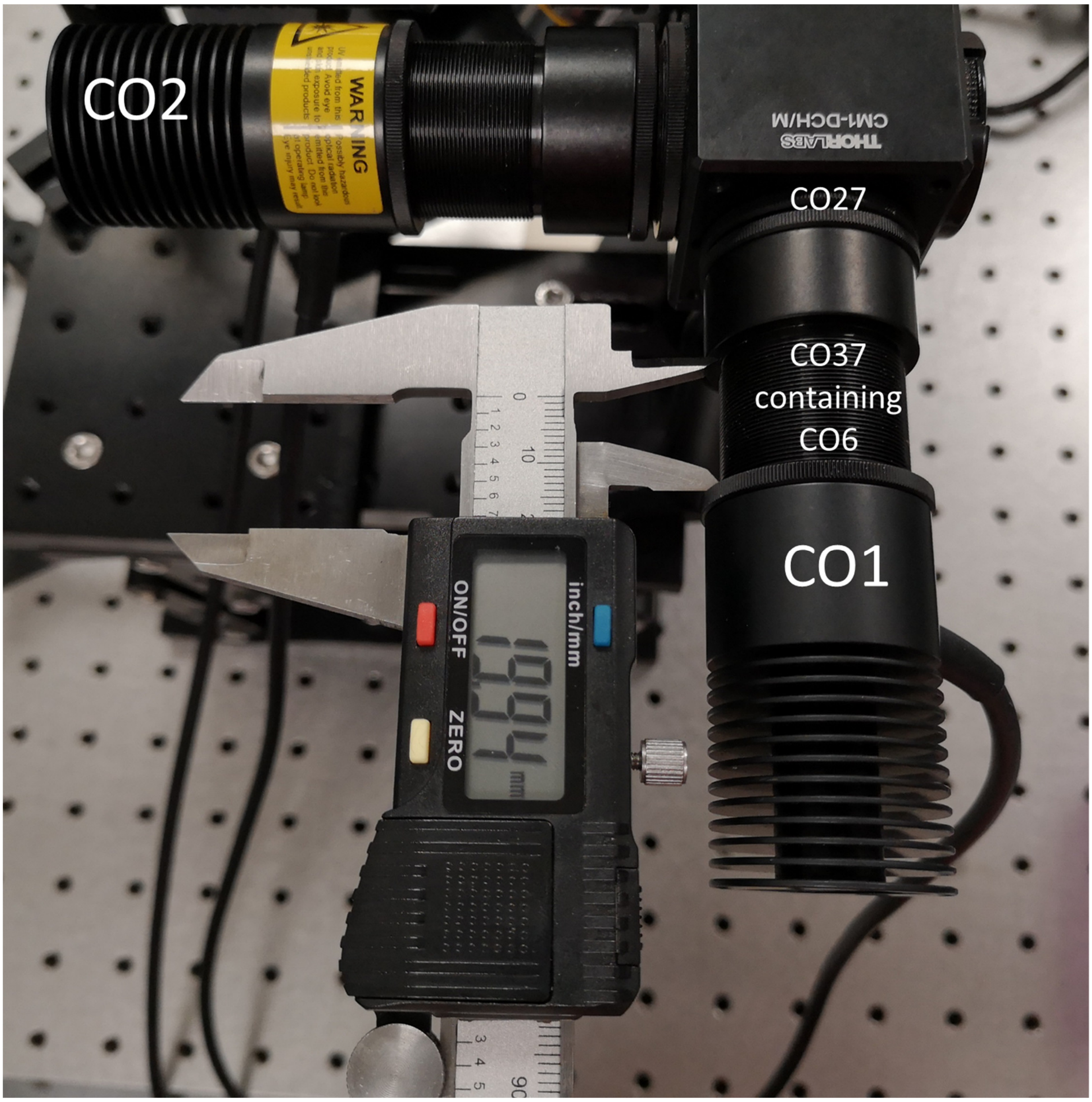

### Figure 4. Testing the LED collimation

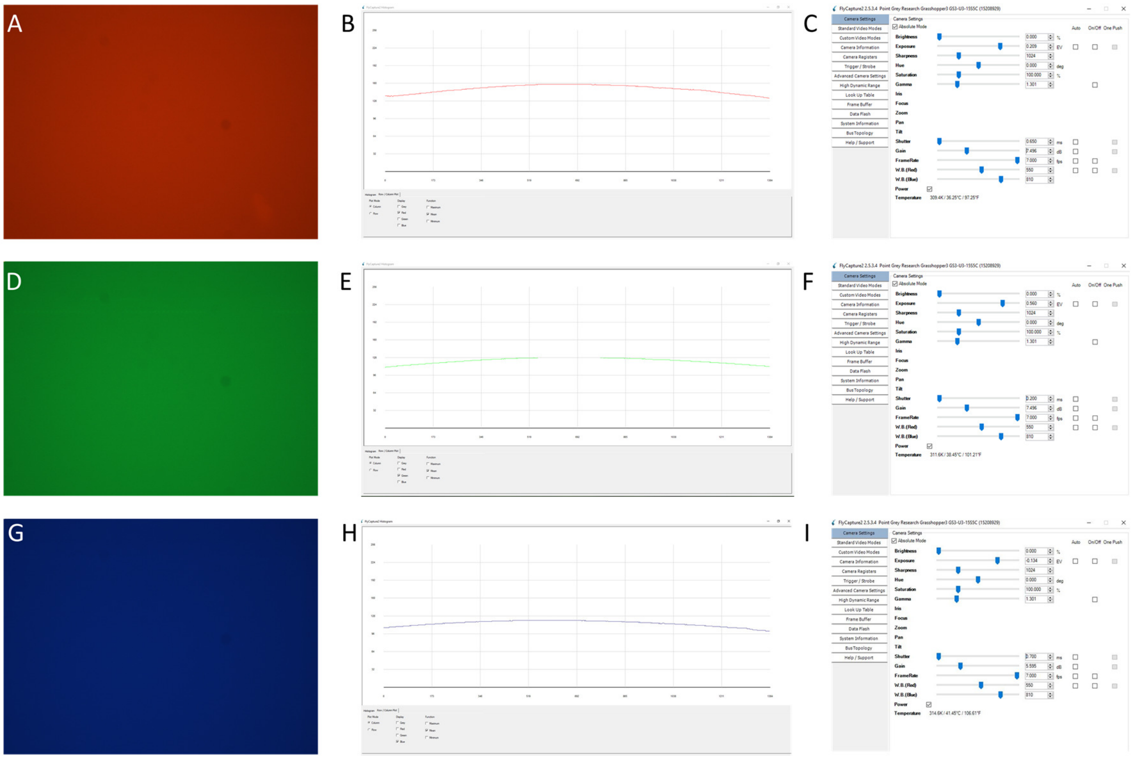

**Figure 5. Determining the pixel size in ImageJ.**

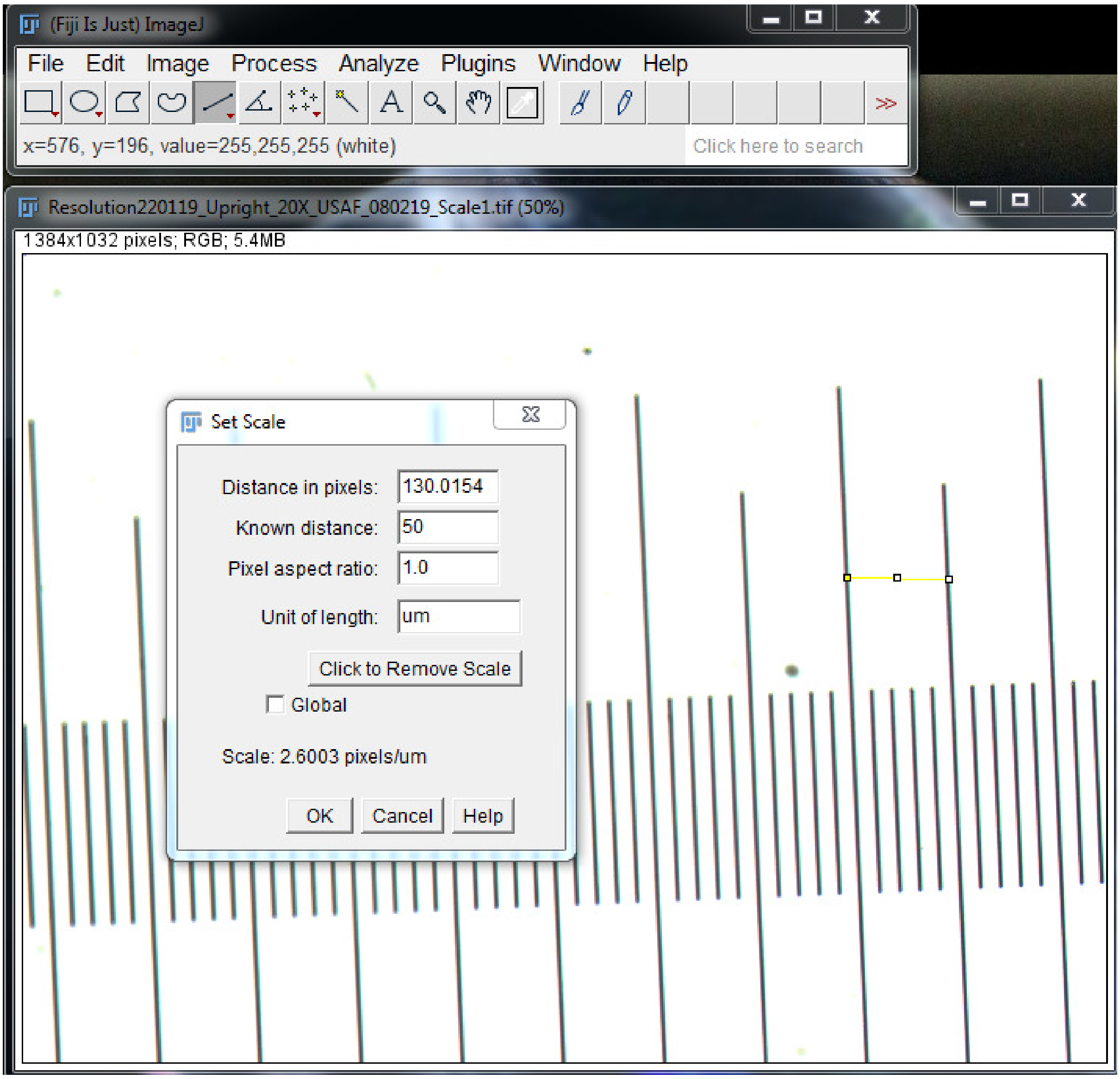
