## Supplementary File 5 for "The Flexiscope: a Low Cost, Flexible, Convertible, and Modular Microscope with Automated Scanning and Micromanipulation": ReadMe.txt

This folder contains 7 stl files. The designators (3D1, 3D2 etc.) are described in table 1.

stl files:

3D1_ SwappableFilterSetMountHolder.stl

3D2_LensTubePart1.stl

3D3_LensTubePart2.stl

3D4_CameraAlignment.stl

3D5_Coupler_stepper(5mm)toActuator(13mm)_Height37_M3_ExDiam25_Gap1.stl

3D6_Y-axisMounting.stl

3D7_Z-axisMounting.stl
